# Nonmodular oscillator and switch based on RNA decay drive regeneration of multimodal gene expression

**DOI:** 10.1101/2022.01.12.475956

**Authors:** Benjamin Nordick, Polly Y. Yu, Guangyuan Liao, Tian Hong

## Abstract

Periodic gene expression dynamics are key to cell and organism physiology. Studies of oscillatory expression have focused on networks with intuitive regulatory negative feedback loops, leaving unknown whether other common biochemical reactions can produce oscillations. Oscillation and noise have been proposed to support mammalian progenitor cells’ capacity to restore heterogenous, multimodal expression from extreme subpopulations, but underlying networks and specific roles of noise remained elusive. We use mass-action-based models to show that regulated RNA degradation involving as few as two RNA species—applicable to nearly half of human protein-coding genes—can generate sustained oscillations without imposed feedback. Diverging oscillation periods synergize with noise to robustly restore cell populations’ bimodal expression. The global bifurcation organizing this divergence relies on an oscillator and bistable switch which cannot be decomposed into two structural modules. Our work reveals surprisingly rich dynamics of post-transcriptional reactions and a potentially widespread mechanism useful for development and regeneration.

## INTRODUCTION

Gene expression variations caused by non-genetic factors are widely observed in mammalian cells (Chang et al., 2008; Jordan et al., 2016; Min and Spencer, 2019; Shaffer et al., 2017; Spencer et al., 2009). These variations have functional consequences such as altered differentiation potentials of stem cells and drug resistance of cancer cells (Abranches et al., 2014; Chang *et al*., 2008; Jordan *et al*., 2016; Peláez et al., 2015; Shaffer *et al*., 2017). For progenitor cells that exhibit multimodal expression patterns, a small subpopulation with a relatively homogenous expression profile recovers the parental population’s heterogeneity of individual gene products after several days (Chakraborty et al., 2020; Chang *et al*., 2008; Kalmar et al., 2009; Pina et al., 2012) (Fig. 1 box). Although stochasticity in transcriptional activities can cause expression variation and associated cell state changes (Raj et al., 2006; Shaffer *et al*., 2017), this type of noise influences expression at a much faster timescale (minutes) than the fluctuations required to achieve observed cell state transitions (days) (Chakraborty *et al*., 2020; Chang *et al*., 2008; Corrigan et al., 2016; Jordan *et al*., 2016; Raj and Van Oudenaarden, 2008). It was proposed that deterministically oscillatory dynamics may also be necessary for the recovery of heterogeneity (Huang, 2009; Kalmar *et al*., 2009; Udomlumleart et al., 2021), but instances of transcriptional network structure (e.g. transcriptional negative feedback loop) supporting the observed dynamics have not been found experimentally. Furthermore, how stochasticity works together with oscillation to recover cellular heterogeneity remains elusive. Finally, the multimodality of the expression patterns suggests the possibility of multistability (i.e. co-existing point attractors, Fig. 1 top right) (Enver et al., 2009; Huang, 2009). While both oscillatory and multistable systems allow slow cell state changes, they seem to contradict each other in terms of the underlying regulatory networks (Thomas, 1981), and it is unclear which mechanism restores the heterogeneous patterns more robustly.

**Figure 1.**
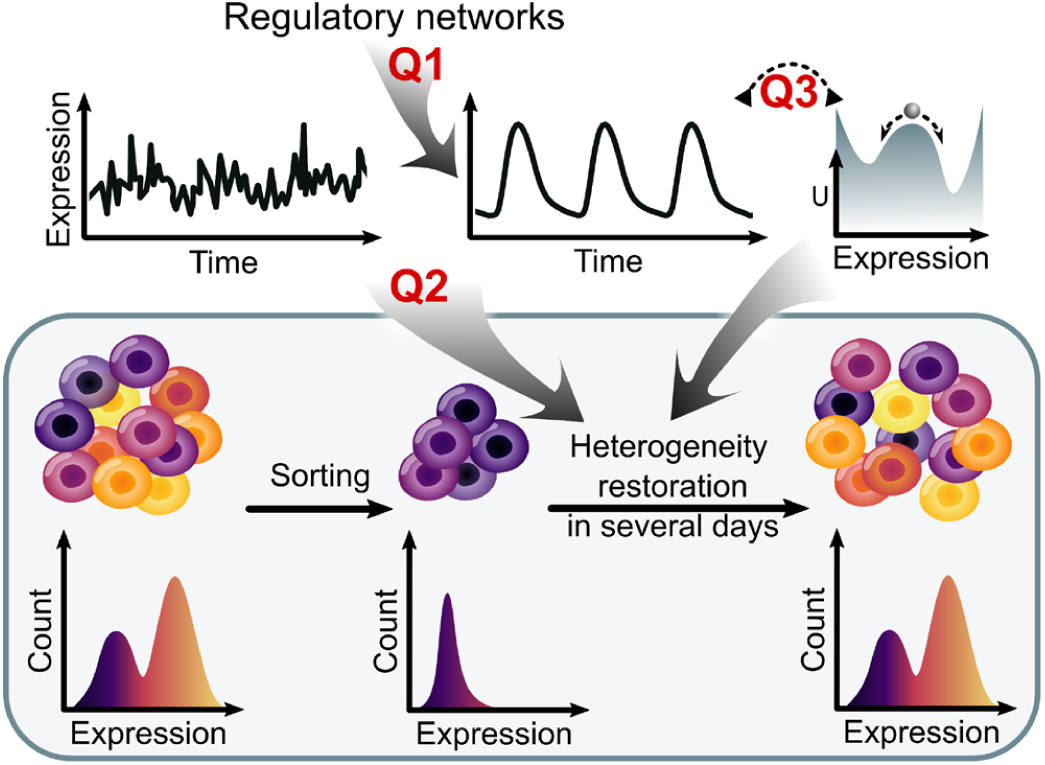
Questions related to dynamics of heterogeneity restoration process of expression patterns in progenitor cells. The study aims to provide insights into three questions: 1) Is there a widespread and hitherto unknown regulatory network structure that can generate oscillatory dynamics of gene expression at timescales of hours and days? 2) How do oscillation and stochasticity work together to drive restoration of heterogeneity from a subpopulation with relatively homogeneous expression patterns? 3) How can the theories of multistability and oscillation be reconciled in the context of heterogeneity restoration and multimodal distribution of gene expression?

Most human mRNA transcripts are subject to microRNA-mediated regulation at the post-transcriptional level (Miranda et al., 2006). Earlier findings indicated that microRNAs reduce gene expression noise through feed-forward loops (Li et al., 2009). More recently, however, it was shown that microRNA can also increase the variability in gene expression via more basic molecular mechanisms such as triggering mRNA degradation (Chakraborty *et al*., 2020; Schmiedel et al., 2015; Wei et al., 2021). In particular, the loss of microRNAs led to significantly reduced expression heterogeneity in embryonic stem cells (Chakraborty *et al*., 2020), and the lack of a microRNA binding site on a target mRNA dampened the oscillation of the target’s expression in neural progenitor cells (Bonev et al., 2012). These observations suggest versatile dynamics of microRNA-mRNA reaction networks and potential functions in maintaining heterogeneity in progenitor cells. Recent theories postulated that these reaction networks can produce positive-feedback-like dynamics such as bistability (Li et al., 2021; Tian et al., 2016), but it remains unclear whether more diverse types of dynamical features, such as oscillations, can be generated by the RNA-centric interactions.

Based on recent data of mRNA-microRNA interactions through multiple microRNA binding sites, we used mass-action kinetics to build mathematical models for simple post-transcriptional reaction networks. These networks potentially describe the dynamics of transcripts from nearly half of human protein coding genes. Through computational analysis of these common reaction networks, we identified regions of biologically plausible kinetic rate constants that give rise to sustained oscillations, despite the apparent absence of any imposed feedback loop typically considered necessary for oscillation. We found that the regions corresponding to oscillation and bistability overlap, which not only confers dual functions to the systems, but also allows excitability and abruptly diverging period of oscillations. The oscillatory and the emergent dynamics provide a new explanation for the experimentally observed heterogeneity and multimodality regeneration effects of microRNAs at timescales of hours and days (Chakraborty *et al*., 2020; Schmiedel *et al*., 2015). Remarkably, we found that oscillation and bistability require the same simple set of molecular species, which revealed a dual-function network without structural modularity. Our results uncover surprisingly rich dynamics of post-transcriptional reaction networks widespread in biology and a previously underappreciated mechanism for regenerating multimodal gene expression in cell populations.

## RESULTS

### An mRNA with one microRNA binding site generates spiral sinks but not oscillation or bistability

We used mass action kinetics to describe the dynamics of an mRNA and a microRNA with ordinary differential equations (ODEs). In the first model, we considered an mRNA with one binding site for the microRNA (the MMI1 Model, Fig. 2A). The two molecular species can bind to each other and form a complex. Each species is produced through transcription and is degraded both in its unbound form and in the complex. We simplified the mass-action-based model (Supplemental Information Section 1) into the dimensionless form

**Figure 2.**
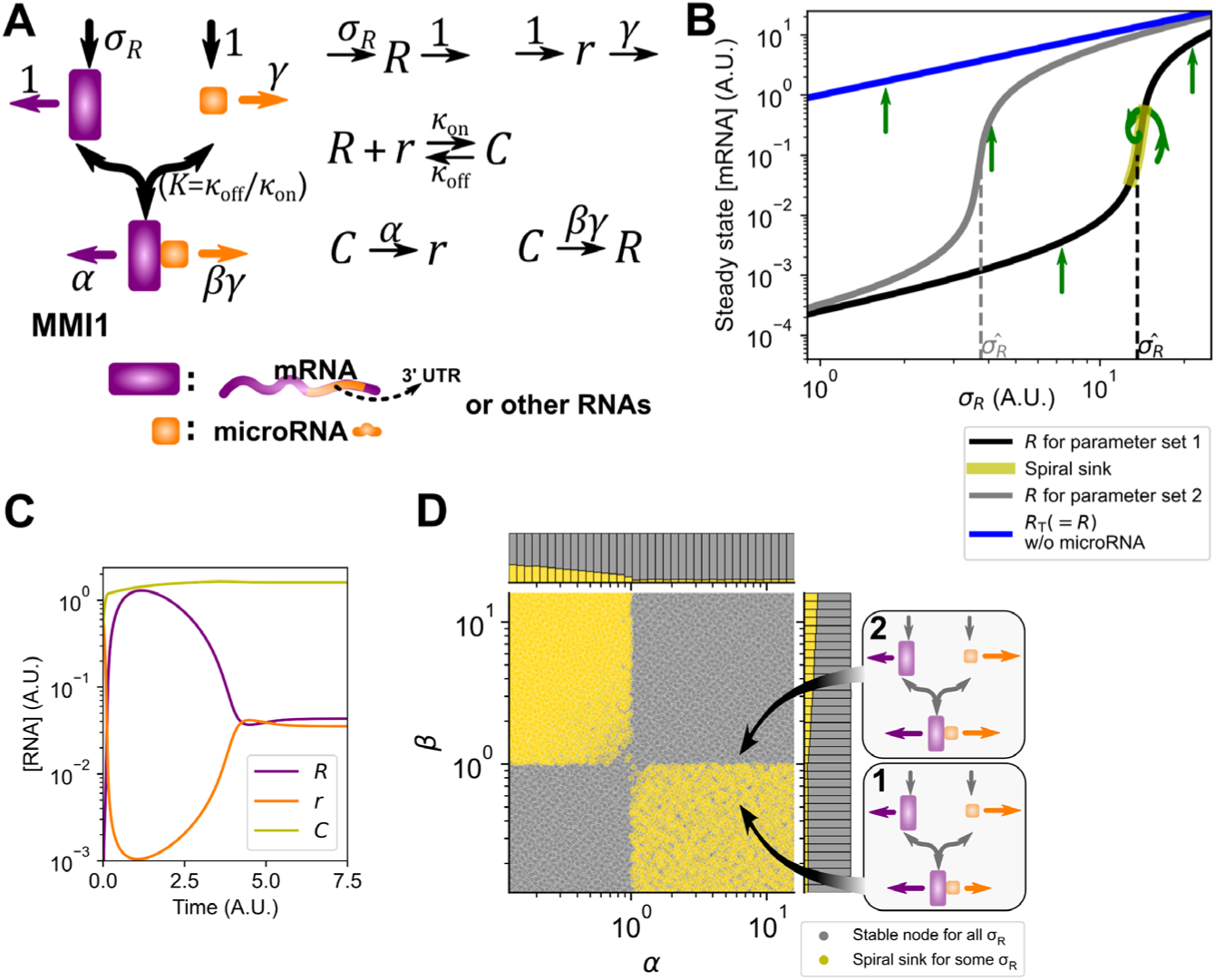
Dynamics of the MMI1 Model. (**A**) Left: illustration of reaction network of the MMI1 Model. Purple icon represents mRNA. Orange icon represents microRNA. Horizontal arrows represent degradation. Straight vertical arrows represent synthesis. Curved arrows represent binding. Right: eight chemical reactions associated with the MMI1 Model, which describes the dynamics of each molecular species with the law of mass-action (equation 1 and Supplemental Information Section 1). UTR, untranslated region. (**B**) Two representative signal-response curves (black and gray) showing steady state levels of *R* in response to transcription rate constant *σ*_*R*_. Green arrows indicate the type of these steady states: straight arrows show stable nodes and spiral arrow shows spiral sinks. Blue curve shows the microRNA-free response for both parameter sets. (**C**) Time-course simulation showing transient oscillation near a spiral sink (parameter set 1). One time unit is approximately 1.*44* × *t*_1/2_ where *t*_1/2_ is the half-life of the mRNA. (**D**) Distribution of the parameter sets with spiral sink steady state (yellow, e.g. parameter set 1) and those without (gray, e.g. parameter set 2) in the space of two representative parameters (RDFs). Callouts illustrate the two representative parameter sets (same as those in B) with degradation rate constants represented by arrow lengths. Only the values of *β* differ between the two sets. Marginal distributions are shown in stacked bars.

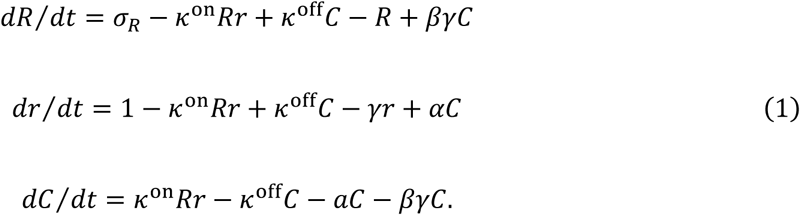

In this model, the three variables *R, r* and *C* represent the dimensionless (scaled) concentrations of unbound mRNA, unbound microRNA, and mRNA-microRNA complex respectively. The parameters (Greek letters) were also scaled, and each is related to a biologically meaningful parameter. *σ*_*R*_ represents the transcription rate constant of mRNA and can be considered a signal input. *k*^on^ represents the association rate constant. *k*^off^ represents the dissociation rate constant. The scaled dissociation constant *K* = *k*^off^/*k*^on^. The degradation rate constant of unbound mRNA and the microRNA production rate constant were scaled to 1 (see more details in Supplemental Information Section 1.1). The degradation rate constant of unbound microRNA is represented by *γ*. We define *α* and *β* as regulated degradation factors (RDFs). *α* represents how fast mRNA is degraded in the complex relative to its unbound form, and *β* is the corresponding factor for microRNA. These two RDFs are important because the gene regulatory function of microRNA primarily depends on the target mRNA degradation upon binding (Eichhorn et al., 2014), and similarly, the target mRNA can alter the degradation rate constant of the mRNA-bound miRNA (de la Mata et al., 2015). Here, we assumed that mRNA and microRNA are degraded independently in the complex, which is supported by previous observations (Baccarini et al., 2011; de la Mata *et al*., 2015; Ghini et al., 2018). Equation 1 can be simplified with representation of rapid chemical processes (e.g. binding and unbinding) using algebraic equations and change of variables to *R*_T_ and *r*_T_, which represent the total scaled concentrations of the mRNA and the microRNA respectively (Borghans et al., 1996; Ciliberto et al., 2007) (see Supplemental Information Section 1.2).

To explore the possible dynamical behaviors of the MMI1 Model, we randomly generated 10^5^ parameter sets with biologically plausible values. Each parameter value was chosen from a range covering at least two orders of magnitude (see Section 1.5 in Supplemental Information for estimation of each parameter). Through numerical bifurcation analysis with respect to signal *σ*_*R*_, which shows steady state signal-response relationships, we found that each parameter set gave rise to a single stable steady state, i.e. point attractor (Fig. 2B shows two representative signal-response curves). Nonetheless, we found that 13.4% of parameter sets generated extremely transient oscillations in narrow ranges of *σ*_*R*_ (Fig. 2B-C, Figs. S1 through S3). In the deterministic systems studied here, we define *sustained oscillation*, abbreviated *oscillation*, as limit cycle oscillation, which is a long-term periodic dynamical pattern. In contrast, *damped oscillation* is transient, characterized by a spiral trajectory which eventually converges to a point attractor, i.e. a spiral sink (Fig. 2C) (Strogatz, 2000). Regardless of whether the damped oscillation was present, the response curves with respect to *σ*_*R*_ show that with a single binding site, the microRNA enabled a threshold at which the unbound mRNA concentration starts to increase significantly (Fig. 2B, Fig. S2), consistent with previous experimental data and models (Mukherji et al., 2011). Furthermore, the presence of the microRNA only gave rise to moderate (< 10-fold) changes of the total mRNA concentrations (Fig. S2). This is consistent with the commonly observed moderate effects of microRNA on protein production, showing that the selected values of RDFs and other parameters in our models are realistic. Interestingly, the damped oscillations primarily occurred near the thresholds (estimated as the level of *σ*_*R*_ at *R* = 0.1, denoted by 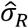) of the unbound mRNA activation (Fig. S2A) and required negatively correlated RDFs (Fig. 2D). In general, the damped oscillations did not strongly depend on the choice of specific values for individual parameters (Fig. S3).

We next used algebraic approaches to corroborate the computational results. We found that the MMI1 Model gives rise to a single stable steady state with any arbitrary combination of positive rate constants (Section 1.3 in Supplemental Information), suggesting that some additional structural component (i.e. molecular species) is required to achieve oscillation.

### Multiple microRNA binding sites enable sustained oscillation and bistability without imposed feedback

In this section, we discuss several variants of the MMI2 Model that describes an mRNA with two binding sites for one or two microRNAs, a biological system more common (see Methods) than the one captured by the MMI1 Model. The inclusion of additional binding sites permits several possibilities in terms of the binding reactions, but we first considered a very basic mechanism of binding: one microRNA binds to the two sites independently with equal affinities. This assumption gave rise to two 1:1 complexes, each with a microRNA molecule bound to the first site (Site 1) or the second site (Site 2) on the mRNA, respectively (Fig. 3A). Because these two complexes were assumed to have identical kinetic properties, their concentrations are always equal in a deterministic system. We therefore used a single state variable *C*_1_ to describe the concentration of each complex. Under this assumption, a 2:1 complex, with concentration denoted by *C*_2_, is formed when the microRNA binds to either 1:1 complex. We named this version of the MMI2 Model with symmetrically sequential binding the MMI2-SSB Model, which is given by

**Figure 3.**
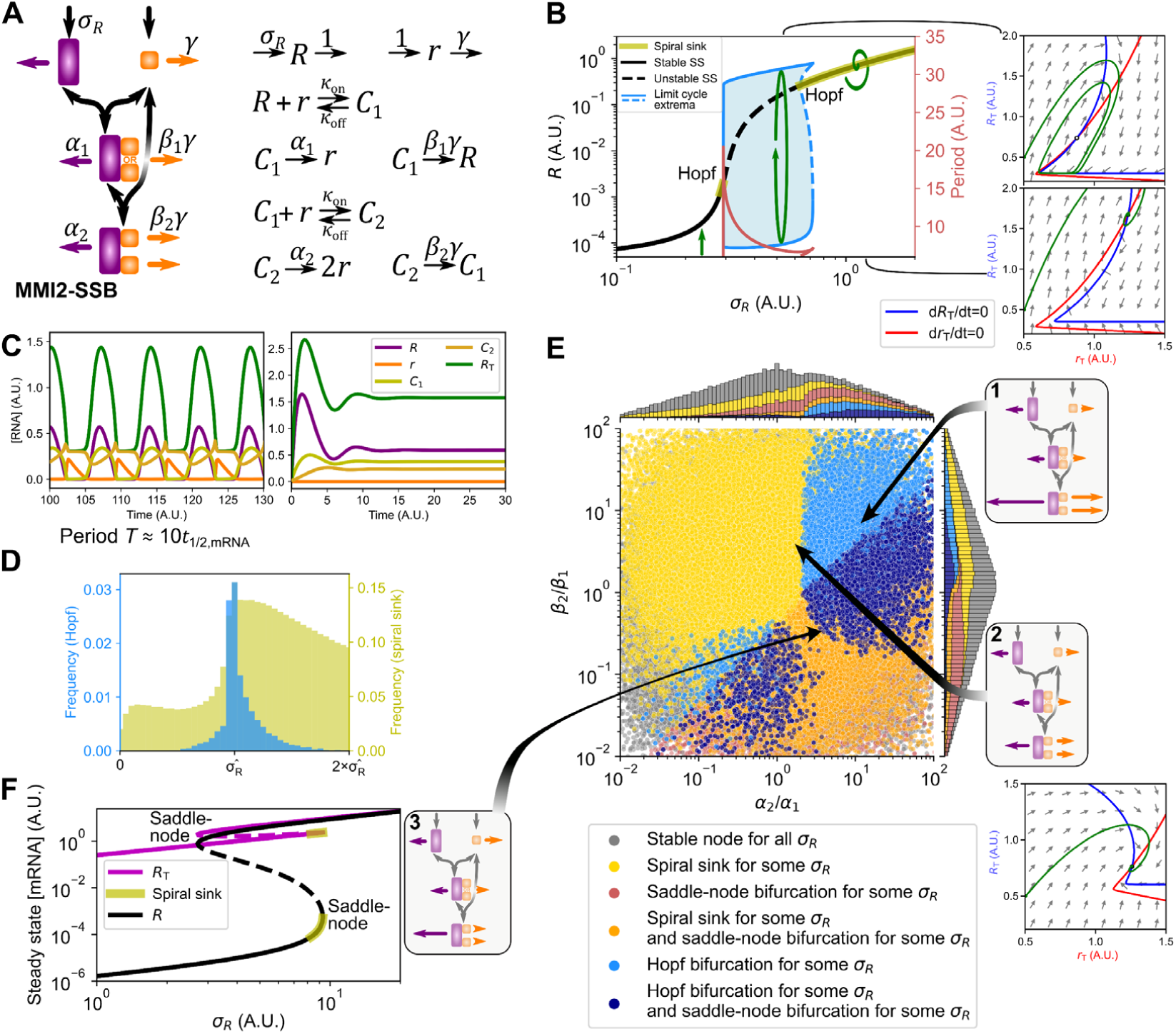
Dynamics of the MMI2-SSB Model. (**A**) Left: illustration of reaction network of the MMI2-SSB Model. Purple icon represents mRNA. Orange icon represents microRNA. Right: twelve chemical reactions associated with the MMI2-SSB Model in equation 2, which describes the dynamics of each molecular species with the law of mass-action. (**B**) Left: bifurcation diagram showing levels of *R* in response to transcription rate constant *σ*_*R*_. Green arrows illustrate the type of these steady states: straight arrow shows stable node, spiral arrow shows spiral sinks, and circulating arrow shows limit cycles. Blue shade: limit cycles’ inner basins of attraction. Right: phase planes (constructed with the 2D version of the MMI2-SSB Model) show sustained (*σ*_*R*_ = 0.5, top) and transient oscillations (*σ*_*R*_ = 1, bottom). Open circle represents unstable steady state. Blue and red curves are nullclines. Green curves show representative solutions. Other parameter values: *K* = 0.001, *β* = 0.25, *α*_1_ = *β*_1_ = 1, *α*_2_ = 12, *β*_2_ = 7. A basal synthesis rate constant 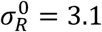 was added to the ODE for *R*. (**C**) Time-course trajectories for the two scenarios shown in B. One time unit is approximately 1.*44* × *t*_1/2_ where *t*_1/2_ is the half-life of the mRNA. (**D**) Yellow histogram shows the distribution of the spiral sink steady states relative to the position of the threshold transcription rate constant 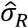 at which *R* = 0.1. Blue histogram shows the distribution of Hopf bifurcation points. € Left: distribution of the parameter sets with various types of steady states obtained with bifurcation analysis with respect to *σ*_*R*_ in the space of the RDF ratios. Callouts show representative parameter sets with arrow lengths representing degradation rate constants (Set 1 was used for results shown in B and C). Phase plane shows a transient oscillation obtained with Set 2. Marginal distributions are shown in stacked bars. (**F**) Left: bifurcation diagram showing the steady states of unbound mRNA and total mRNA with respect to *σ*_*R*_. Solid curves: stable steady states. Dashed curves: unstable steady states. Right: illustration of Set 3 whose behavior is shown. Parameter values: *K* = 0.001, *γ* = 2, *α*_1_ = 1, *β*_1_ = 0.5, *α*_2_ = 4, *β*_2_ = 0.1.

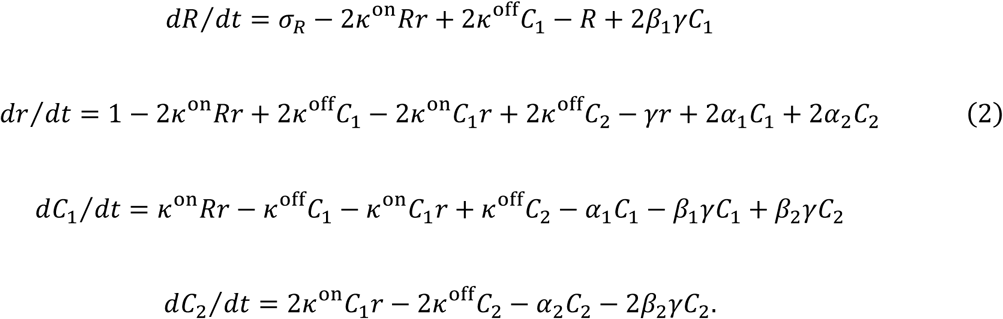

Similar to the MMI1 Model, this model can be reduced to two dimensions representing ‘slow’ variables *R*_T_ and *r*_T_. Strikingly, when we used the sampling strategy described earlier, we found many biologically plausible parameter sets that produced sustained oscillations at some *σ*_*R*_ (9.4% of the 10^5^ sets; Fig. 3B-C, Figs. S4 through S6). In this scenario, the system undergoes a Hopf bifurcation that leads to limit cycle oscillations at intermediate levels of *σ*_*R*_ (Fig. 3B blue shade and top right phase plane). Further increase of *σ*_*R*_ gives rise to another Hopf bifurcation point which marks the approximate upper bound in the *σ*_*R*_ axis for limit cycle oscillations.

With representative parameter sets, damped oscillations occurred at high levels of *σ*_*R*_ (Fig. 3B yellow shade and lower right phase plane). Unlike the extremely transient oscillations with the MMI1 Model, these damped oscillations can have significantly long durations (Fig. 3C) and can be observed in wide ranges of parameter values including *σ*_*R*_ (Fig. 3E, Fig. S4). In the limit cycle oscillations, dramatic variations in fold change of unbound mRNA and microRNA were observed (Fig. 3B, Fig. 3C purple and orange), whereas moderate (≈3-fold) changes of total mRNA occurred, again confirming the moderate overall effect of microRNA (blue curves in Fig. 3B phase planes and Fig. 3C). With the parameter set that enabled sustained oscillation, we found that the period of the oscillation was approximately 10 times of the half-life of the mRNA (Fig. 3C), which corresponds to at least one day for typical mammalian mRNAs (Sharova et al., 2009). Nonetheless, a wide range of periods for biological rhythms might be obtained by the model given the wide distribution of mRNA half-lives (Sharova *et al*., 2009). More importantly, the period increased steeply when the signal approached the Hopf bifurcation near the activation threshold (Fig. 3B red). We will examine the significance and the source of this phenomenon in a later section.

With an estimated threshold of 0.1 units of unbound mRNA, we found that many Hopf bifurcation points were located near the activation threshold, whereas the damped oscillations primarily appeared in the high *σ*_*R*_ region (Fig. 3D). The Hopf bifurcation points were distributed very widely in the space of all other parameters (Fig. S5). To illustrate the regions corresponding to these bifurcations, we focused on a phase space of RDF ratios for the two RNAs, i.e. *α*_2_/*α*_1_ and *β*_2_/*β*_1_ (Fig. 3E). These ratios represent how fast the mRNA and the microRNA are degraded in the 2:1 complex relative to their degradation rate constants in the 1:1 complex, and can be viewed as a functional cooperativity, or synergy, between the two microRNA binding sites. We use the term *functional cooperativity* here to distinguish it from binding cooperativity that often describes the enhanced or reduced binding affinity of a second site upon the binding to the first site. Previous observations suggested that in most cases the two sites have a positive functional cooperativity in terms of the degradation rate constants of mRNAs, and in fact the *α*_2_: *α*_1_ ratios are often equal to or greater than 2 (Grimson et al., 2007). The RDF ratio for microRNA (*β*_2_/*β*_1_) has not been systematically characterized, but we permitted both positive and negative cooperativities. Very interestingly, most Hopf bifurcations and their associated oscillations appeared in a region where RDF ratios (*α*_2_/*α*_1_ and *β*_2_/*β*_1_) are both high (e.g. Parameter Set 1 in Fig. 3E), whereas a small fraction of Hopf bifurcations were observed with low RDF ratios (Fig. 3E lower left region). Damped oscillations were observed in 41.8% of parameter sets (Fig. 3E yellow and orange), including areas of RDF ratio space where Hopf bifurcations were absent (e.g. Parameter Set 2 and phase plane in Fig. 3E).

In addition to oscillations, we found that a large number of parameter values (31.7%) generated bistable systems characteristic of biological switches (e.g. Parameter Set 3 in Fig. 3E-F), a conclusion consistent with a previous study (Li *et al*., 2021). Like oscillations, the bistable switches involved dramatic changes of unbound mRNA concentration and moderate changes of total mRNA concentrations (Fig. 3F). Bistable switches with respect to signal *σ*_*R*_ were observed only if the inequality

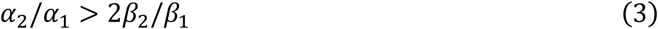

was satisfied (Fig. 3E, boundary between light blue and dark blue regions).

We found that the two parameter regions corresponding to oscillation and bistability respectively had a significant overlap in the space of RDF ratios (Fig. 3E, dark blue), and we will explore this result in a later section. In addition to the RDF ratios, we found that the existence of the sustained oscillations depends on the choice of *γ*. In our simulations, the median of the distribution for *γ* was chosen to be 0.25, which is supported by previous experimental studies (Zlotorynski, 2019). We observed that the oscillations were obtained with high RDF ratios (high *α*_2_/*α*_1_ and high *β*_2_/*β*_1_) when *γ* was low, whereas oscillations with low RDF ratios required high *γ* (Fig. S6). Nonetheless, many biologically plausible values of the parameters in the MMI2-SSB Model, including the median values of distributions for *γ* and *K* in our sampling experiments, gave rise to bistability and oscillation (Fig. S6). In addition, we found that coregulated transcription of microRNA and its target significantly expanded the parameter region for limit cycles (Fig. S7). This type of coregulation may be achieved by localization of a microRNA gene in the intronic region of its target gene, e.g. *mir-196* and its target *Hoxb7* (McGlinn et al., 2009).

The dynamical profile shown in Fig. 3B provides a possible explanation for recent data that showed perplexing roles of microRNAs: Schmiedel et al. and Wei et al. found that microRNA reduces variability of gene expression for lowly expressed genes but increases the variability for highly expressed genes (Schmiedel *et al*., 2015; Wei *et al*., 2021). While the increased variability could be explained by additional stochasticity introduced by microRNA-mediated regulations (Schmiedel *et al*., 2015), our analysis suggests that the observed variability of highly expressed genes controlled by microRNAs could alternatively be due to the spiral nature of the steady state, which may also be related to functional rhythms. Nonetheless, the observation of limit cycle oscillations in the MMI2-SSB Model is surprising because the network structure does not contain any imposed negative feedback loop, a structure considered a necessary condition for biological oscillators (Novák and Tyson, 2008). Furthermore, it is remarkable that a simple system containing so few molecular species can possess very diverse dynamical features.

We next asked whether the observed oscillation and bistability were sensitive to the assumption that the two binding sites were identical in terms of kinetic properties, or the fact that the two 1:1 complexes were described by a single variable *C*_1_ in the model. We therefore considered a modified MMI2 Model in which the binding of microRNA to Site 2 on mRNA requires the binding of Site 1. This asymmetrically sequential binding model, named MMI2-ASB Model, now contains a unique 1:1 complex corresponding to the first binding site occupied by the microRNA (Fig. 4A). We found that this model generated results similar to the MMI2-SSB Model in terms of the parameter regions for oscillation and bistability: 8.4% of parameter sets gave rise to Hopf bifurcation and oscillations, whereas 29.6% of parameter sets produced bistable switches (Fig. 4B). Because the two binding sites in the MMI2-ASB Model were assumed to be distinct, we asked whether the difference in the binding affinity (described by the dissociation constant *K*) can influence the emergence of oscillation and bistability. To test this, we randomly chose *K*_1_ (the scaled dissociation constant for Site 1) as before and set *K*_2_ (for Site 2) equal to *K*_1_ multiplied by a constant. We found that positive binding cooperativities (*K*_2_ < *K*_1_) had a negative effect on producing oscillation and bistability, whereas negative binding cooperativities (*K*_2_ > *K*_1_) enhanced the ability to generate oscillation and bistability until the binding affinity for Site 2 became too low (Fig. 4C). Nevertheless, large numbers of parameter sets produced spiral sinks, oscillation and bistability over a wide range of *K*_2_: *K*_1_ ratios.

**Figure 4.**
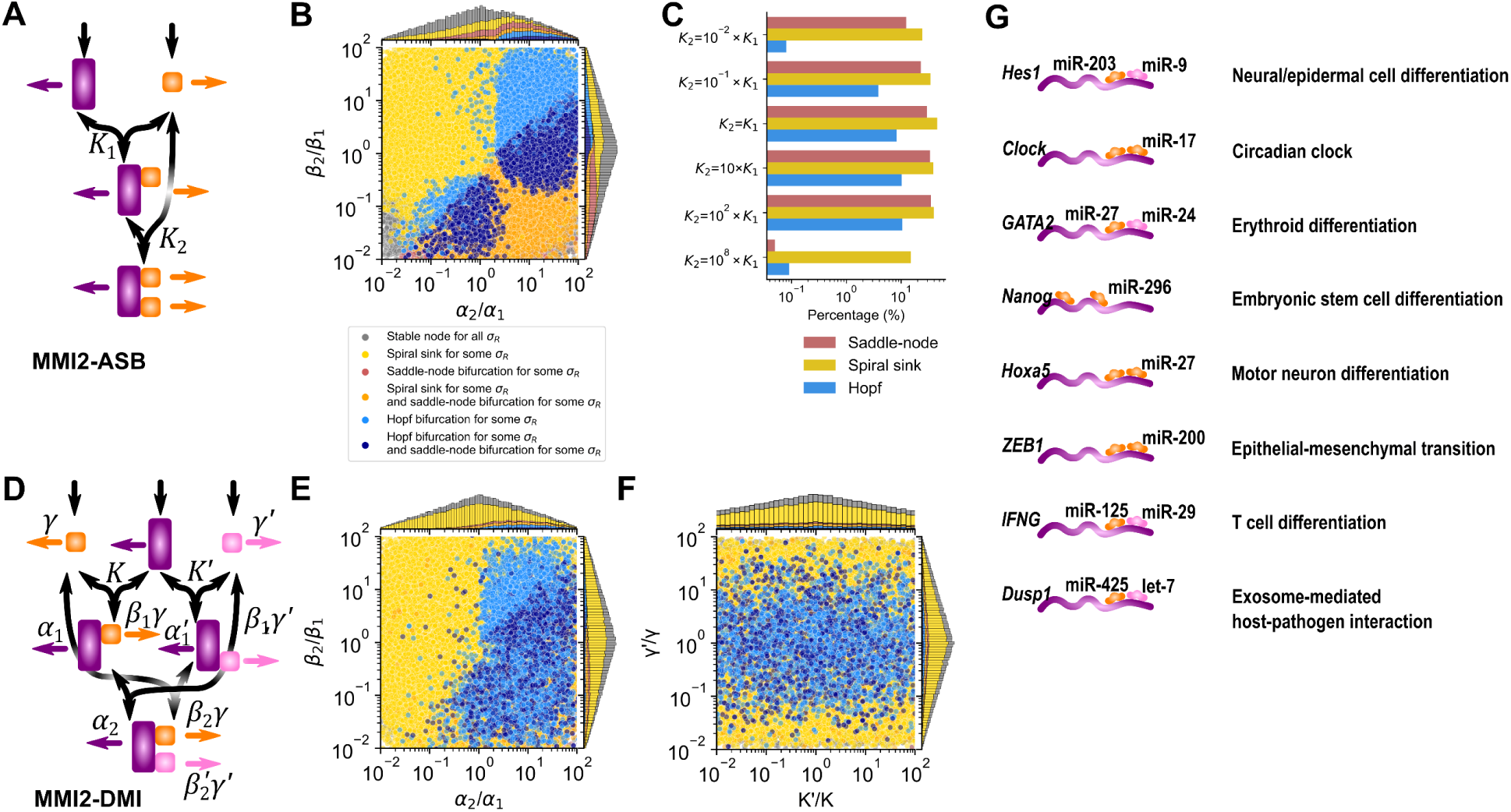
Dynamics of the MMI2-ASB and MMI2-DMI Models. (**A**) Illustration of the MMI2-ASB reaction network. (**B**) Distribution of the randomly selected parameter sets for MMI2-ASB with various types of stead states obtained with bifurcation analysis with respect to *σ*_*R*_ in the space of the RDF ratios. (**C**) Percentages of parameter sets with various types of stead states obtained with bifurcation analysis with respect to *σ*_*R*_. Parameters were randomly selected as in B, except for *K*_2_ which was equal to *K*_1_ multiplied by the indicated factors. (**D**) Illustration of the MMI2-DMI reaction network. Orange and pink icons represent two different microRNAs. € Distribution of the randomly selected parameter sets for MMI2-DMI with various types of stead states obtained with bifurcation analysis with respect to *σ*_*R*_ in the space of the RDF ratios. Parameter values for the two microRNAs are uncorrelated. (**F**) Same data as in E shown in the space of ratios between the parameters for the two microRNAs. In B, E and F, marginal distributions are shown in stacked bars. (**G**) Examples of MMI2 systems with experimental evidence that support direct binding. These circuits are relevant to neural or epidermal cell differentiation (Phillips et al., 2016; Zhou et al., 2018), circadian rhythm (Gao et al., 2016), erythroid differentiation (Wang et al., 2014), embryonic cell differentiation (Tay et al., 2008), motor neuron differentiation (Li et al., 2017; Li *et al*., 2021), epithelial-mesenchymal transition (Burk et al., 2008), T cell differentiation (Ma et al., 2011; Pan et al., 2015), and host-pathogen interaction (Buck *et al*., 2014) respectively.

Because cooperativity among multiple microRNAs in cellular functions has been observed previously (Buck et al., 2014; Cursons et al., 2018), we asked whether the conclusions about oscillation and bistability with the MMI-2 Models can be extended to the scenario where the two binding sites are recognized by two different microRNAs. In this dual-microRNA model (the MMI2-DMI Model, Fig. 4D), two microRNAs were explicitly described and assumed to each bind only their respective site. As expected, with the assumption that the two microRNAs have identical rate constants, the MMI2-DMI Model had similar performance as the MMI2-SSB Model: 8.7% of parameter sets produced oscillation, and 28.8% of parameter sets produced bistability. With the assumption that the two binding sites have distinct, independently chosen dissociation constants and other associated parameters such as RDFs, we again observed similar results in terms of the capacity of oscillation (4.9%) and bistability (13.8%). These dynamical features did not require high similarities of the two microRNAs or the two binding sites (Fig. 4E-F).

We have shown that under several different assumptions, the MMI2 Models generated spiral sinks, oscillation and bistability in biologically plausible parameter regions. We asked how many human protein-coding genes may be directly involved in the MMI Models. With TargetScan, a microRNA binding site prediction program, we found 2554 human protein-coding genes whose 3’ untranslated regions contain only one conserved microRNA binding site (the structure of the MMI1 Model), whereas 10483 genes were predicted to contain two or more conserved binding sites (the structures of the MMI2 Models) (Agarwal et al., 2015). Furthermore, for 8420 of the genes with two putative binding sites, there is experimental evidence that supports two or more microRNA families both targeting each gene (Huang et al., 2020), as in the MMI2-DMI Model structure. We therefore estimate that nearly half of human protein-coding genes are involved in the structures of the MMI2 Models. In contrast, only 62 genes were predicted to be involved in transcriptional negative feedback loops with up to three edges (see Methods). Several examples of the MMI2-like systems with experimental evidence of RNA binding and functional significance are shown in Fig. 4G (Buck *et al*., 2014; Burk *et al*., 2008; Gao *et al*., 2016; Li *et al*., 2017; Li *et al*., 2021; Ma *et al*., 2011; Pan *et al*., 2015; Phillips *et al*., 2016; Tay *et al*., 2008; Wang *et al*., 2014; Zhou *et al*., 2018).

In the next sections, we first discuss the characteristic global features of MMI2-driven dynamics beyond saddle-node and Hopf bifurcations. Next, we explore the possible biological functions of the microRNA-driven oscillators. Finally, we test the decomposability of the MMI2 Model into an oscillator module and a bistable switch module with distinct molecular compositions.

### Multiple microRNA binding sites enable global bifurcation and robust regeneration of heterogenous expression

Because the Hopf bifurcations and bistable switches both occurred near the thresholds of the mRNA activation (Fig. 3D), we hypothesized that the MMI2-SSB Model can generate global bifurcations with features not captured by the local Hopf and saddle-node bifurcations. We examined the parameter sets that gave rise to both Hopf and saddle-node bifurcations with respect to *σ*_*R*_ (Fig. 3E dark blue) and found that all of them produced global bifurcations: about one third of the sets generated saddle-node on invariant circle (SNIC) bifurcations and the rest generated saddle-loop bifurcations (Fig. 5A). During SNIC bifurcation, a saddle-node bifurcation point collides with a limit cycle (Fig. 5B), giving rise to an abrupt appearance of periodic trajectories as *σ*_*R*_ increases gradually (e.g. Fig. 5C-D). The limit cycle eventually disappears through a Hopf bifurcation with further increase of *σ*_*R*_. The periods of limit cycles diverge dramatically near the SNIC bifurcation point, going to infinity at the bifurcation point (Fig. 5B, red).

**Figure 5.**
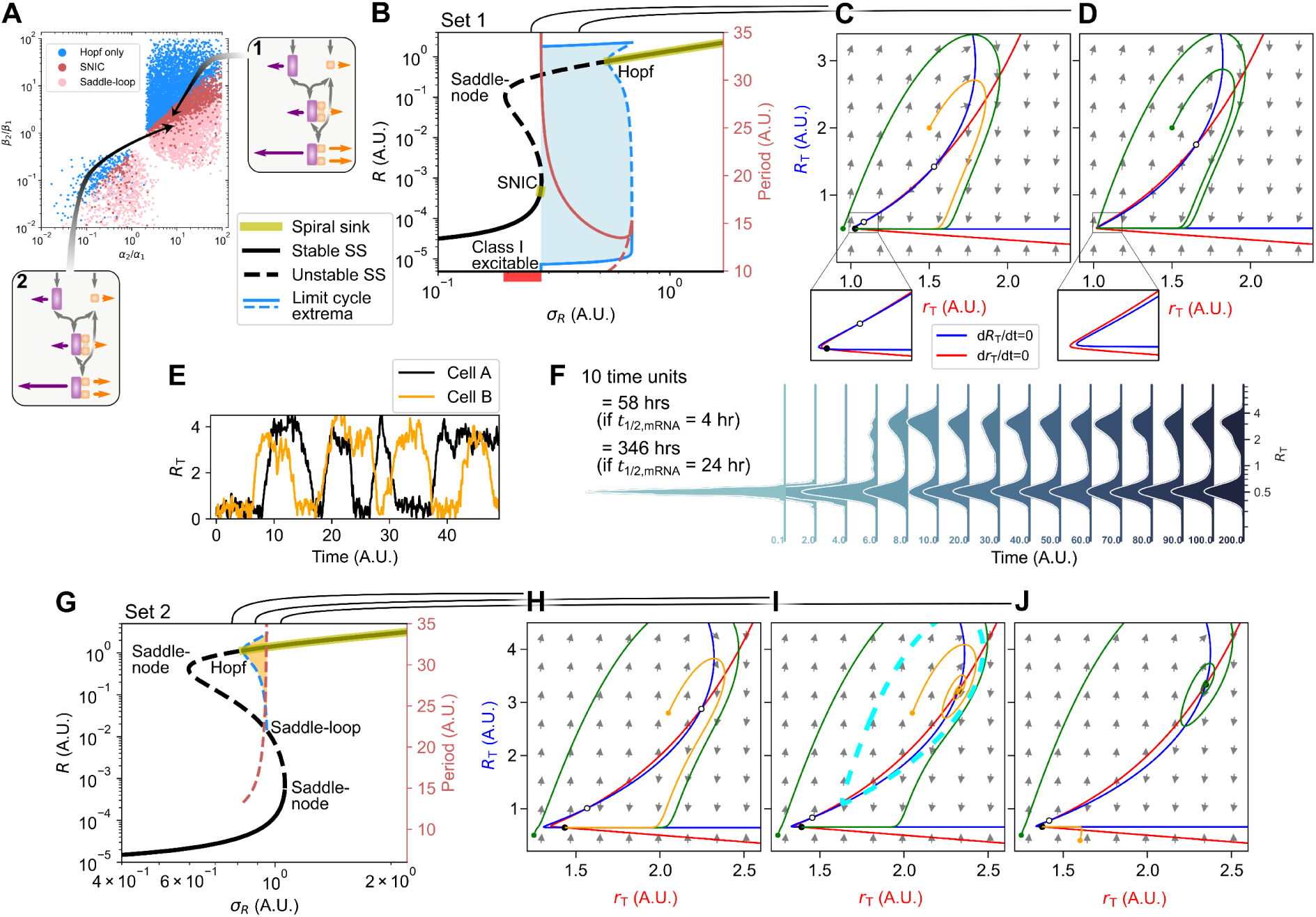
Global bifurcations of the MMI2-SSB Model. (**A**) Scatter plot shows parameter sets producing only Hopf bifurcation (blue) and those producing both Hopf and saddle-node bifurcations (same as Fig. 3 dark blue) in the space of RDF ratios. Two representative sets are shown in callouts. (**B**) Bifurcation diagram shows levels of *R* in response to transcription rate constant *σ*_*R*_. Blue shade: limit cycles’ inner basins of attraction. (**C**-**D**) Phase planes show vector fields (gray), nullclines (red and blue), and representative solutions (green and orange) for *σ*_*R*_ = 0.27 and *σ*_*R*_ = 0.4, respectively. Open and closed circles represent unstable and stable steady states respectively. Other parameter values: *K* = 0.001, *γ* = 0.25, *α*_1_ = *β*_1_ = 1, *α*_2_ = 12, *β*_2_ = 4, and basal mRNA synthesis rate 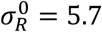. (**E**) Trajectories of stochastic simulation for two representative cells under a SNIC parameter set. Stochastic ODEs have the form *dx* = *f*(*x*)*dt* + *ω*_*x*_*d*W, where *x* represents either *R* or *r, f*(*x*) is the right-hand side of the first two ODEs in equation (2), and *d*W denotes the Wiener process. *ω*_*R*_ = 1.4. *ω*_*r*_ = 0.*3*5. *σ*_*R*_ = 0.*3*. Other parameter values are the same as in B-D. One time unit is approximately 1.*44* × *t*_1/2_ where *t*_1/2_ is the half-life of the mRNA. The initial condition is the deterministic steady state solution obtained with *σ*_*R*_ = 0.1. (**F**) Distributions of total mRNA concentrations in 500 simulated cells at the indicated time points. Initial conditions and parameter values are identical to E. (**G**) Bifurcation diagram of a model exhibiting a saddle-loop bifurcation showing levels of *R* in response to transcription rate constant *σ*_*R*_. Parameter values are the same as for B except *β*_2_ = *3* and 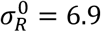. (**H**-**J**) Phase planes show vector fields, nullclines, and representative solutions of the saddle-loop model for *σ*_*R*_ = 0.7, *σ*_*R*_ = 0.85, and *σ*_*R*_ = 1, respectively.

Oscillators that appear abruptly with diverging periods give rise to a particular type of excitability that is known to govern asynchronous oscillators, Class I excitability (Fig. 5B red bar) (Ermentrout, 1996). We therefore hypothesized that the period-diverging property can be used to generate heterogeneous gene expression pattern in the presence of noise that may result from stochasticity in transcription. We first simulated the MMI2-SSB Model with additive transcriptional noise in RNA production and a *σ*_*R*_ level close to the SNIC bifurcation point. With an identical initial condition for multiple simulated cells, we observed asynchronous fluctuations of mRNA levels (Fig. 5E). The fluctuations contained both moderate, frequent changes, and dramatic, infrequent changes of mRNA levels. We define the latter as cell state changes. With 500 simulated cells starting from a low mRNA condition, significant cell state changes at the population level occurred only after at least one mRNA half-life (Fig. 5F). The gene expression pattern of the simulated cell population exhibited a damped oscillation and eventually converged to a bimodal distribution with stabilized fractions. The recovery of bimodal distribution from a selected homogeneous population, the slow onset of cell state transitions, and the nonmonotonic changes of gene expression distribution were consistent with previous observations in haematopoietic progenitor cells (Chang *et al*., 2008). Furthermore, the role of microRNA in multimodality regeneration at the timescale of days is consistent with recent observations in embryonic stem cells (Chakraborty *et al*., 2020). Our results suggest that the mechanisms underlying these features may include diverging oscillations with the influence of both SNIC-like dynamics and stochasticity.

Although SNIC bifurcation was observed in a relatively small number of parameter sets, its key features extend to a wide region. Sudden oscillation appearance and rapid period change also occurred in scenarios without a SNIC bifurcation. For a saddle-loop bifurcation, an unstable limit cycle is first generated by a Hopf bifurcation as *σ*_*R*_ increases (Fig. 5G-J). When *σ*_*R*_ increases further, a saddle point collides with the limit cycle, resulting in an abrupt disappearance of the limit cycle. Even for an oscillation-enabling parameter set without any global bifurcation, a limit cycle’s period, though finite, changes sharply near a Hopf bifurcation point (Fig. 3B). Because the SNIC bifurcation organizes a parameter region that generates limit cycles with rapid changes in period, we named the mechanism in the nearby parameter region a *diverging oscillator*.

Reversible cell state transitions and bimodal distribution of expression may alternatively be explained by a model describing two stable steady states (e.g. two point attractors, Fig. 3F). This *bistable switch* mechanism has been very widely used to explain cell states and their transitions (Li and Wang, 2013; Moris et al., 2016). We found that both diverging oscillator and bistable switch mechanisms produced bimodal distributions of gene expression at a basal level of transcriptional noise. To visualize the two distinct dynamical systems under both deterministic and stochastic influences, we plotted the quasi-potential landscapes for both mechanisms based on the results of stochastic simulations (Fig. 6A-D). Consistent with the bimodal gene expression distributions, both mechanisms generated double-well potentials along the total mRNA axis. However, we observed a striking difference between the two mechanisms in their routes for state transitions: because of the limit cycle, the diverging oscillator has two “valleys” connecting the two potential wells (Fig. 6A cyan), whereas the two wells in the bistable switch mechanism are separated by a saddle point and its associated separatrix (Fig. 6B yellow and orange). The double-valley-double-well landscape of the diverging oscillator allows rapid transition between two states driven by deterministic vector field, while the double-well landscape of the bistable switch only allows stochastic transitions between the two states (blue representative trajectories in Fig. 6A-B and Movie S1). Because of this difference, we hypothesized that the gene expression pattern driven by the bistable switch is more sensitive to the initial conditions and the noise levels. We therefore compared the performance of the two mechanisms in terms of bimodality regeneration with three levels of noise, two initial conditions, and three signal strengths. We first focused on the selected signal strengths *σ*_*R*_ allowing both mechanisms to produce bimodal distributions of mRNA expression at *t* = 100 (equivalent to 576 hours after cell sorting, assuming a 4-hour mRNA half-life) with the same levels of noise and mRNA-low initial conditions (Fig. 6C-D, blue in center panels). Next, we varied the noise level in both systems, finding that the bimodal distributions were retained with the diverging oscillator but not the bistable switch in both noise-reduced and noise-amplified situations (compare rows in Fig. 6C-D; Movies S1 and S2). Furthermore, when we changed the initial conditions to mRNA-high states, we observed significant alteration of gene expression patterns with the bistable switch under medium and low noise conditions, while the diverging oscillator was insensitive to the changes of initial conditions under all tested cases (compare blue and red populations in Fig. 6C-D).

**Figure 6.**
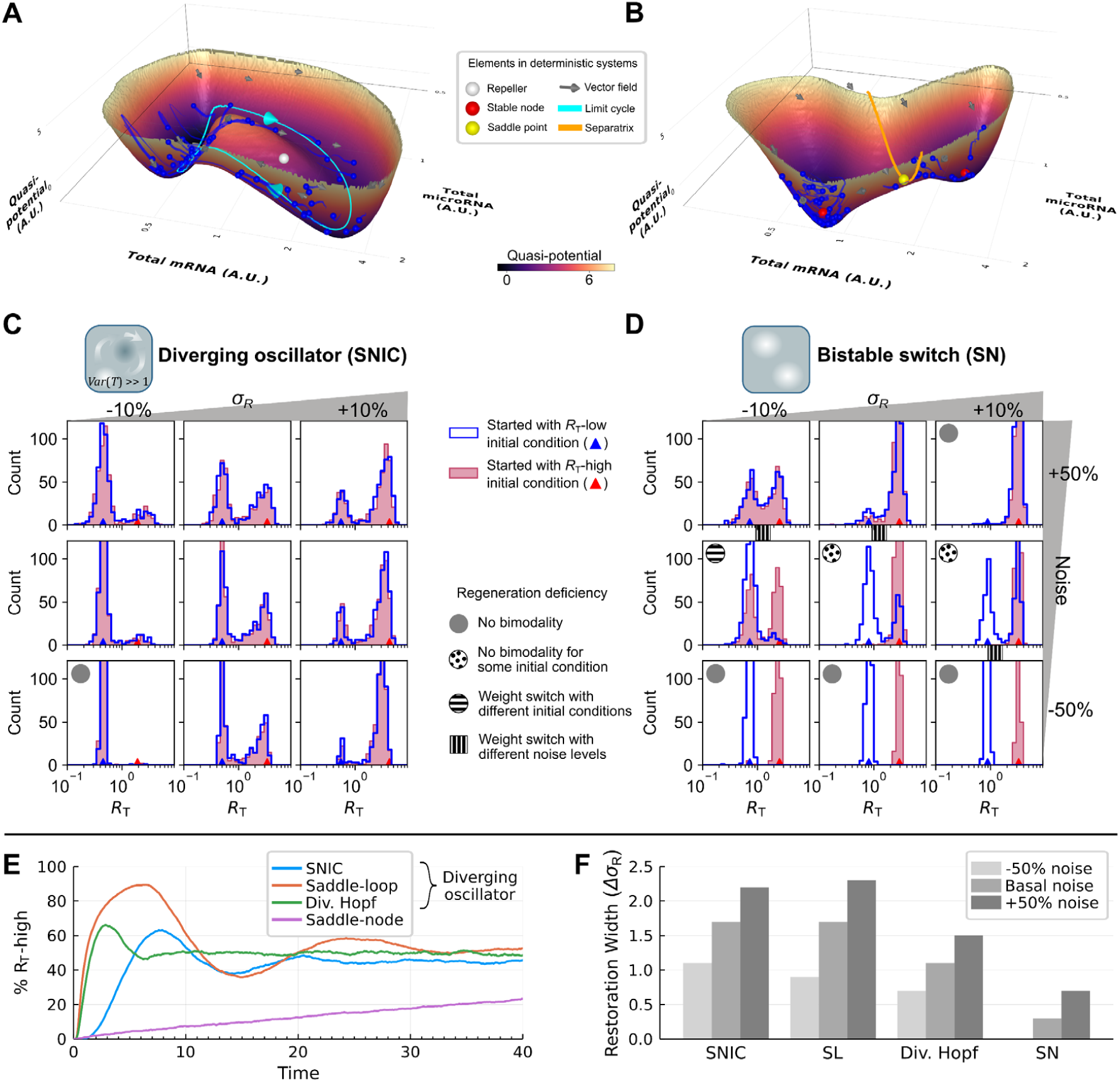
Comparison of MMI2-SSB mechanisms for regenerating bimodal gene expression distributions. (**A**-**B**) Landscapes of the MMI2-SSB Model with additive noise describing stochastic transcription of mRNA and microRNA (basal noise level *ω*_*R*_ = 1.4, *ω*_*r*_ = 0.*3*5). Landscapes were computed based on stationary phase distribution of stochastic simulations of 500 cells. For each landscape, 50 cells were randomly selected and visualized with the positions at *t* = 50 (blue spheres) and trajectories in a 0.4 time-unit period (blue tails). Parameter values for the model generating SNIC bifurcation (A) are the same as in Fig. 5B. Parameter values for the model generating SN bifurcation (B) are as in Fig. 3F. (**C**-**D**) Distributions of total mRNA concentrations at *t* = 100 (576 hours after selecting cells with extreme expression, assuming a 4-hour mRNA half-life) with three values of *σ*_*R*_, three levels of noise, two initial conditions, and two switch mechanisms from A and B. Noise levels for both mechanisms were identical in corresponding panels. The basal noise levels (middle row) are as in A and B. The *σ*_*R*_ values for the two middle columns are 0.3 and 3.2 for the two mechanisms, respectively. The *R*_T_ -low and *R*_T_ -high initial conditions were obtained by sampling 400 initial conditions and their corresponding extrema in the period *t* > 5. (**E**) Early timecourse of the proportion of *R*_T_-high cells in a 5000-cell population starting from the low initial condition under basal noise for several MMI2-SSB Models. For SNIC, *σ*_*R*_ = 0.*3* with *R*_T_ cutoff 1.0; for the saddle-loop (SL) model in Fig. 5G, *σ*_*R*_ = 0.9 with cutoff 1.25; for the diverging (div.) Hopf model in Fig. 3B, *σ*_*R*_ = 0.6 with cutoff 0.75; for saddle-node, *σ*_*R*_ = *3*.0 with cutoff 1.3. All other parameters are compiled in Table S1. (**F**) Width of the region of *σ*_*R*_ in which each model in E can restore bimodal gene expression.

The pronounced robustness of bimodality regeneration by the diverging oscillator compared to the bistable switch was in agreement with the consistent quasi-potential landscapes at different noise levels (Fig. S8A-B). Intuitively, changing noise levels (e.g. by altering the environment or cell volume) can significantly vary the heights of the middle barriers in both mechanisms, but the barrier lies in the main routes of state transitions only for the bistable switch (Fig. 6A-B, Fig. S8A-B) (Li and Wang, 2013). In addition, we observed that the diverging oscillator produced bimodality in a wide range of *σ*_*R*_ (compare columns in Fig. 6C-D), even when the limit cycles were absent (Fig. S8C). This is because the vector field characteristic of the limit cycle has a significant influence on its adjacent parameter region. Furthermore, for systems that have only Hopf bifurcations (Fig. 3B) and those have saddle-loop but not SNIC bifurcations (Fig. 5G), the double-valley-double-well landscapes were still retained (Fig. S8D-E). These results suggest that while SNIC bifurcation occurs in a small parameter region, it acts as an organizing center for the diverging oscillator that can be supported by widely distributed parameter values (Fig. 5A).

Next, we performed a more systematic comparison between three types of diverging oscillators (SNIC, saddle-loop, and diverging Hopf) and a bistable switch with additional stochastic simulations and a metric of bimodality based on Gaussian mixture models (Fig. 6E-F, Section 2.1.7 in Supplemental Information). We found that the diverging oscillators not only regenerated expression patterns with nonlinear time courses of cell fractions observed experimentally (Fig. 6E) (Chang *et al*., 2008), but also produced wider ranges of bimodality compared to the bistable switch (Fig. 6F, Fig. S10). Since all the MMI2-driven oscillations are diverging, we performed the same analysis with a previously known non-diverging oscillator, the repressilator (Elowitz and Leibler, 2000), and confirmed that the non-diverging oscillator cannot produce bimodality robustly (Fig. S16). We used two alternative approaches, multiplicative noise and the Gillespie algorithm, to model noise in the MMI2-SSB Model, confirming that our conclusions about the capabilities of the diverging oscillator are not sensitive to the types of noise (Figs. S9, S11, and S12). Finally, we explicitly incorporated cell division into the model, confirming that the results are applicable to the scenario of a proliferating cell population during heterogeneity recovery (Fig. 1 box, Fig. S13).

In conclusion, the diverging oscillator mechanism based on the MM2-SSB Model gave rise to more robust regeneration of multimodality gene expression patterns compared to the commonly used bistable switch mechanism. Nonetheless, our results do not imply that the diverging oscillator is the only mechanism for dynamics of all cells in a population of progenitor cells, which may contain both self-regenerative cells that exhibit reversible transitions and those stabilized in point attractors (Pina *et al*., 2012). Remarkably, the MMI2-SSB Model can support either mechanism with adjustment of rate constants.

### Sustained oscillation and bistability are achieved without structural modularity

We have shown that the diverging oscillators arise from adjacent limit cycles and saddle-node bifurcation points, which further suggests that the emergent functions of the MMI2 Models depend on their capacity of producing both oscillations and bistable switches. Bifunctional systems are often evaluated to consider whether the two functions are governed by distinct subnetworks. We therefore asked whether a MMI2 Model contains two molecular modules each responsible for one function. According to the definition by Jiménez et al. (Jiménez et al., 2017), the degree of modularity regarding two biological functions generated by a network can be described by the number of molecular species (e.g. genes) shared by two subnetworks necessary for achieving the two functions respectively, divided by number of molecular species in the union of the two subnetworks. This quantity is the Jaccard index

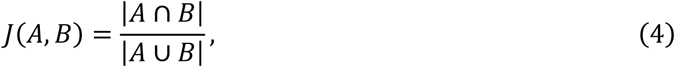

where *A* and *B* are two sets of nodes of the subnetworks necessary to achieve the two functions respectively. If the two subnetworks overlap completely in terms of the structural components (*J* = 1), then the two functions are completely structurally nonmodular in this network. This scenario has a profound implication in evolution: the acquisition of a new function does not require the inclusion of any new molecular species to the existing network; instead, the function can be obtained simply by adjusting the kinetic rate constants in the existing network.

To what extent are bistability and oscillation structurally modular in the MMI2 Models? Addressing this question requires the identification of essential molecular species for bistability and oscillation respectively. We focused on two variants of the MMI2 Model that we described earlier: the MMI2-SSB Model where the two binding sites were assumed to be identical (the simplest biological assumption), and the MMI2-ASB Model where only a unique 1:1 complex is possible (a model structurally simpler than the MMI2-SSB Model). We first asked whether the 2:1 complex is required for bistability and oscillation. Removing the reaction responsible for the formation of the 2:1 complex from the MMI2-ASB Model gave rise to the structure of the MMI1 Model which always converges to a point attractor (Fig. 2 and Fig. 7A, Row 1). Removing the same reaction from the MMI2-SSB Model gave rise to a similar model (the C2KO Model in Fig. 7A, Row 2). We found that neither oscillation nor bistability can be obtained with the C2KO Model, i.e. none of the randomly generated 10^5^ parameter sets produced Hopf bifurcation or saddle-node bifurcation. We proceeded to prove this analytically. We found that the C2KO Model has at most one positive steady state, which is asymptotically stable (Section 3.1 in Supplemental Information). As a consequence, the C2KO Model is incapable of generating oscillations or bistability. The 2:1 complex is therefore required for both functions. To test whether a 1:1 complex is required for bistability and oscillation, we assumed that the second binding site is occupied immediately upon the binding of the first site in both MMI2-SSB and MMI2-ASB Models, which gave rise to the C1KO Model (Fig. 7A, bottom right). With this modification, both oscillation and bistability were lost completely (Fig. 7A, Row 3; see Section 3.2 in Supplemental Information for an analytical proof), suggesting that a 1:1 complex is also required to achieve both functions. Because the unbound forms of the mRNA and the miRNA are required to form the complexes, which cannot be produced through transcription directly, it is impossible to remove these unbound species in a biologically meaningful way without removing the complexes. Removing the transcription reaction of either unbound miRNA or unbound mRNA resulted in trivial cases in which the system has a single point attractor (Fig. 7A, Rows 4 and 5). In addition, significant amounts of each of the unbound mRNA species always appeared in some phases of the sustained oscillations (Fig. 3C). Similarly, in each bistable system observed with the MMI2-SSB and MMI2-ASB Models, unbound mRNA and unbound miRNA existed in at least one stable steady state. We therefore concluded that the unbound species are required for both bistability and oscillation. Taken together, we found that there is no structural modularity of bistability and oscillation in the MMI2 Model (*J* = 1, right endpoint of Fig. 7B).

**Figure 7.**
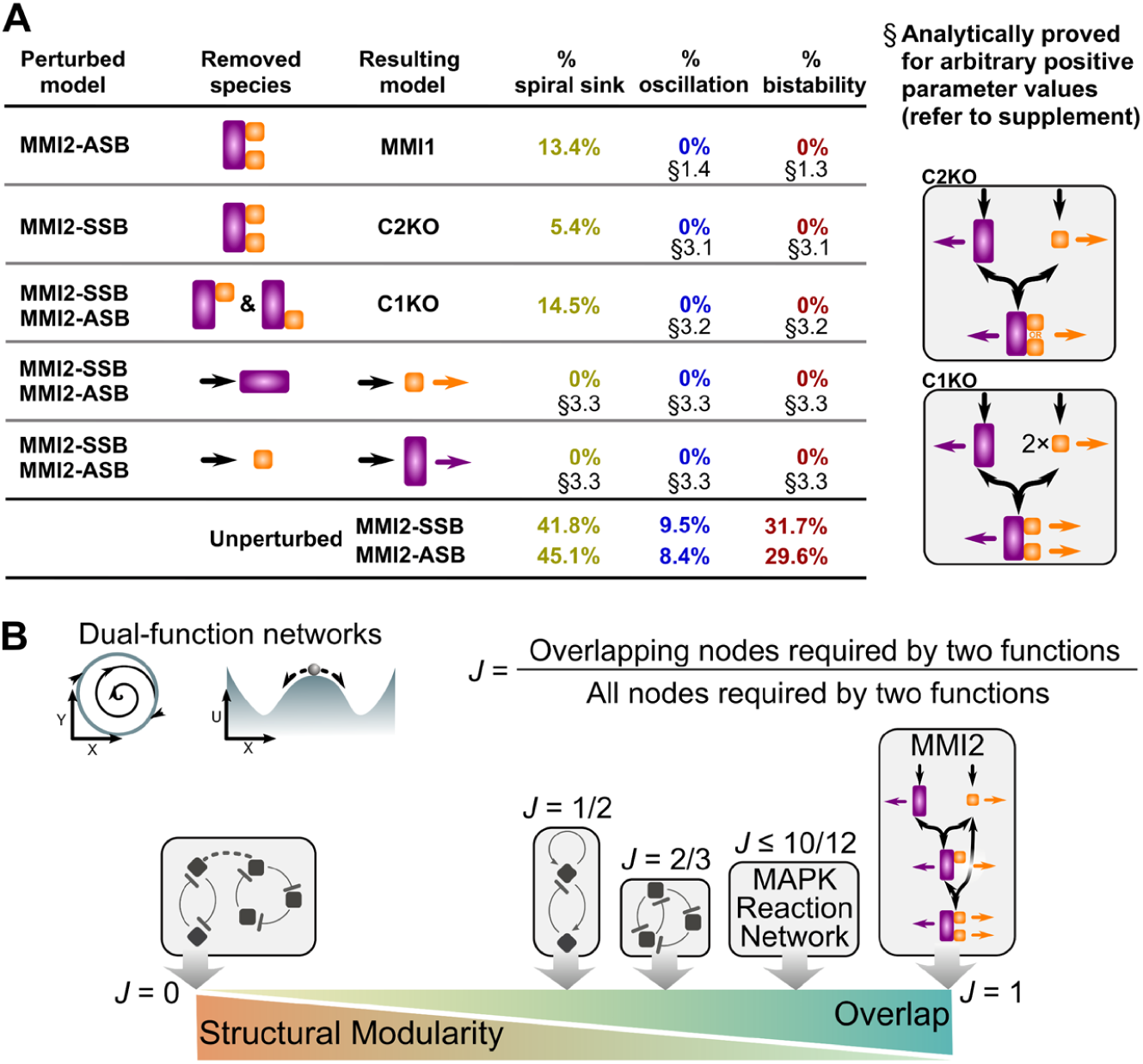
Nonmodular combination of two functions in the MMI2 Models. (**A**) Table shows five biologically plausible ways for removing molecular species from the MMI2-SSB or the MMI2-ASB Models. For each model, 10^5^ parameter sets were randomly generated. Percentages of indicated steady state behaviors through bifurcation analysis with respect to *σ*_*R*_ are shown in the last three columns. In all cases, 0% means that exactly none of the 10^5^ sets produced the indicated type of steady state. Bottom rows show the performance of the unperturbed MMI2-SSB and MMI2-ASB Models. Right diagrams show structures of the two new perturbed models. (**B**) Summary of the structural modularity (quantified by the Jaccard index) of representative networks containing sustained oscillation and bistability. The Jaccard indices of the combined toggle-switch-repressilator circuit and the MAPK Network were estimated based on previous work (Obatake et al., 2019; Perez-Carrasco et al., 2018).

## DISCUSSION

### MicroRNA-mediated gene expression variation

MicroRNA has been extensively studied in its roles in attenuating noise and increasing phenotypic robustness (Li *et al*., 2009). However, recent data suggest that some microRNA can increase gene expression fluctuations and facilitate phenotypic variation (Chakraborty *et al*., 2020; Schmiedel *et al*., 2015). Nonetheless, previous theories of miRNA dynamics assume that microRNA-mRNA interactions alone do not produce oscillatory or excitable systems (Li *et al*., 2021; Osella et al., 2011). Our results raise the possibility that some microRNAs may induce slow-timescale oscillatory dynamics without a transcription-dependent feedback loop. This suggests that the observed variation-amplifying effects of miRNA may be due to limit cycle oscillations or excitability resulting from simple reaction networks of microRNA and mRNA. These destabilizing dynamics at the post-transcriptional level may provide additional strategies for cells to encode and process information, or to regenerate heterogeneity in cell populations.

### Modularity of oscillators and bistable switches

Negative and positive feedback loops have been considered essential components for biological rhythms and switches, respectively (Thomas, 1981; Tyson et al., 2003). Consistent with the difference in structure between the two feedback loops, each of the previously studied biological networks with both switch and rhythm functions contains two distinct modules, which may have partial, but not complete, overlap in terms of molecular species (e.g. gene products) (Gelens et al., 2014; Iglesias and Devreotes, 2012; Liu et al., 2021; Moenke et al., 2017; Obatake *et al*., 2019; Perez-Carrasco *et al*., 2018; Rubinstein et al., 2016). Here, we showed the possibility that the two functions may not have any structural modularity in the RNA-centric reaction networks, which highlights the importance of functional modularity when studying biological networks (Jiménez *et al*., 2017; Verd et al., 2019). Furthermore, this work revealed broad ranges of kinetic rate constants allowing oscillation and bistability in the MMI2 Models, which may guide synthetic biologists to design RNA-based circuits with versatile functions.

### Comparison with other dual oscillator-switches

It was previously shown that the MAPK cascade, a widely studied protein signaling network, produces both oscillation and bistability (Qiao et al., 2007). The periods of the oscillations generated by the protein network (about 30 min) are much shorter than those generated by the MMI2 Models. It was recently found that some molecular species in the MAPK network are required for bistability, but not oscillation (Obatake *et al*., 2019), indicating structural modularity in the network. Nonetheless, because of the highly integrated structural components, we expect that the MAPK network shares some of the emergent properties that we discussed for the RNA-based diverging oscillator (e.g. global bifurcations), and it will be important to investigate the mathematical basis of these oscillator-switches collectively in the future.

Another recent study combined a repressilator and toggle-switch to generate a dual-function circuit based on transcriptional regulations (Perez-Carrasco *et al*., 2018). To achieve both oscillation and bistability in similar parameter regions, the circuit requires an additional feed-forward loop at the signal level. Furthermore, the combined circuit is structurally modular with respect to the two functions (Perez-Carrasco *et al*., 2018). Finally, due to the long distance between the orbit and the point attractors in the state space, global bifurcations are relatively rare with the circuit (Perez-Carrasco *et al*., 2018). All these properties are distinct from those of the diverging oscillator studied here.

With a network containing nine transcriptional regulations, Jutras-Dubé et al. recently showed the superior robustness of SNIC bifurcation in generating embryonic patterns compared to Hopf bifurcation (Jutras-Dubé et al., 2020). The MMI2 Models provide a simple mechanism for generating such global bifurcation at the post-transcriptional level, which could be used for patterning in early development. While it may be possible for a system to achieve dual functions through paradoxical feedback involving one transcription factor (TF) that both activates and inhibits a target gene directly (Kalmar *et al*., 2009; Liu et al., 2020), the paradoxical regulation requires two TF-promoter complexes responsible for activation and inhibition respectively (Perales et al., 2016), which would suggest a fundamental structural modularity.

### Robust regeneration of multimodal gene expression patterns

Spontaneous regeneration of cell populations with multimodal gene expression patterns was observed in progenitor cells and cancer cells, but the mechanism for this phenomenon at slow time scales has been a longstanding problem in biology. While noise-induced transitions between stable steady states (point attractors) have been used to explain some of the observations, it was shown that the modality of the gene expression distribution and its underlying “epigenetic landscape” is very sensitive to the level of noise (Coomer et al., 2021). We showed that oscillations with diverging periods can be useful to regenerate multimodal gene expression. In this mechanism, transitions between very distinct cell states are supported by both deterministic and stochastic dynamics: limit cycles and excitable vector fields ensure the periodic occurrence of dramatic change of gene expression, whereas stochasticity is important for producing the asynchrony in the cell population, and for triggering the oscillatory responses in some cases. The possibility of using excitable systems to generate multimodal expression pattern was discussed previously (Kalmar *et al*., 2009).

As this study examined the dynamics of generalized models, it cannot identify specific networks involved in observed heterogeneity restoration processes. Determining how many real mRNA-microRNA networks fall into the parameter regions responsible for these dynamics would require detailed experimental measurements. The models furthermore do not incorporate impacts on RNA decay or noise levels by other dynamic cellular processes, which may enhance or suppress variation in expression.

Nevertheless, our work showed a simple and potentially widespread mechanism to achieve period-diverging oscillations. This hitherto unknown mechanism can help researchers discover circuits responsible for regenerating heterogenous populations of specific cell types.

## ACKNOWLEDGEMENTS

The authors thank Chunhe Li and John Tyson for helpful discussions. This work was supported by the National Institutes of Health grant R01GM140462 to TH.

## AUTHOR CONTRIBUTIONS

Conceptualization: TH. Formal analysis: PYY and TH. Funding acquisition: TH. Investigation: BN, PYY, GL, and TH. Software: BN and TH. Visualization: BN and TH. Writing—original draft: BN, PYY, and TH.

## METHODS

### Model construction

All models in this study are based on mass-action kinetics. The ODE models were nondimensionalized by scaling the variables and parameters with the degradation rate constant of mRNA, and the synthesis rate constant of the microRNA. One time unit corresponds to approximately 1.*44* × *t*_1/2_ where *t*_1/2_ is the half-life of the mRNA. For the MMI1 and the MMI2-SSB Models, we applied total quasi-steady state assumption (Borghans *et al*., 1996; Ciliberto *et al*., 2007; Kim and Tyson, 2020), and reduced each model to two ODEs. The full models were used in all simulations and bifurcation analyses. The 2-dimensional models were used to construct phase planes and the determinant-trace plot for their Jacobian matrices. The two versions of the model gave consistent results.

### Parameter sampling and numerical bifurcation analysis

To investigate the possible dynamical features of the models, 10^5^ parameter values were randomly selected for each model. The distributions of the parameters were estimated with known ranges of biologically plausible values (Supplemental Information Section 1.5). Synthesis rate constant of mRNA *σ*_*R*_ was used as the control parameter for numerical bifurcation analysis, which quantified the steady state signal-response relationship in the range 0 < *σ*_*R*_ < 25. Local bifurcation points were detected using Tellurium (Choi et al., 2018). Limit cycles were followed using PyDSTool (Clewley, 2012), which further provided the information for global bifurcations. Spiral sinks were detected by identifying complex eigenvalues of Jacobian matrix at the scanned stable steady states numerically.

### Algebraic analysis of the MMI Models

Using the Chemical Reaction Network Toolbox and its underlying theory (Feinberg, 2019), we determined the possibility of obtaining one or more positive steady state for each model. Specifically, the MMI1, C1KO, and C2KO Models cannot admit more than one positive steady state with any positive rate constants, whereas MMI2-SSB, MMI2-ASB and MMI2-DMI can admit two stable positive steady states with some with positive rate constants. The stability of the steady state in the MMI1, the C2KO and C1KO Models were determined by the Routh-Hurwitz Stability Criterion (Hurwitz, 1895; Routh, 1877).

### Estimation of occurrences of the MMI Models

Predicted microRNA-mRNA binding sites from TargetScan were used to determine possible occurrences of the MMI1 and MMI2 Model structures (Agarwal *et al*., 2015). Protein coding genes that have one microRNA binding site were used to estimate the occurrence of the structure of the MMI1 Model: 2554 genes. For the MMI2 Models, we used protein coding genes that have more than one microRNA binding site: 10483 genes. We further used miRTarBase to obtain the number of genes, 12616, that match the MMI2-DMI Model with experimental evidence supporting the microRNA-RNA binding (Huang *et al*., 2020). Each of these genes can be targeted by two microRNA families. Among them, 8420 genes contain more than one microRNA binding sites predicted by TargetScan. Although TargetScan may have many false positives, it is known that false negatives also exist. For example, TargetScan does not include the experimentally validated targeting of *Nanog* by miR-296 at two binding sites (Tay *et al*., 2008).

### Estimation of occurrences of negative transcriptional feedback loops

Negative transcriptional feedback loops were enumerated using HiLoop (Nordick and Hong, 2021) on the full network derived from the TRRUST version 2 database (Han et al., 2018). Limiting loops to at most 3 genes for comparable complexity with the MMI Models, 52 negative feedback loops were found. Of the 2862 genes in the network, 62 were involved in at least one such negative feedback loop.

### Stochastic simulation and quasi-potential landscape

We performed stochastic simulations for the MMI2-SSB Model using either additive or multiplicative noise in the ODEs. To select a cell state with a representative extreme expression pattern, we first solved the ODEs deterministically with 400 initial conditions and selected the state with either the lowest or the highest level of *R*_T_ in the period 40 < *t* < 200. For one simulation of a cell population, we used this state as the initial conditions for 500 cells, and we solved the stochastic ODEs for another 200 time-units (Rackauckas and Nie, 2017). Bimodality was determined by Gaussian-mixture models and Bayesian information criterion. Noise levels and other parameters were changed to examine the robustness of the bimodality regeneration. The simulation results were subsequently used to construct quasi-potential landscapes with potential *U*(*x*) = − log *P*_S_(*x*), where *P*_S_(*x*) is the probability density function at the stationary phase (*t* > 100). To consider intrinsic noise alone in an accurate manner, we implemented an additional form of stochastic simulation based on the Gillespie algorithm and propensity functions derived from reactions shown in Fig. 2A. Using simulations of 200 cells, we examined the consistency between the stochastic ODEs and the Gillespie algorithm under different levels of noise and signals *σ*_*R*_. The stochastic model with additive noise was also extended to capture the dynamics of cell proliferation (Supplemental Information Section 2.1.8).

### Statistical analysis

Bimodality of cell populations was tested by comparing the Bayesian Information Criterion of a one-Gaussian fit of mRNA levels to a two-Gaussian fit. Gaussian mixture models were fit by expectation maximization with GaussianMixtures.jl. The two-Gaussian mixture models were further tested for distinguishability of the modes (Muratov and Gnedin, 2010), requiring 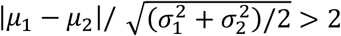.

## Code availability

*Data and code availability:* Computer code for reproducing the results in this study will be publicly available upon completion of peer review.

## SUPPLEMENTAL INFORMATION

**Supplemental Information PDF**. Details of models, mathematical analyses, and numerical methods. Includes Figures S1 to S16 and Tables S1 to S4.

**Movie S1**. Quasi-potential landscapes, dynamics of *R* -high cell fractions and representative trajectories of the diverging oscillator mechanism (left) and the bistable switch mechanism (right) under medium noise condition. Parameter values are listed in Table S1. 50 cells were randomly selected from each 500-cell population. A snapshot of this movie is shown in Figure 6A-B.

**Movie S2**. Quasi-potential landscapes, dynamics of *R* -high cell fractions and representative trajectories of the diverging oscillator mechanism (left) and the bistable switch mechanism (right) under high noise condition. Parameter values are listed in Table S1. 50 cells were randomly selected from each 500-cell population.

## REFERENCES

Abranches, E., Guedes, A.M.V., Moravec, M., Maamar, H., Svoboda, P., Raj, A., and Henrique, D. (2014). Stochastic NANOG fluctuations allow mouse embryonic stem cells to explore pluripotency. Development 141, 2770–2779.

Agarwal, V., Bell, G.W., Nam, J.-W., and Bartel, D.P. (2015). Predicting effective microRNA target sites in mammalian mRNAs. eLife 4, e05005.

Baccarini, A., Chauhan, H., Gardner, T.J., Jayaprakash, A.D., Sachidanandam, R., and Brown, B.D. (2011). Kinetic analysis reveals the fate of a microRNA following target regulation in mammalian cells. Curr. Biol. 21, 369–376.

Bonev, B., Stanley, P., and Papalopulu, N. (2012). MicroRNA-9 modulates Hes1 ultradian oscillations by forming a double-negative feedback loop. Cell reports 2, 10–18.

Borghans, J.A.M., De Boer, R.J., and Segel, L.A. (1996). Extending the quasi-steady state approximation by changing variables. Bull. Math. Biol. 58, 43–63.

Buck, A.H., Coakley, G., Simbari, F., McSorley, H.J., Quintana, J.F., Le Bihan, T., Kumar, S., Abreu-Goodger, C., Lear, M., and Harcus, Y. (2014). Exosomes secreted by nematode parasites transfer small RNAs to mammalian cells and modulate innate immunity. Nat. Commun. 5, 1–12.

Burk, U., Schubert, J., Wellner, U., Schmalhofer, O., Vincan, E., Spaderna, S., and Brabletz, T. (2008). A reciprocal repression between ZEB1 and members of the miR-200 family promotes EMT and invasion in cancer cells. EMBO Rep. 9, 582–589. 10.1038/embor.2008.74.

Chakraborty, M., Hu, S., Visness, E., Del Giudice, M., De Martino, A., Bosia, C., Sharp, P.A., and Garg, S. (2020). MicroRNAs organize intrinsic variation into stem cell states. Proc. Natl. Acad. Sci. USA 117, 6942–6950.

Chang, H.H., Hemberg, M., Barahona, M., Ingber, D.E., and Huang, S. (2008). Transcriptome-wide noise controls lineage choice in mammalian progenitor cells. Nature 453, 544–547. 10.1038/nature06965.

Choi, K., Medley, J.K., König, M., Stocking, K., Smith, L., Gu, S., and Sauro, H.M. (2018). Tellurium: An extensible python-based modeling environment for systems and synthetic biology. Biosystems 171, 74–79.

Ciliberto, A., Capuani, F., and Tyson, J.J. (2007). Modeling networks of coupled enzymatic reactions using the total quasi-steady state approximation. PLoS Comput. Biol. 3, e45.

Clewley, R. (2012). Hybrid Models and Biological Model Reduction with PyDSTool. PLoS Comput. Biol. 8, e1002628. 10.1371/journal.pcbi.1002628.

Coomer, M.A., Ham, L., and Stumpf, M.P.H. (2021). Noise distorts the epigenetic landscape and shapes cell-fate decisions. Cell Systems.

Corrigan, A.M., Tunnacliffe, E., Cannon, D., and Chubb, J.R. (2016). A continuum model of transcriptional bursting. Elife 5, e13051.

Cursons, J., Pillman, K.A., Scheer, K.G., Gregory, P.A., Foroutan, M., Hediyeh-Zadeh, S., Toubia, J., Crampin, E.J., Goodall, G.J., Bracken, C.P., and Davis, M.J. (2018). Combinatorial Targeting by MicroRNAs Co-ordinates Post-transcriptional Control of EMT. Cell Systems 7, 77-91.e77. 10.1016/j.cels.2018.05.019.

de la Mata, M., Gaidatzis, D., Vitanescu, M., Stadler, M.B., Wentzel, C., Scheiffele, P., Filipowicz, W., and Großhans, H. (2015). Potent degradation of neuronal miRNAs induced by highly complementary targets. EMBO Rep. 16, 500–511.

Eichhorn, S.W., Guo, H., McGeary, S.E., Rodriguez-Mias, R.A., Shin, C., Baek, D., Hsu, S.-h., Ghoshal, K., Villén, J., and Bartel, D.P. (2014). mRNA destabilization is the dominant effect of mammalian microRNAs by the time substantial repression ensues. Mol. Cell 56, 104–115.

Elowitz, M.B., and Leibler, S. (2000). A synthetic oscillatory network of transcriptional regulators. Nature 403, 335–338. 10.1038/35002125.

Enver, T., Pera, M., Peterson, C., and Andrews, P.W. (2009). Stem cell states, fates, and the rules of attraction. Cell Stem Cell 4, 387–397.

Ermentrout, B. (1996). Type I membranes, phase resetting curves, and synchrony. Neural Comput. 8, 979–1001.

Feinberg, M. (2019). Foundations of Chemical Reaction Network Theory (Springer International Publishing).

Gao, Q., Zhou, L., Yang, S.-Y., and Cao, J.-M. (2016). A novel role of microRNA 17-5p in the modulation of circadian rhythm. Sci. Rep. 6, 1–12.

Gelens, L., Anderson, G.A., and Ferrell Jr, J.E. (2014). Spatial trigger waves: positive feedback gets you a long way. Mol. Biol. Cell 25, 3486–3493.

Ghini, F., Rubolino, C., Climent, M., Simeone, I., Marzi, M.J., and Nicassio, F. (2018). Endogenous transcripts control miRNA levels and activity in mammalian cells by target-directed miRNA degradation. Nat. Commun. 9, 3119.

Grimson, A., Farh, K.K.-H., Johnston, W.K., Garrett-Engele, P., Lim, L.P., and Bartel, D.P. (2007). MicroRNA targeting specificity in mammals: determinants beyond seed pairing. Mol. Cell 27, 91–105.

Han, H., Cho, J.-W., Lee, S., Yun, A., Kim, H., Bae, D., Yang, S., Kim, C.Y., Lee, M., Kim, E., et al. (2018). TRRUST v2: an expanded reference database of human and mouse transcriptional regulatory interactions. Nucleic Acids Res 46, D380–D386. 10.1093/nar/gkx1013.

Huang, H.-Y., Lin, Y.-C.-D., Li, J., Huang, K.-Y., Shrestha, S., Hong, H.-C., Tang, Y., Chen, Y.-G., Jin, C.-N., and Yu, Y. (2020). miRTarBase 2020: updates to the experimentally validated microRNA–target interaction database. Nucleic Acids Res. 48, D148–D154.

Huang, S. (2009). Non-genetic heterogeneity of cells in development: more than just noise. Development 136, 3853–3862.

Hurwitz, A. (1895). Ueber die Bedingungen, unter welchen eine Gleichung nur Wurzeln mit negativen reellen Theilen besitzt. Mathematische Annalen 46, 273–284.

Iglesias, P.A., and Devreotes, P.N. (2012). Biased excitable networks: how cells direct motion in response to gradients. Curr. Opin. Cell Biol. 24, 245–253.

Jiménez, A., Cotterell, J., Munteanu, A., and Sharpe, J. (2017). A spectrum of modularity in multi-functional gene circuits. Mol. Syst. Biol. 13, 925.

Jordan, N.V., Bardia, A., Wittner, B.S., Benes, C., Ligorio, M., Zheng, Y., Yu, M., Sundaresan, T.K., Licausi, J.A., and Desai, R. (2016). HER2 expression identifies dynamic functional states within circulating breast cancer cells. Nature 537, 102–106.

Jutras-Dubé, L., El-Sherif, E., and François, P. (2020). Geometric models for robust encoding of dynamical information into embryonic patterns. Elife 9, e55778.

Kalmar, T., Lim, C., Hayward, P., Muñoz-Descalzo, S., Nichols, J., Garcia-Ojalvo, J., and Martinez Arias, A. (2009). Regulated fluctuations in nanog expression mediate cell fate decisions in embryonic stem cells. PLoS Biol. 7, e1000149.

Kim, J.K., and Tyson, J.J. (2020). Misuse of the Michaelis–Menten rate law for protein interaction networks and its remedy. PLoS Computational Biology 16, e1008258.

Li, C.-J., Hong, T., Tung, Y.-T., Yen, Y.-P., Hsu, H.-C., Lu, Y.-L., Chang, M., Nie, Q., and Chen, J.-A. (2017). MicroRNA filters Hox temporal transcription noise to confer boundary formation in the spinal cord. Nat. Commun. 8, 14685. 10.1038/ncomms14685.

Li, C., and Wang, J. (2013). Quantifying cell fate decisions for differentiation and reprogramming of a human stem cell network: landscape and biological paths. PLoS computational biology 9, e1003165.

Li, C.J., Liau, E.S., Lee, Y.H., Huang, Y.Z., Liu, Z., Willems, A., Garside, V., McGlinn, E., Chen, J.A., and Hong, T. (2021). MicroRNA governs bistable cell differentiation and lineage segregation via a noncanonical feedback. Mol. Syst. Biol. 17, e9945.

Li, X., Cassidy, J.J., Reinke, C.A., Fischboeck, S., and Carthew, R.W. (2009). A microRNA imparts robustness against environmental fluctuation during development. Cell 137, 273–282.

Liu, Y., Rens, E.G., and Edelstein-Keshet, L. (2021). Spots, stripes, and spiral waves in models for static and motile cells. J. Math. Biol. 82, 1–38.

Liu, Z., Shpak, E.D., and Hong, T. (2020). A mathematical model for understanding synergistic regulations and paradoxical feedbacks in the shoot apical meristem. Computational and Structural Biotechnology Journal 18, 3877–3889.

Ma, F., Xu, S., Liu, X., Zhang, Q., Xu, X., Liu, M., Hua, M., Li, N., Yao, H., and Cao, X. (2011). The microRNA miR-29 controls innate and adaptive immune responses to intracellular bacterial infection by targeting interferon-γ. Nat. Immunol. 12, 861–869.

McGlinn, E., Yekta, S., Mansfield, J.H., Soutschek, J., Bartel, D.P., and Tabin, C.J. (2009). In ovo application of antagomiRs indicates a role for miR-196 in patterning the chick axial skeleton through Hox gene regulation. Proc. Natl. Acad. Sci. USA 106, 18610–18615.

Min, M., and Spencer, S.L. (2019). Spontaneously slow-cycling subpopulations of human cells originate from activation of stress-response pathways. PLoS Biol. 17, e3000178.

Miranda, K.C., Huynh, T., Tay, Y., Ang, Y.-S., Tam, W.-L., Thomson, A.M., Lim, B., and Rigoutsos, I. (2006). A pattern-based method for the identification of MicroRNA binding sites and their corresponding heteroduplexes. Cell 126, 1203–1217.

Moenke, G., Cristiano, E., Finzel, A., Friedrich, D., Herzel, H., Falcke, M., and Loewer, A. (2017). Excitability in the p53 network mediates robust signaling with tunable activation thresholds in single cells. Sci. Rep. 7, 1–14.

Moris, N., Pina, C., and Arias, A.M. (2016). Transition states and cell fate decisions in epigenetic landscapes. Nat Rev Genet 17, 693–703. 10.1038/nrg.2016.98.

Mukherji, S., Ebert, M.S., Zheng, G.X.Y., Tsang, J.S., Sharp, P.A., and van Oudenaarden, A. (2011). MicroRNAs can generate thresholds in target gene expression. Nat. Genet. 43, 854.

Muratov, A.L., and Gnedin, O.Y. (2010). Modeling the Metallicity Distribution of Globular Clusters. The Astrophysical Journal 718, 1266–1288. 10.1088/0004-637x/718/2/1266.

Nordick, B., and Hong, T. (2021). Identification, visualization, statistical analysis and mathematical modeling of high-feedback loops in gene regulatory networks. BMC Bioinformatics 22, 481. 10.1186/s12859-021-04405-z.

Novák, B., and Tyson, J.J. (2008). Design principles of biochemical oscillators. Nature reviews Molecular cell biology 9, 981–991.

Obatake, N., Shiu, A., Tang, X., and Torres, A. (2019). Oscillations and bistability in a model of ERK regulation. J. Math. Biol. 79, 1515–1549.

Osella, M., Bosia, C., Corá, D., and Caselle, M. (2011). The role of incoherent microRNA-mediated feedforward loops in noise buffering. PLoS computational biology 7, e1001101.

Pan, W., Zhu, S., Dai, D., Liu, Z., Li, D., Li, B., Gagliani, N., Zheng, Y., Tang, Y., and Weirauch, M.T. (2015). MiR-125a targets effector programs to stabilize Treg-mediated immune homeostasis. Nat. Commun. 6, 1–12.

Peláez, N., Gavalda-Miralles, A., Wang, B., Navarro, H.T., Gudjonson, H., Rebay, I., Dinner, A.R., Katsaggelos, A.K., Amaral, L.A.N., and Carthew, R.W. (2015). Dynamics and heterogeneity of a fate determinant during transition towards cell differentiation. Elife 4, e08924.

Perales, M., Rodriguez, K., Snipes, S., Yadav, R.K., Diaz-Mendoza, M., and Reddy, G.V. (2016). Threshold-dependent transcriptional discrimination underlies stem cell homeostasis. Proc. Natl. Acad. Sci. USA 113, E6298–E6306.

Perez-Carrasco, R., Barnes, C.P., Schaerli, Y., Isalan, M., Briscoe, J., and Page, K.M. (2018). Combining a toggle switch and a repressilator within the AC-DC circuit generates distinct dynamical behaviors. Cell systems 6, 521–530.

Phillips, N.E., Manning, C.S., Pettini, T., Biga, V., Marinopoulou, E., Stanley, P., Boyd, J., Bagnall, J., Paszek, P., and Spiller, D.G. (2016). Stochasticity in the miR-9/Hes1 oscillatory network can account for clonal heterogeneity in the timing of differentiation. eLife 5, e16118.

Pina, C., Fugazza, C., Tipping, A.J., Brown, J., Soneji, S., Teles, J., Peterson, C., and Enver, T. (2012). Inferring rules of lineage commitment in haematopoiesis. Nat. Cell Biol. 14, 287–294.

Qiao, L., Nachbar, R.B., Kevrekidis, I.G., and Shvartsman, S.Y. (2007). Bistability and oscillations in the Huang-Ferrell model of MAPK signaling. PLoS computational biology 3, e184.

Rackauckas, C., and Nie, Q. (2017). Differentialequations. jl–a performant and feature-rich ecosystem for solving differential equations in julia. Journal of Open Research Software 5.

Raj, A., Peskin, C.S., Tranchina, D., Vargas, D.Y., and Tyagi, S. (2006). Stochastic mRNA synthesis in mammalian cells. PLoS Biol. 4, e309.

Raj, A., and Van Oudenaarden, A. (2008). Nature, nurture, or chance: stochastic gene expression and its consequences. Cell 135, 216–226.

Routh, E.J. (1877). A Treatise on the Stability of a Given State of Motion, Particularly Steady Motion: Being the Essay to which the Adams Prize was Adjudged in 1877, in the University of Cambridge (Macmillan and Company).

Rubinstein, B.Y., Mattingly, H.H., Berezhkovskii, A.M., and Shvartsman, S.Y. (2016). Long-term dynamics of multisite phosphorylation. Mol. Biol. Cell 27, 2331–2340.

Schmiedel, J.M., Klemm, S.L., Zheng, Y., Sahay, A., Blüthgen, N., Marks, D.S., and van Oudenaarden, A. (2015). MicroRNA control of protein expression noise. Science 348, 128–132.

Shaffer, S.M., Dunagin, M.C., Torborg, S.R., Torre, E.A., Emert, B., Krepler, C., Beqiri, M., Sproesser, K., Brafford, P.A., and Xiao, M. (2017). Rare cell variability and drug-induced reprogramming as a mode of cancer drug resistance. Nature 546, 431.

Sharova, L.V., Sharov, A.A., Nedorezov, T., Piao, Y., Shaik, N., and Ko, M.S.H. (2009). Database for mRNA half-life of 19 977 genes obtained by DNA microarray analysis of pluripotent and differentiating mouse embryonic stem cells. DNA Res. 16, 45–58.

Spencer, S.L., Gaudet, S., Albeck, J.G., Burke, J.M., and Sorger, P.K. (2009). Non-genetic origins of cell-to-cell variability in TRAIL-induced apoptosis. Nature 459, 428–432.

Strogatz, S.H. (2000). Nonlinear Dynamics and Chaos: With Applications to Physics, Biology, Chemistry and Engineering (Westview).

Tay, Y., Zhang, J., Thomson, A.M., Lim, B., and Rigoutsos, I. (2008). MicroRNAs to Nanog, Oct4 and Sox2 coding regions modulate embryonic stem cell differentiation. Nature 455, 1124–1128.

Thomas, R. (1981). On the relation between the logical structure of systems and their ability to generate multiple steady states or sustained oscillations. In Numerical methods in the study of critical phenomena, (Springer), pp. 180–193.

Tian, X.J., Zhang, H., Zhang, J., and Xing, J. (2016). Reciprocal regulation between mRNA and microRNA enables a bistable switch that directs cell fate decisions. FEBS Lett. 590, 3443–3455.

Tyson, J.J., Chen, K.C., and Novak, B. (2003). Sniffers, buzzers, toggles and blinkers: dynamics of regulatory and signaling pathways in the cell. Curr. Opin. Cell Biol. 15, 221–231.

Udomlumleart, T., Hu, S., and Garg, S. (2021). Lineages of embryonic stem cells show non-Markovian state transitions. Iscience 24, 102879.

Verd, B., Monk, N.A.M., and Jaeger, J. (2019). Modularity, criticality, and evolvability of a developmental gene regulatory network. Elife 8, e42832.

Wang, F., Zhu, Y., Guo, L., Dong, L., Liu, H., Yin, H., Zhang, Z., Li, Y., Liu, C., and Ma, Y. (2014). A regulatory circuit comprising GATA1/2 switch and microRNA-27a/24 promotes erythropoiesis. Nucleic Acids Res. 42, 442–457.

Wei, L., Li, S., Zhang, P., Hu, T., Zhang, M.Q., Xie, Z., and Wang, X. (2021). Characterizing microRNA-mediated modulation of gene expression noise and its effect on synthetic gene circuits. Cell Reports 36, 109573.

Zhou, Z., Shu, B., Xu, Y., Liu, J., Wang, P., Chen, L., Zhao, J., Liu, X., Qi, S., and Xiong, K. (2018). microRNA-203 modulates wound healing and scar formation via suppressing Hes1 expression in epidermal stem cells. Cell. Physiol. Biochem. 49, 2333–2347.

Zlotorynski, E. (2019). Insights into the kinetics of microRNA biogenesis and turnover. Nature Reviews Molecular Cell Biology 20, 511–511.

