## Supplemental Information for "Nonmodular oscillator and switch based on RNA decay drive regeneration of multimodal gene expression"

Contents

1. The MMI1 Model ... 3
1.1 Model description and nondimensionalization ... 3
1.2 Reduction of the MMI1 Model to a 2-dimensional system ... 4
1.3 The absence of multistability in the MMI1 Model ... 5
1.4 The absence of oscillations in the MMI1 Model ... 6
1.5 Estimation of biologically meaningful rate constants and parameter sampling procedure ... 7
1.5.1 Estimation of basal degradation rate constants ... 7
1.5.2 Estimation of association, dissociation, and microRNA synthesis rate constants ... 8
1.5.3 Estimation of RDFs ... 8
1.5.4 Numerical sampling and bifurcation analysis ... 9
2. The MMI2 Models ... 11
2.1 The MMI2 Model with sequentially symmetrical binding (MMI2-SSB) ... 11
2.1.1 Construction of the MMI2-SSB Model ... 11
2.1.2 Reduction of the MMI2-SSB Model to a 2-dimensional system ... 12
2.1.3 Numerical sampling for the MMI2-SSB Model ... 13
2.1.4 Sequentially symmetrical binding of a coregulated microRNA ... 15
2.1.5 Stochastic simulation of the MMI2-SSB Model ... 15
2.1.6 Quasi-potential landscapes ... 17
2.1.7 Robustness and speed of heterogeneity restoration ... 20
2.1.8 Stochastic simulation with cell division ... 27
2.2 The MMI2 Model with sequentially asymmetrical binding (MMI2-ASB) ... 28
2.3 The MMI2 Model with dual microRNAs (MMI2-DMI) ... 29
3. The structurally perturbed MMI2 Models ... 30
3.1 The C2-Knockout (C2KO) Model ... 30
3.2 The C1-Knockout (C1KO) Model ... 32
3.3 Other structurally perturbed models ... 33
3.4 Summary of dynamical features of all MMI models ... 33

### 1. The MMI1 Model

#### *1.1 Model description and nondimensionalization*

The eight chemical reactions for the MMI1 Model are

$$\xrightarrow{s_R} \hat{R} \qquad \hat{R} \xrightarrow{k_R^0} \emptyset \qquad \xrightarrow{s_r} \hat{r} \qquad \hat{r} \xrightarrow{k_r^0} \emptyset \qquad \hat{R} + \hat{r} \xrightarrow{k^{\text{on}}} \hat{C} \qquad \hat{C} \xrightarrow{k^{\text{off}}} \hat{R} + \hat{r} \qquad \hat{C} \xrightarrow{k_r^1} \hat{R} \qquad \hat{C} \xrightarrow{k_R^1} \hat{r}. \quad (1)$$

Here, $\hat{R}$, $\hat{r}$ and $\hat{C}$ represent both the molecular species and the corresponding concentrations of unbound mRNA, unbound microRNA, and mRNA-microRNA complex respectively. $s_R$ is the transcription rate constant of mRNA. $s_r$ is the transcription rate constant of microRNA. $k_R^0$ is the degradation rate constant of the unbound mRNA. $k_R^1$ is the degradation rate constant of the mRNA in the complex. $k_r^0$ is the degradation rate constant of the unbound microRNA. $k_r^1$ is the degradation rate constant of the microRNA in the complex. $\kappa^{\text{on}}$ is the association rate constant. $\kappa^{\text{off}}$ is the dissociation rate constant. With the law of mass action, we use the following ordinary differential equations (ODEs) to describe the dynamics of the three species

$$\begin{aligned} d\hat{R}/d\hat{t} &= s_R - k^{\text{on}}\hat{R}\hat{r} + k^{\text{off}}\hat{C} - k_R^0\hat{R} + k_r^1\hat{C} \\ d\hat{r}/d\hat{t} &= s_r - k^{\text{on}}\hat{R}\hat{r} + k^{\text{off}}\hat{C} - k_r^0 r + k_R^1\hat{C} \qquad (2) \\ d\hat{C}/d\hat{t} &= k^{\text{on}}\hat{R}\hat{r} - k^{\text{off}}\hat{C} - k_R^1\hat{C} - k_r^1\hat{C}. \end{aligned}$$

We make the following changes of variables to nondimensionalize the model:

$$\hat{t} = t/k_R^0, \qquad \hat{R} = Rs_r/k_R^0, \qquad \hat{r} = rs_r/k_R^0, \qquad \hat{C} = Cs_r/k_R^0, \qquad (3)$$

which yields

$$\begin{aligned} dR/dt &= \sigma_R - \kappa^{\text{on}}Rr + \kappa^{\text{off}}C - R + \beta\gamma C \\ dr/dt &= 1 - \kappa^{\text{on}}Rr + \kappa^{\text{off}}C - \gamma r + \alpha C \qquad (4) \\ dC/dt &= \kappa^{\text{on}}Rr - \kappa^{\text{off}}C - aC - \beta\gamma C. \end{aligned}$$

In these ODEs, Greek letters are scaled parameters that have the following relationships with the original parameters in Eq 1:

$$\sigma_R = s_R/s_r, \quad \kappa^{\text{on}} = k^{\text{on}}s_r/{k_R^0}^2, \quad \kappa^{\text{off}} = k^{\text{off}}/k_R^0, \quad \gamma = k_r^0/k_R^0, \quad \alpha = k_R^1/k_R^0, \quad \beta = k_r^1/k_r^0. \quad (5)$$

Here, $\sigma_R$ represents the synthesis rate constant of the second mRNA relative to that of the microRNA. $\gamma$ represents the degradation rate constant of the unbound microRNA relative to that of the unbound mRNA. We define $\alpha$ and $\beta$ relative degradation factors (RDFs), and they represent the degradation rate constants of the mRNA and microRNA in the complex relative to those of the unbound forms of the same molecules, respectively.

#### *1.2 Reduction of the MMI1 Model to a 2-dimensional system*

Eq 4 was used in our numerical simulations for the MMI1 Model. However, we also considered another form of the model which only has two ODEs. Using the total-quasi-steady-state assumption (tQSSA), Eq 4 can be approximated by the following differential-algebraic equations (DAEs):

$$dR_{\mathrm{T}}/dt = \sigma_R - (R_{\mathrm{T}} - C) - \alpha C \tag{6a}$$

$$dr_{\mathrm{T}}/dt = 1 - \gamma(r_{\mathrm{T}} - C - \beta C) \tag{6b}$$

$$KC = (R_{\mathrm{T}} - C)(r_{\mathrm{T}} - C). \tag{6c}$$

Here, $K = \kappa^{\mathrm{off}}/\kappa^{\mathrm{on}}$ ($K$ is Kappa), and it is related to the dissociation constant $k^{\mathrm{off}}/k^{\mathrm{on}}$. $R_{\mathrm{T}}$ and $r_{\mathrm{T}}$ represent the total scaled concentrations of the mRNA and the microRNA respectively. Eq 6c represents the binding/unbinding kinetics that is significantly faster than synthesis and degradation of the molecules. With this equation, $C$ can be solved in terms of $R_{\mathrm{T}}$ and $r_{\mathrm{T}}$

$$C = \frac{R_{\mathrm{T}} + r_{\mathrm{T}} + K \pm A}{2}, \tag{7}$$

where

$$A = \sqrt{(R_{\mathrm{T}} - r_{\mathrm{T}} - K)^2 + 4KR_{\mathrm{T}}} = \sqrt{(R_{\mathrm{T}} + r_{\mathrm{T}} + K)^2 - 4R_{\mathrm{T}}r_{\mathrm{T}}}. \tag{8}$$

Because all parameters and variables are positive, this shows that

$$|R_{\mathrm{T}} + r_{\mathrm{T}} + K| > A > |R_{\mathrm{T}} - r_{\mathrm{T}} - K| > 0. \tag{9}$$

Our goal is to show that only one solution shown in Eq 7 is biologically relevant with all positive parameter values and concentrations, even though both solutions appear to be positive. The concentration of the unbound mRNA is

$$R = R_{\mathrm{T}} - C, \tag{10}$$

and $R > 0$ is a biological constraint. Suppose

$$C = \frac{R_{\mathrm{T}} + r_{\mathrm{T}} + K + A}{2}, \tag{11}$$

then

$$R = \frac{R_{\mathrm{T}} - r_{\mathrm{T}} - K - A}{2}. \tag{12}$$

If $R_{\mathrm{T}} - r_{\mathrm{T}} - K < 0$, then there is no positive $R$ that can satisfy the quasi-steady-state condition and conservation relationship shown in Eq 11 and Eq 12 since $A > 0$. Therefore $KR_{\mathrm{T}} - Kr_{\mathrm{T}} - 1 > 0$ in order for Eq 11 to be satisfied under biologically relevant conditions. It follows this conclusion and Eq 9 that

$$A > R_{\mathrm{T}} - r_{\mathrm{T}} - K > 0. \tag{13}$$

Therefore, $R < 0$ for all positive parameter values if Eq 11 is satisfied. Similarly, we can show that if

$$C = \frac{R_{\mathrm{T}} + Kr_{\mathrm{T}} + K - A}{2}, \tag{14}$$

then $R > 0$ for all biologically relevant conditions. We conclude that Eq 14 is always the only biologically meaningful solution to Eq 6c. In fact, based on Eq 9 we can show that $C > 0$ for all positive parameter values. With the substitution shown in Eq 14, Eq 6 can be rewritten as

$$dR_{\mathrm{T}}/dt = f(R_{\mathrm{T}}, r_{\mathrm{T}}) = \sigma_R - \frac{\alpha(R_{\mathrm{T}} + r_{\mathrm{T}} + K - A) + R_{\mathrm{T}} - r_{\mathrm{T}} - K + A}{2} \tag{15a}$$

$$dr_{\mathrm{T}}/dt = g(R_{\mathrm{T}}, r_{\mathrm{T}}) = 1 - \frac{\beta\gamma(R_{\mathrm{T}} + r_{\mathrm{T}} + K - A) - \gamma(R_{\mathrm{T}} - r_{\mathrm{T}} + K - A)}{2} \tag{15b}$$

where $A = \sqrt{(R_{\mathrm{T}} - r_{\mathrm{T}} - K)^2 + 4KR_{\mathrm{T}}}$. Together with the readily derived Jacobian matrix

$$J = \begin{pmatrix} \partial f/\partial R_{\mathrm{T}} & \partial f/\partial r_{\mathrm{T}} \\ \partial g/\partial R_{\mathrm{T}} & \partial g/\partial r_{\mathrm{T}} \end{pmatrix} \tag{16}$$

(full expression not shown here), Eq 15 is used to construct phase planes and to illustrate the stability of the steady state in the trace-determinant space of the Jacobian matrix (Fig. S1).

#### *1.3 The absence of multistability in the MMI1 Model*

Using Deficiency One Theorem under the Chemical Reaction Network Theory (CRNT), it can be shown that the MMI1 Model can only admit one positive steady state (Feinberg, 1988). To use the theorem, we first calculate the number of sets of reactants and products based on Eq 1, and we obtain this number $n = 5$. We next construct the stochiometric subspace for species $\{R, r, C\}$ (equivalently for unscaled species $\{\hat{R}, \hat{r}, \hat{C}\}$) based on reactions shown in Eq 1:

$$S = \mathrm{span}\left\{ \begin{bmatrix}1\\0\\0\end{bmatrix}, \begin{bmatrix}-1\\0\\0\end{bmatrix}, \begin{bmatrix}0\\1\\0\end{bmatrix}, \begin{bmatrix}0\\-1\\0\end{bmatrix}, \begin{bmatrix}-1\\-1\\1\end{bmatrix}, \begin{bmatrix}1\\1\\-1\end{bmatrix}, \begin{bmatrix}1\\0\\-1\end{bmatrix}, \begin{bmatrix}0\\1\\-1\end{bmatrix} \right\}$$

$$= \mathrm{span}\left\{ \begin{bmatrix}1\\0\\0\end{bmatrix}, \begin{bmatrix}-1\\-1\\1\end{bmatrix}, \begin{bmatrix}1\\0\\-1\end{bmatrix} \right\}. \tag{17}$$

The rank of the stochiometric subspace $s = 3$. Because all reaction and product sets are connected, the number of linkage classes $l = 1$. The deficiency of the network is therefore given by $\delta = n - s - l = 1$. It follows the Deficiency One Theorem that we have proved the following statement.

The mass action system described by Eq 4 (or equivalently Eq 2) has a single positive steady state. It is therefore impossible for the MMI1 Model to be bistable for any set of positive rate constants.

#### *1.4 The absence of oscillations in the MMI1 Model*

The conclusion in Section 1.3 obtained with Deficiency One Theorem does not exclude the possibility that the MMI1 Model has an unstable steady state, which is a necessary condition for a stable limit cycle via Hopf bifurcation. The goal of this section is to show that the only positive steady state for the MMI1 Model is stable for all positive rate constants.

To analyze the local stability of a steady state, we check where in the complex plane the eigenvalues of the Jacobian matrix lie. If all the eigenvalues have negative real parts, then the steady state is asymptotically stable.

One method to determine the location of eigenvalues with respect to the imaginary axis is by the Routh-Hurwitz stability criterion (Bodson, 2020; Gantmacher and Brenner, 2005; Meinsma, 1995).

**Theorem (Routh-Hurwitz stability criterion):** Consider the polynomial $p(\lambda) = a_n\lambda^n + a_{n-1}\lambda^{n-1} + \cdots + a_1\lambda + a_0$ with $a_0 \neq 0$ and $a_n > 0$. Construct the following table with $n+1$ rows:

$$\begin{matrix} a_n & a_{n-2} & a_{n-4} & \cdots \\ a_{n-1} & a_{n-3} & a_{n-5} & \cdots \\ r_{3,1} & r_{3,2} & r_{3,3} & \cdots \\ r_{4,1} & r_{4,2} & \cdots & \cdots \\ \vdots & \vdots & \vdots & \ddots \\ r_{n+1,1} & & & \end{matrix}$$

where $r_{1,k}, r_{2,k}$ denote elements of the first two rows, and $r_{i,k} = \frac{-1}{r_{i-1,1}} \det \begin{bmatrix} r_{i-2,1} & r_{i-2,k+1} \\ r_{i-1,1} & r_{i-1,k+1} \end{bmatrix}$ for any $i \geq 3$ and $k \geq 1$. Suppose $r_{i,1} \neq 0$ for all $i \geq 1$. Then no roots of $p(\lambda)$ lie on the imaginary axis. Moreover, the number of roots with positive real parts is equal to the number of sign changes in the first column in the table.

Although the Routh-Hurwitz stability criterion holds for real polynomials of any degree, we only make use of the degree 3 case here (and the degree 4 case in Section 3.1).

**Theorem (Routh-Hurwitz stability criterion for degree 3 polynomials):** Consider the polynomial $p(\lambda) = a_3\lambda^3 + a_2\lambda^2 + a_1\lambda + a_0$ with $a_0 \neq 0$ and $a_3 > 0$. Build the table with four rows:

$$\begin{matrix} a_3 & a_1 \\ a_2 & a_0 \\ u & 0 \\ a_0 & 0 \end{matrix}$$

where $u = \frac{a_2a_1 - a_3a_0}{a_2}$. All the roots of $p(\lambda)$ have negative real parts if and only if $a_2, a_0 > 0$ and $a_2a_1 - a_3a_0 > 0$.

The following expressions are obtained using Mathematica (MMI1.nb). The ODE system in Eq 4 has the Jacobian matrix

$$J = \begin{bmatrix} -1 - \kappa^{\text{on}}r & -\kappa^{\text{on}}R & \kappa^{\text{off}} + \beta\gamma \\ -\kappa^{\text{on}}r & -\kappa^{\text{on}}R - \gamma & \kappa^{\text{off}} + \alpha \\ \kappa^{\text{on}}r & \kappa^{\text{on}}R & -\kappa^{\text{off}} - \beta\gamma - \alpha \end{bmatrix}, \tag{18}$$

whose characteristic polynomial $p(\lambda) = \det(\lambda - J)$ is a degree-3 polynomial. We found that $p(\lambda) = \lambda^3 + a_2\lambda^2 + a_1\lambda + a_0$, where

$$a_2 = 1 + \kappa^{\text{off}} + \kappa^{\text{on}}(r + R) + \alpha + \gamma + \beta\gamma,$$
$$a_1 = \kappa^{\text{off}} + \gamma\kappa^{\text{off}} + \kappa^{\text{on}}R + \alpha\kappa^{\text{on}}r + \gamma\kappa^{\text{on}}r + \beta\gamma\kappa^{\text{on}}R + \beta\gamma + \beta\gamma^2 + \alpha\gamma + \gamma + \alpha, \quad (19)$$
$$a_0 = \kappa^{\text{off}}\gamma + \alpha\gamma\kappa^{\text{on}}r + \beta\gamma\kappa^{\text{on}}R + \alpha\gamma + \beta\gamma^2.$$

It is clear from the expression that for any positive rate constants, with steady state values for $R$ and $r$, we always have $a_0, a_2 > 0$. Lastly, to check the inequality $u > 0$, we extracted the coefficients of $u$, as a polynomial of $\kappa^{\text{off}}, \kappa^{\text{on}}, \alpha, \beta, \gamma, R, r$. The minimum non-zero coefficient is 1, i.e., there are no terms in $u$ with negative coefficients, so $u > 0$.

By the Routh-Hurwitz stability criterion, we conclude that for any positive rate constants, any eigenvalue of the system has negative real part, and the unique positive steady state is asymptotically stable.

#### *1.5 Estimation of biologically meaningful rate constants and parameter sampling procedure*

The analytical results for the MMI1 Model do not exclude the possibility that transient oscillations and associated "spiral sink" steady states can be generated by the model. Furthermore, algebraic analysis for other models described in later sections is generally not feasible. We therefore performed parameter sampling in biologically plausible ranges and examined the models using numerical solutions to the ODEs (e.g. Eq 4 for the MMI1 Model) and bifurcation analysis with respect to $\sigma_R$, the scaled synthesis rate constant of the mRNA.

For each scaled parameter, we first estimated the median value based on reported experimental measurements. We then estimated a biologically reasonable range covering at least two orders of magnitude to consider biological variabilities and measurement errors. This procedure was performed for all parameters except the RDFs, for which we directly estimated the ranges.

##### *1.5.1 Estimation of basal degradation rate constants*

The median mammalian mRNA half-life without post-transcriptional control was estimated to be 4 hours (Sharova et al., 2009). We therefore estimated the median $\tilde{k}_R^0 = \ln(2)/4\ \text{hr}^{-1} \approx 0.17\ \text{hr}^{-1}$. Because this parameter does not appear in the nondimensionalized ODEs, we did not consider its variation explicitly during parameter sampling. However, it is important to consider its plausible range in biology when interpreting the time-course simulations because the independent variable (time $\hat{t} = t/k_R^0$) was scaled with $k_R^0$. Based on this scaling procedure for time, our estimated $\tilde{k}_R^0$ implies that one time unit in our simulations corresponds to 5.88 hr. The general relationship between the clock time that one time unit in models represents ($\tau$) and the half-life of the mRNA ($t_{1/2}$) is $\tau = t_{1/2}/\ln(2)$.

A recent study reported a short mean half-life, 5 min, of mRNAs in yeast (Chan et al., 2018). Our main interest is to study animal and plant cells where microRNAs are present, but we still consider this measurement for the upper bound of $k_R^0$, and this would imply that one time unit in our simulations corresponds to 7.14 min.

The estimated median half-life $\tilde{k}_R^0$ does not include a minor population (a second mode of the distribution in Ref (Sharova *et al.*, 2009)) of stable mammalian mRNAs whose half-lives are 24 hr or longer. If we use

24 hr as an upper bound of half-life, then one time unit in our simulations corresponds to 34.6 hr as the upper bound. An older study reported that the mean half-life of human mRNAs is 10 hr (Yang et al., 2003), which would imply that one time unit in our simulations corresponds to 14.3 hr.

In conclusion, one time unit in our simulations corresponds approximately 5.88 hr, but it can vary in the range of 7.14 min for rare unstable mRNAs in animals and plants to 34.6 hr for relatively stable mRNAs.

It was estimated that the half-lives of microRNA are approximately four times of the mRNA half-lives (Reichholf et al., 2019; Zlotorynski, 2019). We therefore used this value as the median for $\gamma$, i.e. $\tilde{\gamma} = 1/4$. It has been shown that microRNA half-lives can vary from $\approx 4$ hr to $> 48$ hr (Marzi et al., 2016), suggesting that it is possible that some microRNAs may have shorter half-lives than their targets. We therefore sampled $\gamma$ in the interval $[1/40, 5/2]$ with a log-uniform distribution.

*1.5.2 Estimation of association, dissociation, and microRNA synthesis rate constants*

Similar to $k_R^0$, the microRNA synthesis rate constant $s_r$ is a scaling factor that does not appear in the nondimensionalized ODEs. To relate the scaled variables and parameters to realistic biological quantities, we considered a representative value of $s_r$ for a relatively abundant microRNA. We first estimated the cytoplasmic volume of a murine myoblast to be $V = 1.8 \times 10^{-12}$ L ($1700 \mu\text{M}^3$) (Gingras et al., 2009; Moore et al., 2019). The mean number of two highly expressed microRNA in this cell type under a physiological condition was estimated to be $n = 9.7 \times 10^3$ copies per cell (Pinzón et al., 2017). We therefore estimated that the molar concentration of a highly expressed microRNA is $\tilde{r} = n/N_A/V \approx 9.5 \times 10^{-9}\text{M}$, where $N_A$ is the Avogadro constant. Assuming that this microRNA has a degradation rate constant $\tilde{k}_r^0 = \ln(2)/16\ \text{hr}^{-1} \approx 4.3 \times 10^{-2}\ \text{hr}^{-1}$ (Reichholf *et al.*, 2019), the estimated synthesis rate constant $\tilde{s}_r = \tilde{k}_r^0 \tilde{r} \approx 4.1 \times 10^{-10}\text{M hr}^{-1}$. The dissociation constant of microRNA was estimated to be 3.7 pM (Wee et al., 2012). The scaled dissociation constant $K$ was estimated with $\tilde{K} = (\kappa^{\text{off}}/\kappa^{\text{on}})\,(\tilde{k}_R^0/\tilde{s}_r) = 3.7 \times 10^{-12}\text{M} \times 0.17\ \text{hr}^{-1}/(4.1 \times 10^{-10}\text{M hr}^{-1}) \approx 1.5 \times 10^{-3}$. Because there are significant variations in cell volume, RNA half-lives, microRNA concentration, and binding affinity, we sampled $K$ from a range covering four orders of magnitudes, i.e $[1.5 \times 10^{-5}, 1.5 \times 10^{-1}]$, with a log-uniform distribution to account for the biological variation and measurement errors. This wide range effectively encompasses the possible scenario in which about 10 copies of microRNA are available for the reactions in a cell, due to either low total concentration or existing occupancy by other mRNA targets. To estimate $\kappa^{\text{on}}$, we first took the diffusion limit for $k^{\text{on}}$, i.e. $\tilde{k}^{\text{on}} = 3.6 \times 10^{11}\text{M}^{-1}\text{hr}^{-1}$ (Jarmoskaite et al., 2020). The scaled association rate constant $\tilde{\kappa}^{\text{on}} = \tilde{k}^{\text{on}}\tilde{s}_r/{\tilde{k}_R^0}^2 \approx 5.1 \times 10^3$. $\kappa^{\text{off}}$ was subsequently estimated based on this representative value of $\kappa^{\text{on}}$ and the estimated range of $K$ mentioned above.

*1.5.3 Estimation of RDFs*

The RDFs (e.g. $\alpha$ and $\beta$ for the MMI1 Model) were sampled in the interval [1/8, 16] which was estimated based on previous experimental data (de la Mata et al., 2015; Eichhorn et al., 2014). In most biological scenarios, the RDFs for mRNA (e.g. $\alpha$) are greater than 1, and we focused on this region in all of our discussion. Nonetheless, in some rare cases it is possible that microRNA can decrease the degradation rate of mRNA upon binding (Vasudevan et al., 2007), so we did not exclude the relevant RDF values. We used a log-uniform distribution for random sampling.

*1.5.4 Numerical sampling and bifurcation analysis*

$10^5$ biologically plausible parameter values were randomly drawn from the distributions described above. For each parameter set, numerical continuation was performed with respect to signal $\sigma_R$. They category of the steady state was determined by Jacobian matrix for Eq 4, or equivalently that for Eq 6. The 2D Jacobian matrix can be further analyzed graphically using a determinant-trace plot. For example, the two representative bifurcation curves shown in Fig. 2B (black and gray) are shown in the determinant-trace space in Fig. S1.

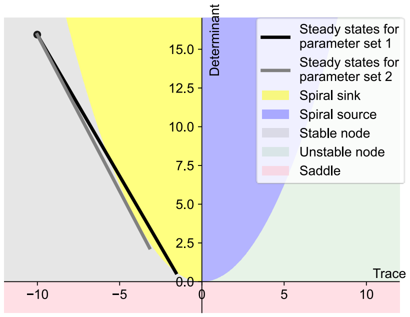

**Fig. S1. Two representative steady state curves projected on to determinant-trace space of the Jacobian matrix.** The representative parameter sets' steady state signal-response curves with respect to $\sigma_R$ were plotted in Fig. 2B. The curves were projected on to the determinant-trace space of Jacobian matrix of the reduced model (Eq 6). Colored areas show categories of steady states. Two curves show the steady states of the system starting from $\sigma_R = 0$ (black dot) to $\sigma_R = 20$. The black starts from a stable node, changes to a spiral sink, and then switches back to a stable node.

Each signal-response curve obtained from the MMI1 Model had a threshold for increase of unbound mRNA concentration (Fig. S2A). The spiral sinks were primarily observed near the threshold $\hat{\sigma}_R$ (Fig. S2A). Furthermore, they gave rise to abrupt changes of unbound mRNA concentration at $\hat{\sigma}_R$. The changes of the mRNA total concentration were relatively moderate (Fig. S2B).

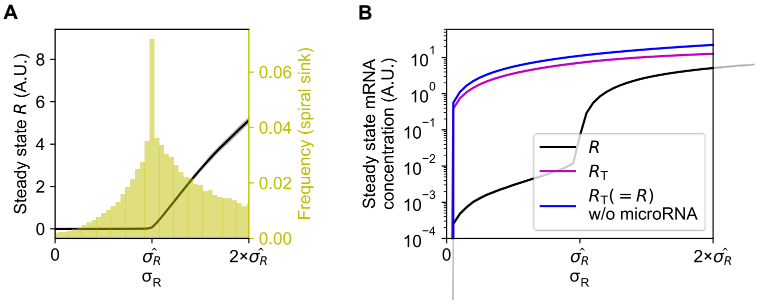

**Fig. S2. Mean response curves of all sampled parameter sets**. **(A)** Distribution of parameter sets with spiral sink steady state (yellow) and those without (gray) among the $10^5$ tested parameter sets. **(B)** Mean response curves with respect to $\sigma_R$ for all $10^5$ parameter sets are shown in black (unbound mRNA) and purple (total mRNA). The curves were first centered at $\hat{\sigma}_R$, where the response $R = 0.01$. A microRNA-free curve is shown in blue as a control response.

We observed that the spiral sink steady states were distributed widely in individual parameters except for $\sigma_R$ (Fig. 2D). However, they require negatively correlated $\alpha$ and $\beta$, as well as positively correlated $\alpha$ and $\gamma$ (Fig. S3). $\gamma$ had a positive effect in generating spiral sinks (Fig. S3).

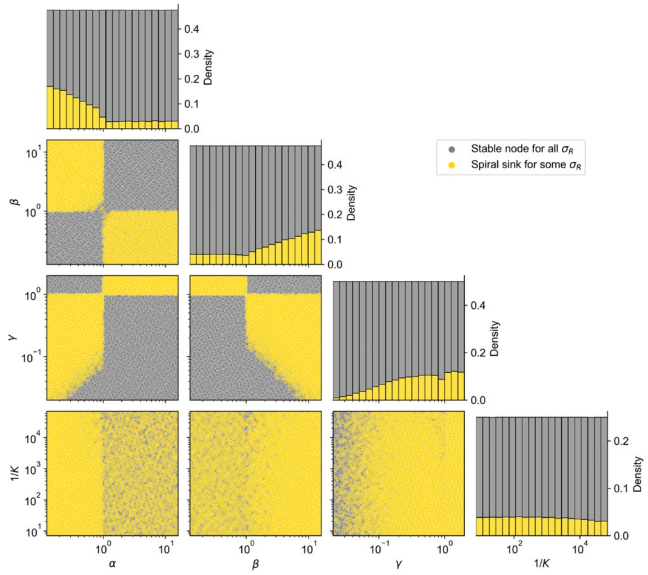

**Fig. S3. Distributions of two types of steady states over individual parameters.** Scatter plots show distributions of $10^5$ random parameter sets categorized by whether they produced spiral sink steady states for some values of $\sigma_R$. Marginal distributions are shown in stacked bars.

### 2. The MMI2 Models

#### *2.1 The MMI2 Model with sequentially symmetrical binding (MMI2-SSB)*

*2.1.1 Construction of the MMI2-SSB Model*

The twelve chemical reactions for the MMI2-SSB Model are

$$\xrightarrow{s_R} \hat{R} \qquad \hat{R} \xrightarrow{k_R^0} \emptyset \qquad \xrightarrow{s_r} \hat{r} \qquad \hat{r} \xrightarrow{k_r^0} \emptyset \qquad \hat{R} + \hat{r} \xrightarrow{k^{\text{on}}} \hat{C}_1 \qquad \hat{C}_1 \xrightarrow{k^{\text{off}}} \hat{R} + \hat{r} \qquad \hat{C}_1 \xrightarrow{k_r^1} \hat{R} \qquad \hat{C}_1 \xrightarrow{k_R^1} \hat{r}$$

$$\hat{C}_1 + \hat{r} \xrightarrow{k^{\text{on}}} \hat{C}_2 \qquad \hat{C}_2 \xrightarrow{k^{\text{off}}} \hat{C}_1 + \hat{r} \qquad \hat{C}_2 \xrightarrow{k_r^1} \hat{C}_1 \qquad C_2 \xrightarrow{k_R^2} 2r \qquad (20)$$

Here, the definitions of the variables and parameters are the same as those for the MMI1 Model except the following: $\hat{C}_1$ and $\hat{C}_2$ are the names (or the concentrations) of a 1:1 complex and the 2:1 complex respectively. Because the mRNA has two binding sites of the microRNA, which are assumed to be identical, $\hat{C}_1$ represents either form of the 1:1 complex and its concentration. The total concentration of 1:1 complex is $2\hat{C}_1$. $k_R^1$ and $k_R^2$ are the degradation rate constants of the mRNA in the 1:1 complex and the 2:1 complex respectively. $k_r^1$ and $k_r^2$ are the degradation rate constants of the microRNA in the 1:1 complex and in the 2:1 complex respectively. With the law of mass action and nondimensionalization, we use the following ordinary differential equations (ODEs) to describe the dynamics of the four species

$$dR/dt = f(R, r, C_1, C_2) = \sigma_R - 2\kappa^{\text{on}} Rr + 2\kappa^{\text{off}} C_1 - R + 2\beta_1 \gamma C_1$$

$$dr/dt = g(R, r, C_1, C_2) = 1 - 2\kappa^{\text{on}} Rr + 2\kappa^{\text{off}} C_1 - 2\kappa^{\text{on}} C_1 r + 2\kappa^{\text{off}} C_2 - \gamma r + 2\alpha_1 C_1 + 2\alpha_2 C_2$$

$$dC_1/dt = h(R, r, C_1, C_2) = \kappa^{\text{on}} Rr - \kappa^{\text{off}} C_1 - \kappa^{\text{on}} C_1 r + \kappa^{\text{off}} C_2 - \alpha_1 C_1 - \beta_1 \gamma C_1 + \beta_2 \gamma C_2 \qquad (21)$$

$$dC_2/dt = k(R, r, C_1, C_2) = 2\kappa^{\text{on}} C_1 r - 2\kappa^{\text{off}} C_2 - \alpha_2 C_2 - 2\beta_2 \gamma C_2.$$

The following changes are made to relate the dimensionless variables and parameters to the originals:

$$\hat{t} = t/k_R^0, \qquad \hat{R} = R s_r/k_R^0, \qquad \hat{r} = r s_r/k_R^0, \qquad \hat{C}_1 = C_1 s_r/k_R^0, \qquad \hat{C}_2 = C_2 s_r/k_R^0$$

$$\sigma_R = s_R/s_r, \qquad \kappa^{\text{on}} = k^{\text{on}} s_r/{k_R^0}^2, \qquad \kappa^{\text{off}} = k^{\text{off}}/k_R^0, \qquad \gamma = k_r^0/k_R^0, \qquad (22)$$

$$\alpha_1 = k_R^1/k_R^0, \qquad \alpha_2 = k_R^2/k_R^0, \qquad \beta_1 = k_r^1/k_r^0, \qquad \beta_2 = k_r^2/k_r^0.$$

Note that because of the symmetrically binding assumption, there are two identical 1:1 complexes described by the same variable $C_1$ (or equivalently $\hat{C}_1$). Therefore, all the reaction rates that involve the 1:1 complex and appear in variables $R$, $r$, $C_2$ are multiplied by 2 in Eq 21. We will relax this assumption in Section 2.2.

In the MMI2-SSB Model, there are four RDFs ($\alpha_1$, $\alpha_2$, $\beta_1$ and $\beta_2$). $\alpha_1$ and $\alpha_2$ represent the degradation rate constants of the mRNA in the 1:1 and 2:1 complexes, respectively, relative to its degradation rate constant in the unbound form. $\beta_1$ and $\beta_2$ are the corresponding RDFs for the microRNA. The definitions of all other parameters are the same as those for the MMI1 Model.

*2.1.2 Reduction of the MMI2-SSB Model to a 2-dimensional system*

The system in Eq 21 is used to perform numerical simulations and bifurcation analysis for the MMI2-SSB Model. However, we also considered another form of the model which only has two ODEs. Using the tQSSA, we approximate Eq 21 with the differential algebraic equations

$$dR_{\mathrm{T}}/dt = f_{\mathrm{T}}(R_{\mathrm{T}}, r_{\mathrm{T}}) = \sigma_R - (R_{\mathrm{T}} - 2C_1 - C_2 + 2\alpha_1 C_1 + \alpha_2 C_2) \tag{23a}$$

$$dr_{\mathrm{T}}/dt = g_{\mathrm{T}}(R_{\mathrm{T}}, r_{\mathrm{T}}) = 1 - \gamma(r_{\mathrm{T}} - 2C_1 - 2C_2 + 2\beta_1 C_1 + 2\beta_2 C_2) \tag{23b}$$

$$KC_1 = (R_{\mathrm{T}} - 2C_1 - C_2)(r_{\mathrm{T}} - 2C_1 - 2C_2) \tag{23c}$$

$$KC_2 = C_1(r_{\mathrm{T}} - 2C_1 - 2C_2). \tag{23d}$$

Here, Kappa $K = \kappa^{\mathrm{off}}/\kappa^{\mathrm{on}}$, related to the dissociation constant $k^{\mathrm{off}}/k^{\mathrm{on}}$. $R_{\mathrm{T}}$ and $r_{\mathrm{T}}$ represent the total scaled concentrations of the mRNA and the microRNA respectively.

We first solve Eq 23d for $C_2$:

$$C_2 = \frac{C_1(-2C_1 + r)}{2C_1 + K}. \tag{24}$$

Substituting this into Eq 23c yields

$$0 = \frac{K\left(4{C_1}^2 R_{\mathrm{T}} + K^2 C_1 + 2KC_1 R_{\mathrm{T}} + 2KC_1 r_{\mathrm{T}} - 2C_1 r_{\mathrm{T}} R_{\mathrm{T}} + C_1 {r_{\mathrm{T}}}^2 - K r_{\mathrm{T}} R_{\mathrm{T}}\right)}{(2C_1 + K)^2}. \tag{25}$$

where $A = \sqrt{(2R_{\mathrm{T}} - r_{\mathrm{T}} - K)^2 + 8KR_{\mathrm{T}}}$, and $B = -K^2 - 2KR_{\mathrm{T}} - 2Kr_{\mathrm{T}} + 2r_{\mathrm{T}}R_{\mathrm{T}} - {r_{\mathrm{T}}}^2$. We next show that only one solution shown in above is positive for all positive rate constants. Suppose

$$C_1 = \frac{B \pm (K + r_{\mathrm{T}})A}{8R_{\mathrm{T}}}. \tag{26}$$

If $B < 0$, then $C_1 < 0$ for all positive rate constants and positive steady states. If $B > 0$, since

$$C_1 = \frac{B - (K + r_{\mathrm{T}})A}{8R_{\mathrm{T}}}. \tag{27}$$

for all positive rate constants and positive steady states, then $C_1 < 0$ under these conditions. Therefore, the only biologically plausible solution to Eq 25 is

$$C_1 = \frac{B + (K + r_{\mathrm{T}})A}{8R_{\mathrm{T}}}. \tag{29}$$

Substituting this into Eq 24 and Eq 23 gives a 2-ODE system that is used for construct phase planes and to illustrate the stability of the steady state in the trace-determinant space of the Jacobian matrix. In particular, the 2D Jacobian matrix

$$J = \begin{pmatrix} \partial f_{\mathrm{T}}/\partial R_{\mathrm{T}} & \partial f_{\mathrm{T}}/\partial r_{\mathrm{T}} \\ \partial g_{\mathrm{T}}/\partial R_{\mathrm{T}} & \partial g_{\mathrm{T}}/\partial r_{\mathrm{T}} \end{pmatrix} \tag{30}$$

was evaluated at various steady state to infer the hidden feedback loops in the MMI2 systems.

*2.1.3 Numerical sampling for the MMI2-SSB Model*

$10^5$ biologically plausible parameter values were randomly drawn from the distributions described in Section 1.5. For each parameter set, numerical continuation was performed with respect to signal $\sigma_R$. The category of the steady state was determined by the Jacobian matrix for Eq 21, or equivalently that for Eq 23. The 2D Jacobian matrix can be further analyzed graphically using a determinant-trace plot. For example, the two representative bifurcation curves shown in Fig. 3B and F are shown in the determinant-trace space in Fig. S4.

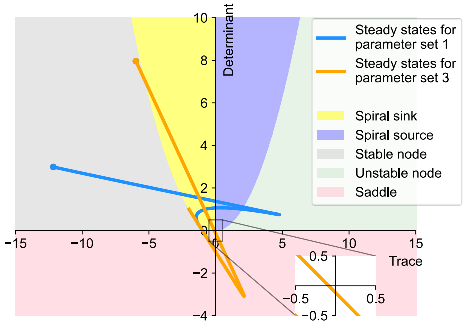

**Fig. S4. Two representative steady state curves projected onto determinant-trace space of the Jacobian matrix.** Two representative parameter sets were chosen. Their steady state signal-response curves with respect to $\sigma_R$ were plotted in Fig. 3B (set 1) and F (set 3) respectively. The curves were projected on to the determinant-trace space of Jacobian matrix of the reduced model (Eq 23). Four colored areas show categories of steady states based. Two curves show the steady states of the system starting from $\sigma_R = 0$ (orange and blue dots) to $\sigma_R = 20$.

We observed that the spiral sink steady states, saddle-node bifurcation points, and Hopf bifurcation points were distributed widely in individual parameters (Fig. S5). However, the ratios of the RDFs had a significant influence on the types of steady state behaviors (Fig. 3E).

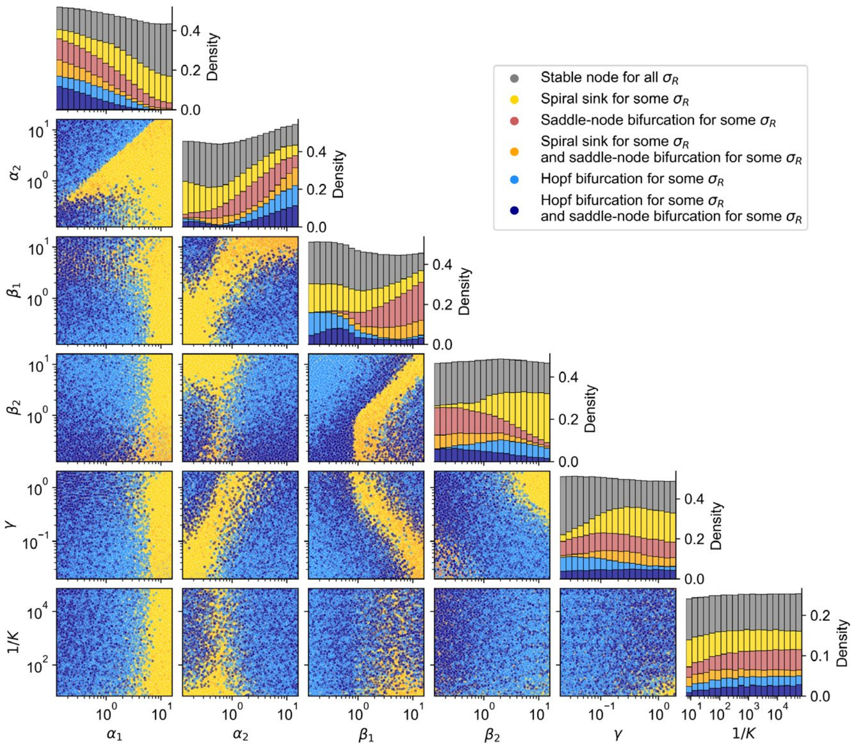

**Fig. S5. Distributions of six types of steady states over individual parameters.** Scatter plots show distributions of $10^5$ random parameters sets categorized by whether they produced spiral sink steady states, saddle-node and Hopf bifurcation points for some values of $\sigma_R$. Spiral sink and Hopf bifurcation points were not considered in a mutually exclusive manner. Marginal distributions are shown in stacked bars.

Interestingly, two regions in the RDF ratio space produced Hopf bifurcations at some values of $\sigma_R$ (Fig. 3E). We found that values of $\gamma$ had a significant influence on the distributions of the parameter sets between the two regions (Fig. S6A). In contrast, $K$ had limited influence on both the appearance of the bifurcation points and the distributions between the two regions for Hopf bifurcation (Figs. S5 and S6). Furthermore, based on the numerical values of the Jacobian matrix near the Hopf bifurcations, we found that the two regions correspond to two implicit feedbacks between mRNA and microRNA respectively (Fig. S6A square heatmaps).

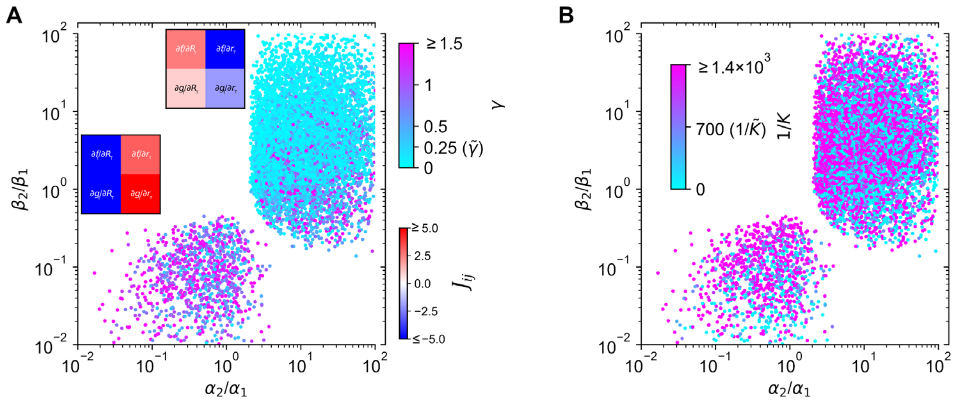

**Fig. S6. Influence of parameters $\gamma$ and $K$ on the distribution of parameter sets that produced Hopf bifurcations**. 9.4% of the $10^5$ random parameters sets produced Hopf bifurcation points for some values of $\sigma_R$ (this plot and Fig. 3E blue). The values of (**A**) $\gamma$, the basal microRNA degradation rate constant, and (**B**) $K$, the dissociation constant, are shown with the indicated color maps. Median values for parameter sampling are labeled in the color map. In panel A, numerical values of Jacobian matrix are shown in square heatmaps for two representative parameter sets near Hopf bifurcations in the two regions respectively. Functions $f_\mathrm{T}$ and $g_\mathrm{T}$ are shown in Eq 23 (2D version of the MMI2-SSB Model).

#### *2.1.4 Sequentially symmetrical binding of a coregulated microRNA*

To test whether coregulation of mRNA and microRNA transcription can affect the parameter region of limit cycle oscillation, we modified the MMI2-SSB Model so that a regulated transcription rate constant $\omega\sigma_R$ is added to the ODE for the microRNA (Fig. S7A). With $\omega = 0.25$, we observed an increase in the distance between the two Hopf bifurcation points (compare Fig. 3B and Fig. S7B), as well as the region allowing for limit cycles (blue areas in Fig. 3B and Fig. S7B). This suggests that the coregulated transcription of mRNA and microRNA (e.g. *mir-196* and *HoxB7*) has a positive effect on the robustness of sustained oscillation. Nonetheless, abrupt changes of the period were still observed with the modified model (Fig. S7B). We further performed two-parameter bifurcation analysis and confirmed the expansion of the region for limit cycles with the increase of $\omega$ (Fig. S7C).

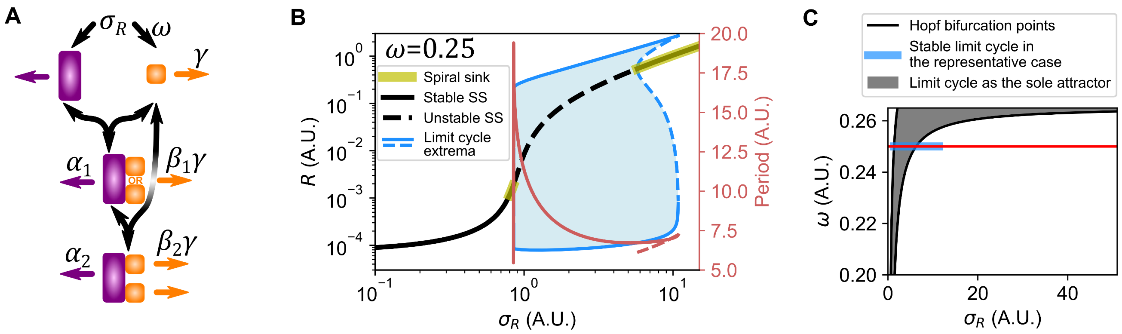

**Fig. S7. Effects of coregulation of microRNA and mRNA transcription**. (**A**) Network of a modified version of the MMI2-SSB Model. The basal synthesis rate constant for $r$ was changed to 0.5. A regulated synthesis constant for $r$, $\omega\sigma_R$, was added to the ODE. (**B**) Bifurcation diagram showing levels of $R$ in response to transcription rate constant $\sigma_R$. Blue shade: limit cycles' inner basins of attraction. Other parameter values: $K = 0.001, \gamma = 0.25, \alpha_1 = \beta_1 = 1, \alpha_2 = 12, \beta_2 = 7, \omega = 0.25$. A basal synthesis rate constant $\sigma_R^0 = 1.5$ was added to the ODE for $R$. The parameter set is similar to that used for Fig. 3B. (**C**) Two-parameter bifurcation diagram with respect to $\sigma_R$ and $\omega$. Red line marks the position of the bifurcation diagram shown in B.

#### *2.1.5 Stochastic simulation of the MMI2-SSB Model*

<u>Additive noise.</u> Noise of biochemical networks can be intrinsic or extrinsic. Intrinsic noise is generated by inherent stochasticity in processes such as binding, transcription, and degradation. Two simulation approaches are widely used to simulate intrinsically stochastic biochemical processes: the chemical Langevin equation (CLE) or the Gillespie algorithm, both of which accurately describe intrinsic noise of a system. However, it should be noted that the two algorithms only consider noise generated by the most elementary steps of the biochemical reactions. In practice, the assumptions for the intrinsic nature are often not realistic. For example, the two algorithms assume that the production of molecules (i.e. transcription, splicing, nuclear transport etc.) is a Poisson process (Lan et al., 2008), which does not reflect the pulsatory dynamics of mRNA production controlled by sophisticated machinery beyond the modeled molecular

species (Chubb et al., 2006). Therefore, extrinsic noise can be a reasonable description of the stochasticity in the RNA networks. Here, we first use a general form of stochastic ODE based on Eq 21 to simulate the MMI2-SSB Model:

$$\begin{aligned} dR &= f(R, r, C_1, C_2)dt + \omega_R d\xi_R \\ dr &= g(R, r, C_1, C_2)dt + \omega_r d\xi_r \\ dC_1 &= h(R, r, C_1, C_2)dt \\ dC_2 &= k(R, r, C_1, C_2)dt \end{aligned} \tag{31}$$

where $d\xi_i$ describes standard Brownian motion, $\omega$ represents the amplitude of the noise, and $f$, $g$, $h$, and $k$ are the deterministic rates of change. We chose representative noise levels with $\omega_R = 1.4$ and $\omega_r = 0.35$ such that the MMI2-SSB Model produced bimodal distributions of total mRNA with two parameter sets: $\sigma_R = 0.3$, $K = 0.001$, $\gamma = 0.25$, $\alpha_1 = \beta_1 = 1$, $\alpha_2 = 7$, $\beta_2 = 4$, and a basal synthesis rate constant $\sigma_R^0 = 5.7$ for $R$ added to $f$ (a condition for diverging oscillator); $\sigma_R = 3.0$, $K = 0.001$, $\gamma = 2$, $\alpha_1 = 1$, $\beta_1 = 0.5$, $\alpha_2 = 4$, $\beta_2 = 0.1$ (a condition for bistable switch, Fig. 3F). To test the sensitivity of gene expression patterns to noise level, we changed the amplitudes through multiplication of $\omega_R$ and $\omega_r$ by some constants (high noise: 1.5; low noise: 0.5) reflecting the dynamical nature of the stochasticity, e.g. the change of the cell volume or the global control of the transcriptional activities (Fig. 6C-D).

To simulate a cell population with homogeneous initial conditions, a scenario similar to a sorted cell population, we first solved the ODEs with 400 initial conditions obtained by scanning a 20×20 grid in the $R$ and $r$ space with each variable ranging in $[0.001, 10.0]$. Other variables were assumed to be 0 at $t = 0$. Because the sorted $R$-high and a $R$-low cells do not need to be at a stable point attractor, we took the state where the level of $R$ is the overall minimum value in the time window $[40, 200]$ among all initial conditions as the $R$-low state. Similarly, the state where the level of $R$ is the overall maximum value in the time window was assumed to be the $R$-high state. We next used one of these states as the initial condition for 500 cells and performed simulations for the stochastic ODEs described above (Rackauckas and Nie, 2017). Trajectories of single cells and the time evolution of the expression profiles of the population were subsequently analyzed (see the upcoming Section 2.1.7 for details).

Multiplicative noise. Because additive noise and multiplicative noise may influence the dynamics differently (Coomer et al., 2021), we considered an alternative model with multiplicative noise,

$$\begin{aligned} dR &= f(R, r, C_1, C_2)dt + R\omega_R d\xi_R \\ dr &= g(R, r, C_1, C_2)dt + r\omega_r d\xi_r \\ dC_1 &= h(R, r, C_1, C_2)dt + C_1\omega_{C_1} d\xi_{C_1} \\ dC_2 &= k(R, r, C_1, C_2)dt + C_2\omega_{C_2} d\xi_{C_2}. \end{aligned} \tag{32}$$

Two variables, equivalent deterministically but subject to independent stochastic processes, represented the two forms of the complex $C_1$. Because multiplicative noise reflects more intrinsic properties of the system, we considered the noise for all variables. We took the simple assumption that the basal noise $\omega_R = \omega_r = \omega_{C_1} = \omega_{C_2} = 1.4$. We performed the simulations with this model under the same conditions used for the model with additive noise.

Gillespie's algorithm. We considered a third stochastic model using the full Gillespie algorithm (Choi et al., 2018; Gillespie, 1977) with propensities specified by scaling (Lecca, 2013) the MMI2-SSB Model, i.e.

$$\xrightarrow{\sigma_R V} \hat{R} \qquad \hat{R} \xrightarrow{1} \emptyset \qquad \xrightarrow{\sigma_r V} \hat{r} \qquad \hat{r} \xrightarrow{\gamma} \emptyset \qquad \hat{R} + \hat{r} \xrightarrow{\kappa^{\mathrm{on}}/V} \hat{C}_1 \qquad \hat{C}_1 \xrightarrow{\kappa^{\mathrm{off}}} \hat{R} + \hat{r} \qquad \hat{C}_1 \xrightarrow{\gamma\beta_1} \hat{R}$$

$$\hat{C}_1 \xrightarrow{\alpha_1} \hat{r} \qquad \hat{C}_1 + \hat{r} \xrightarrow{\kappa^{\mathrm{on}}/V} \hat{C}_2 \qquad \hat{C}_2 \xrightarrow{\kappa^{\mathrm{off}}} \hat{C}_1 + \hat{r} \qquad \hat{C}_2 \xrightarrow{\gamma\beta_2} \hat{C}_1 \qquad \hat{C}_2 \xrightarrow{\alpha_2} 2r \qquad (33)$$

Each reaction producing or consuming $\hat{C}_1$ was simulated as two separate reactions with the same propensity function but different $C_1$ species to reflect the two forms of the 1:1 complex. Change of intrinsic noise in this case can be viewed as the change of cell volume, reflected in the adjustment of the cell volume $V$ in our simulations. To convert concentrations to the number of molecules or vice versa, we multiplied or divided each variable by the volume $V$. Because simulating the system with the full Gillespie algorithm is computationally expensive, we only considered some representative conditions and simulated 200 cells for each condition/parameter set. The conclusions about the compared models' ability to restore heterogeneity were consistent with those obtained with the stochastic ODEs (Fig. S12).

#### *2.1.6 Quasi-potential landscapes*

We constructed quasi-potential landscapes of the MMI2-SSB Model with additive and multiplicative noise (Eqs 31 and 32 respectively). We first chose a parameter set that generates SNIC bifurcation (Fig. 5B) as a representative *diverging oscillator* model. We simulated the stochastic ODEs for 500 cells and 200 time units. We assumed that the trajectories of the cells are near the stationary phase after 100 time units and computed the quasi-potential as $U(x) = -\log P_S(x)$, where $P_S(x)$ is the probability density function at the stationary phase (Fig. 6A). To visualize the trajectories of the cells on the landscape, we randomly selected 50 cells and plotted the position of the cells at time 50 in the state-potential space with their trajectories in the previous 0.4 time units (Fig. 6A dark blue). We performed the same analysis for a model with only saddle-node bifurcations (Fig. 3F). The landscapes with additive noise are shown in Fig. 6A-B and Fig. S8, and those with multiplicative noise are shown in Fig. S9C-D. Landscapes under medium and high additive noise conditions, as well as representative trajectories of stochastic simulations, are also shown in Movies S1 and S2.

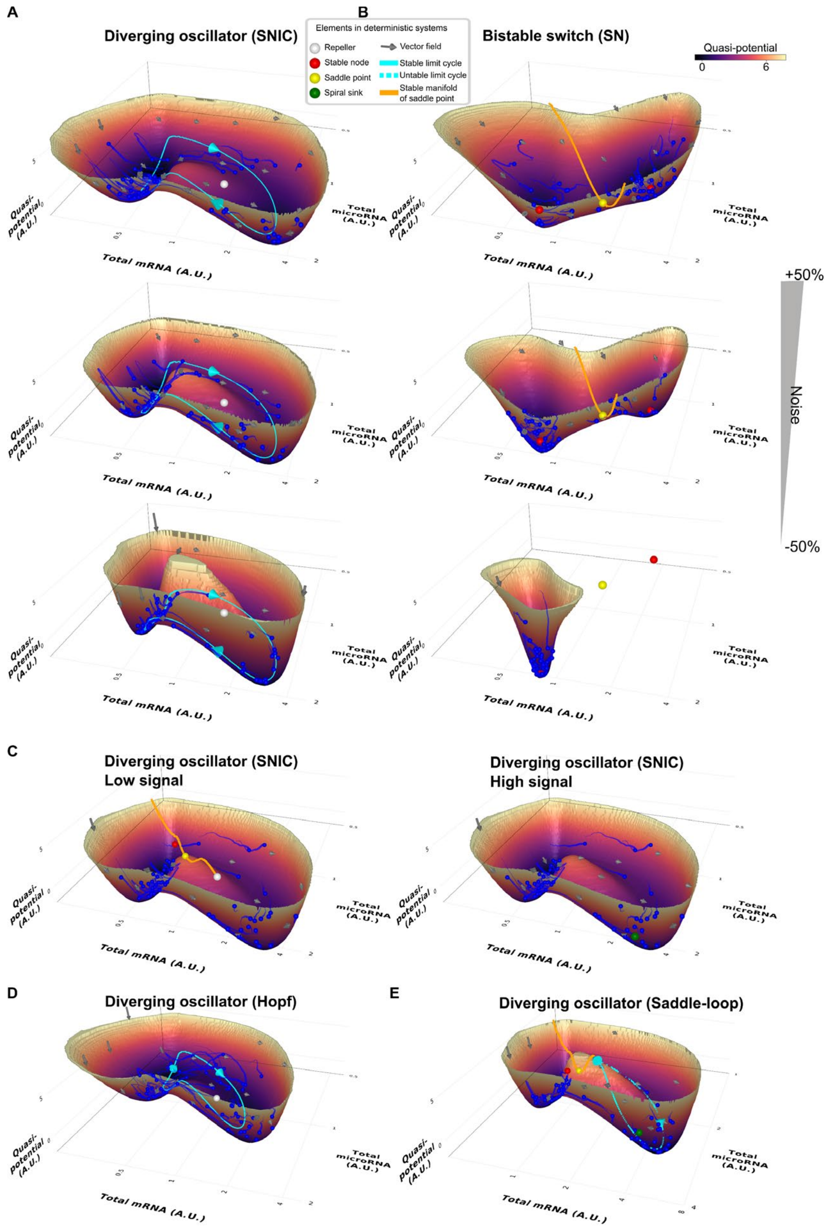

**Fig. S8. Quasi-potential landscapes with additive noise**. (**A**) Landscapes (shown as surface) of the SNIC-generating MMI2-SSB Model with additive noise describing stochastic transcription of mRNA and microRNA (basal noise level: $\omega_R = 1.4, \omega_r = 0.35$). Landscapes were computed based on stationary phase distribution of stochastic simulations of 500 cells. For each landscape, 50 cells were randomly selected and visualized with the positions at Time 50 (blue spheres) and trajectories in a 0.4 time-unit period (blue tails). Top, middle, and bottom panels show landscapes under high (+50%), basal, and low (-50%) noise levels respectively. Parameter values: $K = 0.001, \gamma = 0.25, \alpha_1 = \beta_1 = 1, \alpha_2 = 12, \beta_2 = 4, \sigma_R = 0.3, \sigma_R^0 = 5.7$ (same as in Fig. 5B). (**B**) Landscape, as A, for the model generating SN bifurcation: $K = 0.001, \gamma = 2,$ $\alpha_1 = 1, \beta_1 = 0.5, \alpha_2 = 4, \beta_2 = 0.1, \sigma_R = 3.0, \sigma_R^0 = 0$ (same as in Fig. 3F). (**C**) Landscape for the SNIC-generating model with $\sigma_R = 0.24$ and $\sigma_R = 0.7$. Other parameter values are the same as in A. (**D**) Landscape for the model that generates Hopf bifurcation but not homoclinic bifurcation (Fig. 3B). $\sigma_R =$ 0.6. Other parameter values are the same as in Fig. 3B. (**E**) Landscape for the model that generates saddle-loop bifurcation but not SNIC bifurcation. $\sigma_R = 0.92$. Other parameter values are the same as in Fig. 5G.

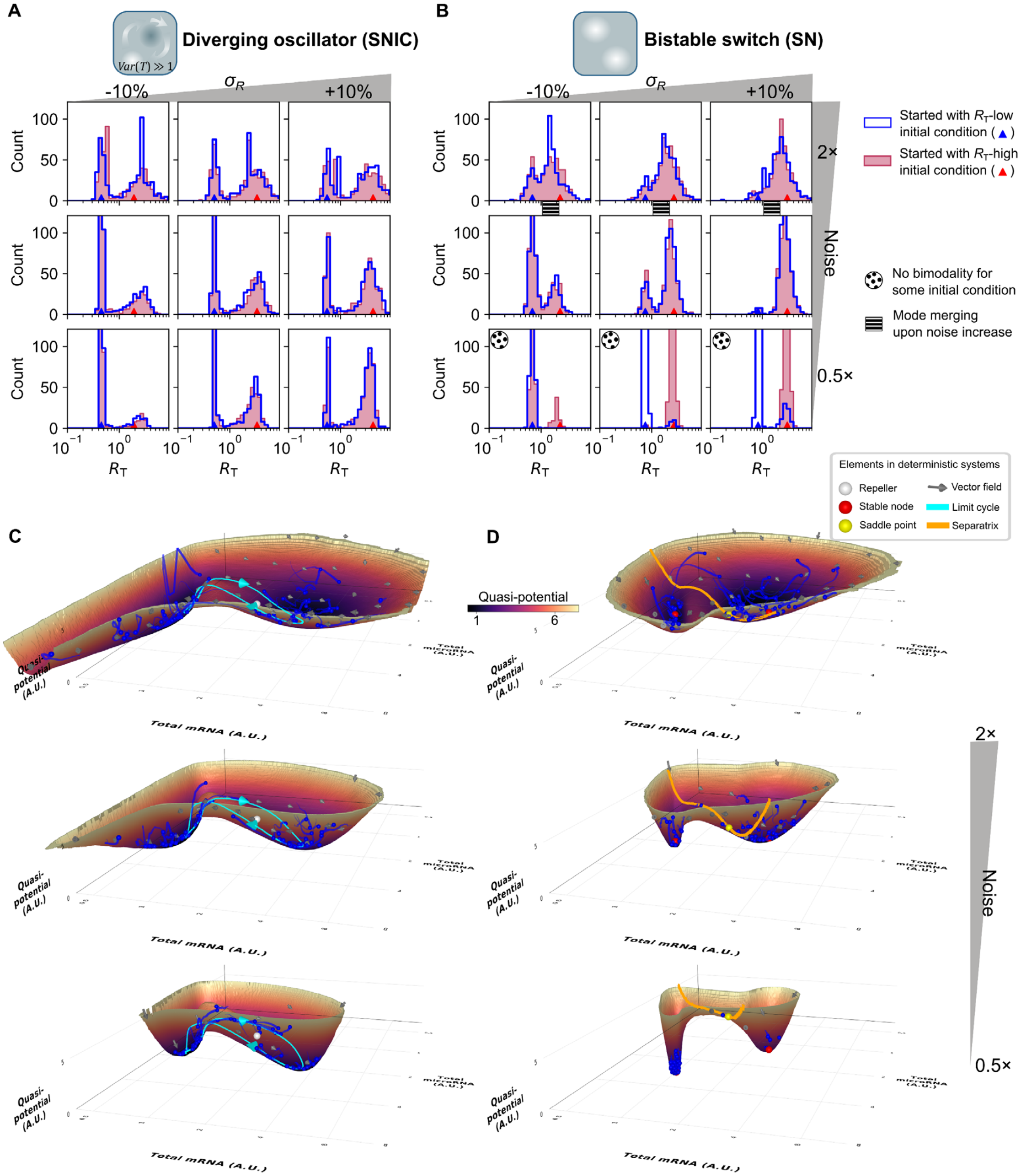

**Fig. S9. Bimodality regeneration and quasi-potential landscapes with multiplicative noise.** (**A**-**B**) Distribution of the total mRNA in the MMI2-SSB Model at time 100 with multiplicative noise describing stochasticity of all molecular species (basal noise level: $\omega_R = \omega_r = \omega_{C_1} = \omega_{C_2} = 1.4$). Time 100 corresponds to 576 hours after cell sorting, assuming a 4-hour mRNA half-life. Parameter values for the model generating SNIC bifurcation (A): $K = 0.001, \gamma = 0.25, \alpha_1 = \beta_1 = 1, \alpha_2 = 12, \beta_2 = 4$, basal signal level (middle column): $\sigma_R = 0.3, \sigma_R^0 = 5.7$. Parameter values for the model generating SN bifurcation (B): $K = 0.001$, $\gamma = 2$, $\alpha_1 = 1$, $\beta_1 = 0.5$, $\alpha_2 = 4$, $\beta_2 = 0.1$. Basal signal level: $\sigma_R = 3.0$, $\sigma_R^0 = 0$ (middle column). Top, middle, and bottom panels show distributions under high (2× basal), basal, and low (0.5× basal) noise levels respectively. (**C**-**D**) Landscapes (shown as surface) of the MMI2-SSB Model under the same noise conditions with the basal signal levels.

*2.1.7 Robustness and speed of heterogeneity restoration*

To compare the heterogeneity-restoring performance of different dynamical/bifurcation behaviors (parameter sets in Table S1), we simulated populations of cells under several noise levels—those used in Fig. 6C-D—and a range of $\sigma_R$ signal levels in increments of 0.1 in or near each parameter set's bifurcating regime.

| **Parameter** | **SNIC** | **Saddle-loop** | **Diverging Hopf** | **Saddle-node** |
|---|---|---|---|---|
| $\sigma_R^0$ | 5.7 | 6.9 | 3.1 | 0 |
| $\sigma_r$ | 1 | 1 | 1 | 1 |
| $\alpha_1$ | 1 | 1 | 1 | 1 |
| $\alpha_2$ | 12 | 12 | 12 | 4 |
| $\beta_1$ | 1 | 1 | 1 | 0.5 |
| $\beta_2$ | 4 | 3 | 7 | 0.1 |
| $\gamma$ | 0.25 | 0.25 | 0.25 | 2 |
| $\sigma_R$ (minimum) | -1.0 (add. noise)<br>-2.0 (mul. noise) | -1.0 (add. noise)<br>-2.0 (mul. noise) | -1.0 | 1.5 |
| $\sigma_R$ (maximum) | 2.0 (add. noise)<br>3.0 (mul. noise) | 3.0 | 2.0 | 3.5 |

**Table S1.** Parameter sets giving rise to each bifurcation behavior with $\sigma_R$ as the control variable; ranges of $\sigma_R$ tested for heterogeneity restoration. In all cases, $\kappa^{\text{off}} = 10^2$ and $\kappa^{\text{on}} = 10^5$. Add., additive; mul., multiplicative.

After simulation for 200 time units, the distribution of the 500 cells' final total mRNA concentration was tested for bimodality using two metrics. First, the Bayesian information criterion (BIC) of a single-Gaussian fit was compared to that of a two-Gaussian mixture fit. The GaussianMixtures.jl library was used to fit the models and compute the average log-likelihood $\mu_{\text{LL}}$. Following the conventions of scikit-learn (Pedregosa et al., 2011), each BIC was computed as

$$\text{BIC}_m = (3m - 1)\ln n_{\text{obs}} - 2\mu_{\text{LL}} n_{\text{obs}} \tag{34}$$

where $m$ is the number of Gaussians, required to compute the number of free parameters, and $n_{\text{obs}}$ is the number of observations, i.e. cells. To be considered bimodal, a simulation endpoint was required to satisfy

$$\Delta\text{BIC} = \text{BIC}_1 - \text{BIC}_2 > 0. \tag{35}$$

Because the data can be nonnormally distributed without possessing the bimodality expected from the restoration of two subpopulations, two-Gaussian mixture models were further tested for separation of the modes (Muratov and Gnedin, 2010), requiring

$$D = \frac{|\mu_1 - \mu_2|}{\sqrt{\frac{\sigma_1^2 + \sigma_2^2}{2}}} > 2 \tag{36}$$

where $\mu_n$ is a mean of a Gaussian and $\sigma_n^2$ is its variance. By testing these constraints, we detected distinguishable subpopulations with little sensitivity to their relative weights. To reduce the effect of nondeterminism in model fitting, the Gaussian models were fit four times for each dataset, producing four values for each of the two bimodality metrics, of which the minimum was taken. The robustness of a mechanism in regenerating heterogeneity at a given noise level was quantified as the largest range of $\sigma_R$ in which it consistently produced a bimodal final population (Figs. S10 and S11).

To compare the speeds at which the different mechanisms restore heterogeneity, each timecourse was analyzed to estimate the point at which the population reached equilibrium, settling into its consistent long-term behavior. Because the experimentally measurable state of the population may depend on total mRNA while the change in the system depends on individual species, it is difficult to distinguish stochastic fluctuations from an incomplete population-level transient based on the timecourse of only one state value. Therefore, metrics involving both total and free mRNA were examined. The similarity of each timepoint's total mRNA distribution was compared to that at the final timepoint by the root mean square deviation (RMSD) between the histogram bin counts with boundaries ranging from 0 to 10 in steps of 0.25. The maximum value of this RMSD reached "late," after time 100, was taken as the deviation permitted by stochastic fluctuations at equilibrium. To be considered to have reached equilibrium, the beginning of the simulation was required to attain an RMSD less than half of the late maximum and remain below 110% of the late maximum for the next 5 time units after the putative equilibrium point. Additionally, the mean free mRNA concentration was computed at each timepoint. The minimum, mean, and maximum values of this population-level mean from time 100 to the end of the simulation were similarly recorded. Reaching equilibrium required that the free mRNA mean cross the late mean mean and remain between the late minimum mean and late maximum mean for the next 5 time units. The equilibrium time was considered the first timepoint within the first 100 time units at which both the total mRNA RMSD and free mRNA mean conditions were satisfied.

| Parameter set | Noise | Minimum $\sigma_R$ | Maximum $\sigma_R$ | Median ET | | Maximum ET |
|---|---|---|---|---|---|---|
| | | | | Lo IC | Hi IC | |
| SNIC | -50% | -0.1 | 1.0 | 16.5 | 21.3 | 59.6 |
| | Basal | -0.3 | 1.4 | 10.7 | 16.3 | 32.7 |
| | +50% | -0.6 | 1.6 | 11.2 | 8.9 | 23.6 |
| Saddle-loop | -50% | 0.4 | 1.3 | 18.1 | 25.3 | 69.8 |
| | Basal | 0.1 | 1.8 | 17.4 | 17.2 | 37.7 |
| | +50% | -0.2 | 2.1 | 11.3 | 12.1 | 25.0 |
| Diverging Hopf | -50% | 0.2 | 0.9 | 11.3 | 9.3 | 23.6 |
| | Basal | 0.2 | 1.3 | 5.5 | 5.3 | 23.2 |
| | +50% | 0.0 | 1.5 | 6.1 | 4.4 | 11.2 |
| Saddle-node | -50% | NA | NA | NA | NA | NA |
| | Basal | 2.8? | 3.1? | > 100 | > 100 | > 100 |
| | +50% | 2.5 | 3.2 | 27.4 | 29.1 | 71.5 |

**Table S2A.** Heterogeneity restoration performance for MMI2-SSB parameter sets under additive noise. Summary statistics for equilibrium times were computed only for signal values that restored heterogeneity from both high and low initial conditions. ET, equilibrium time; IC, initial condition; NA, not applicable due to nonexistent or undetected range of consistent heterogeneity restoration; question mark, lasting heterogeneity restoration unclear due to very slow and potentially incomplete population-level transient.

| Parameter set | Noise | Minimum $\sigma_R$ | Maximum $\sigma_R$ | Median ET | | Maximum ET |
|---|---|---|---|---|---|---|
| | | | | Lo IC | Hi IC | |
| SNIC | -50% | -0.3 | 2.0 | 9.6 | 10.3 | 34.8 |
| | Basal | -1.7 | 2.3 | 8.4 | 8.1 | 23.8 |
| | +50% | -0.2 | 0.1 | 8.7 | 7.0 | 27.5 |
| Saddle-loop | -50% | -0.9 | ≥ 3.0 | 12.3 | 11.9 | 47.9 |
| | Basal | -0.7 | 2.7 | 9.8 | 10.0 | 35.4 |
| | +50% | -0.2 | 1.2 | 8.1 | 6.9 | 16.9 |
| Diverging Hopf | -50% | 0.1 | 1.1 | 8.5 | 9.6 | 16.0 |
| | Basal | -0.3 | 1.3 | 5.7 | 5.8 | 20.9 |
| | +50% | 0.1 | 0.4 | 8.9 | 4.8 | 13.1 |
| Saddle-node | -50% | 2.8? | 3.0? | > 100 | > 100 | > 100 |
| | Basal | 2.6 | 3.0 | 28.7 | 28.4 | 47.1 |
| | +50% | 2.5 | 2.6 | 20.3 | 15.1 | 30.2 |

**Table S2B.** As A, for multiplicative noise.

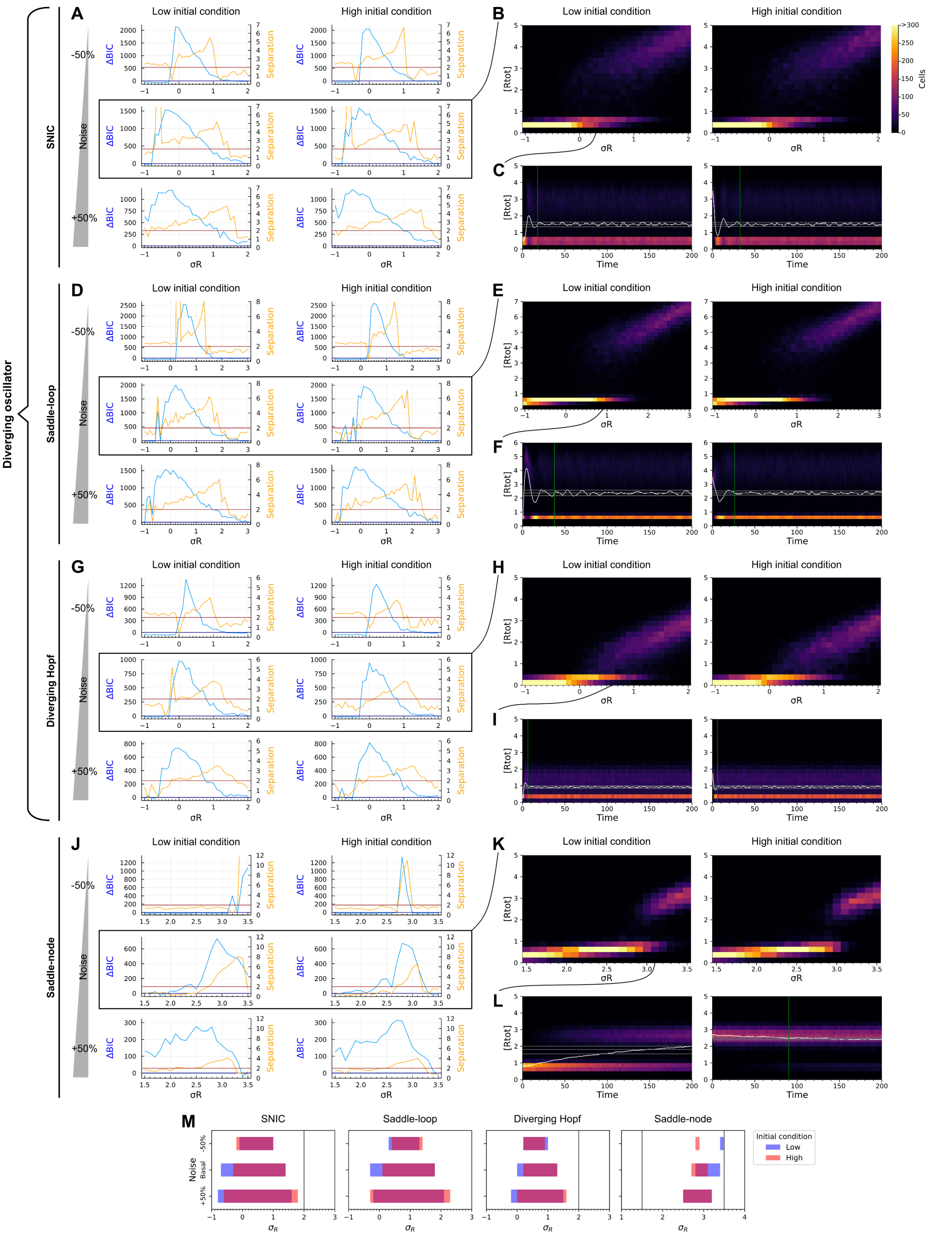

**Fig. S10. Bimodality restoration performance scan of the MMI2-SSB Models under additive noise.** (**A**) Representative values of bimodality metrics for the SNIC model under a range of signals for different initial conditions and noise levels. When the blue line is above the dark blue line, ΔBIC > 0. When the orange line is above the dark orange line, $D > 2$. (**B**) Distribution of total mRNA concentration after 200 time units of the SNIC model under basal noise. (**C**) Timecourses of total mRNA distribution for $\sigma_R = 0.3$ in the SNIC model under basal noise. White line, population mean; gray lines, "late" maximum, mean, and minimum mean after time 100; green line, equilibrium time if detectable. Color scale as in B. (**D-E**) As A-B for the saddle-loop model. (**F**) As C for $\sigma_R = 0.9$ in the saddle-loop model. (**G-H**) As A-B for the diverging Hopf model. (**I**) As C for $\sigma_R = 0.6$ in the diverging Hopf model. (**J-K**) As A-B for the saddle-node model. (**L**) As C for $\sigma_R = 3.1$ in the saddle-node model. (**M**) Ranges of $\sigma_R$ in which each model restored bimodality in each noise level for each initial condition's simulation. Purple overlap, reliable restoration; gray lines, extrema of $\sigma_R$ scan when different from axis limits.

The diverging oscillators were able to restore heterogeneity within the tested time range under a much broader range of signal than the switch-like saddle-node system at all noise levels (Table S2A, Fig. S10M). Increasing the level of additive noise increased all models' ability to restore heterogeneity. The diverging oscillators exhibited transient population-level oscillations as seen experimentally (Fig. S10C/F/I), while cell state changes in the saddle-node system occurred near-linearly and very slowly, sometimes failing to reach equilibrium within 100 time units (Fig. S10L). The SNIC model in particular exhibits a delay of at least one time unit before any appreciable appearance of cells in the opposite state as selected (Fig. S10C). The diverging oscillators' higher robustness (Table S2B, Fig. S11M) and nonlinear transients (Fig. S11C/F/I) were also observed under multiplicative noise, though at high levels of multiplicative noise, all models produced less detectable bimodality because the populations began to overlap each other (low separation in Fig. S11A/D/G/J).

When simulated with Gillespie's exact algorithm, the diverging oscillators were again able to restore bimodal gene expression under wider signal ranges and lower noise levels than the bistable switch (Fig. S12A light green). Indeed, the bistable switch was only able to produce bimodal expression at very high noise, simulated as low cell volume. The diverging oscillators typically exhibited the experimentally observed damped oscillations (Fig. S12B), with the SNIC model showing a distinct delay of several time units before the appearance of cells in the opposite state as the initial condition.

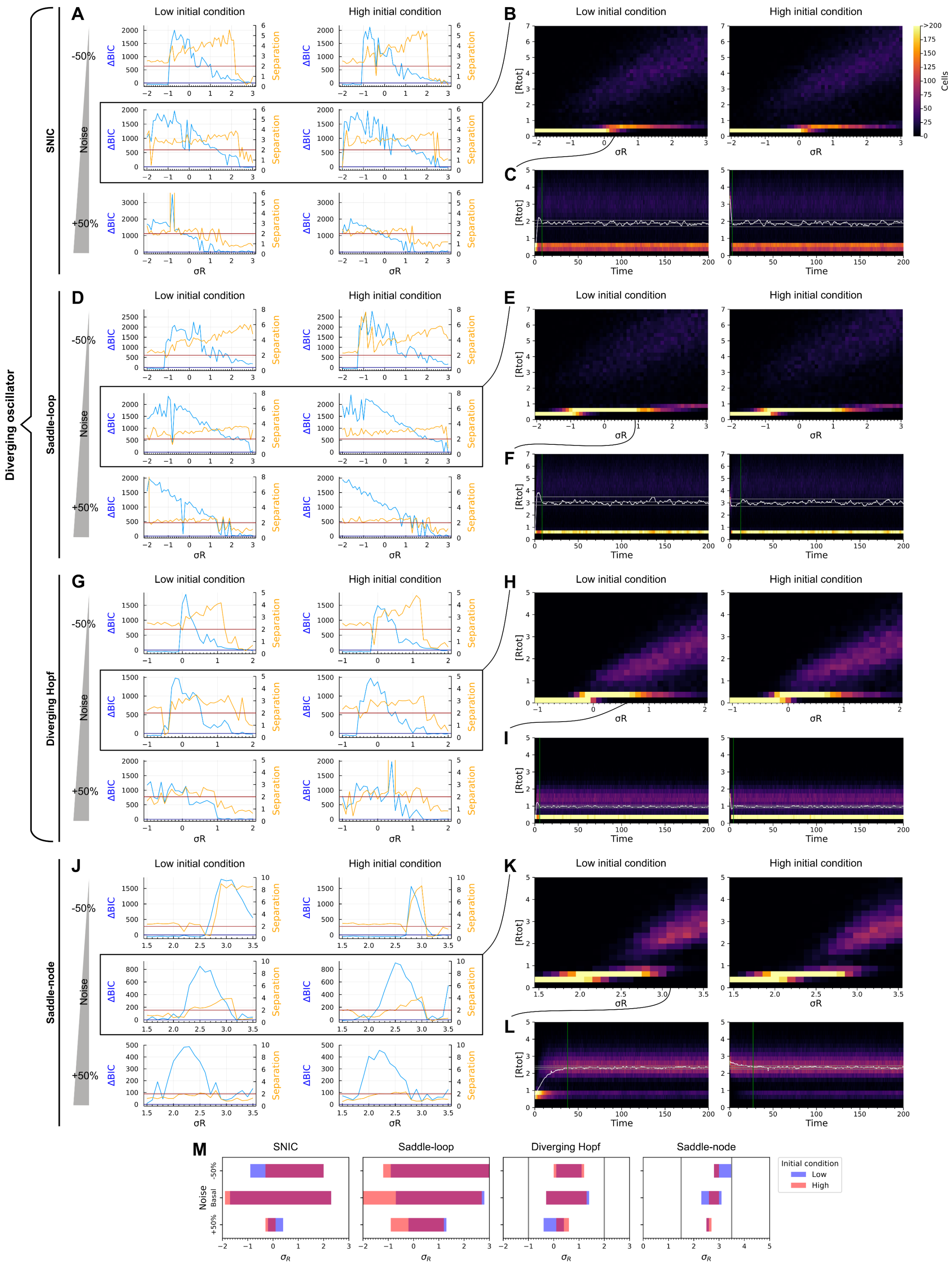

**Fig. S11. Bimodality restoration performance scan of the MMI2-SSB Models under multiplicative noise.** Analogous to Fig. S10.

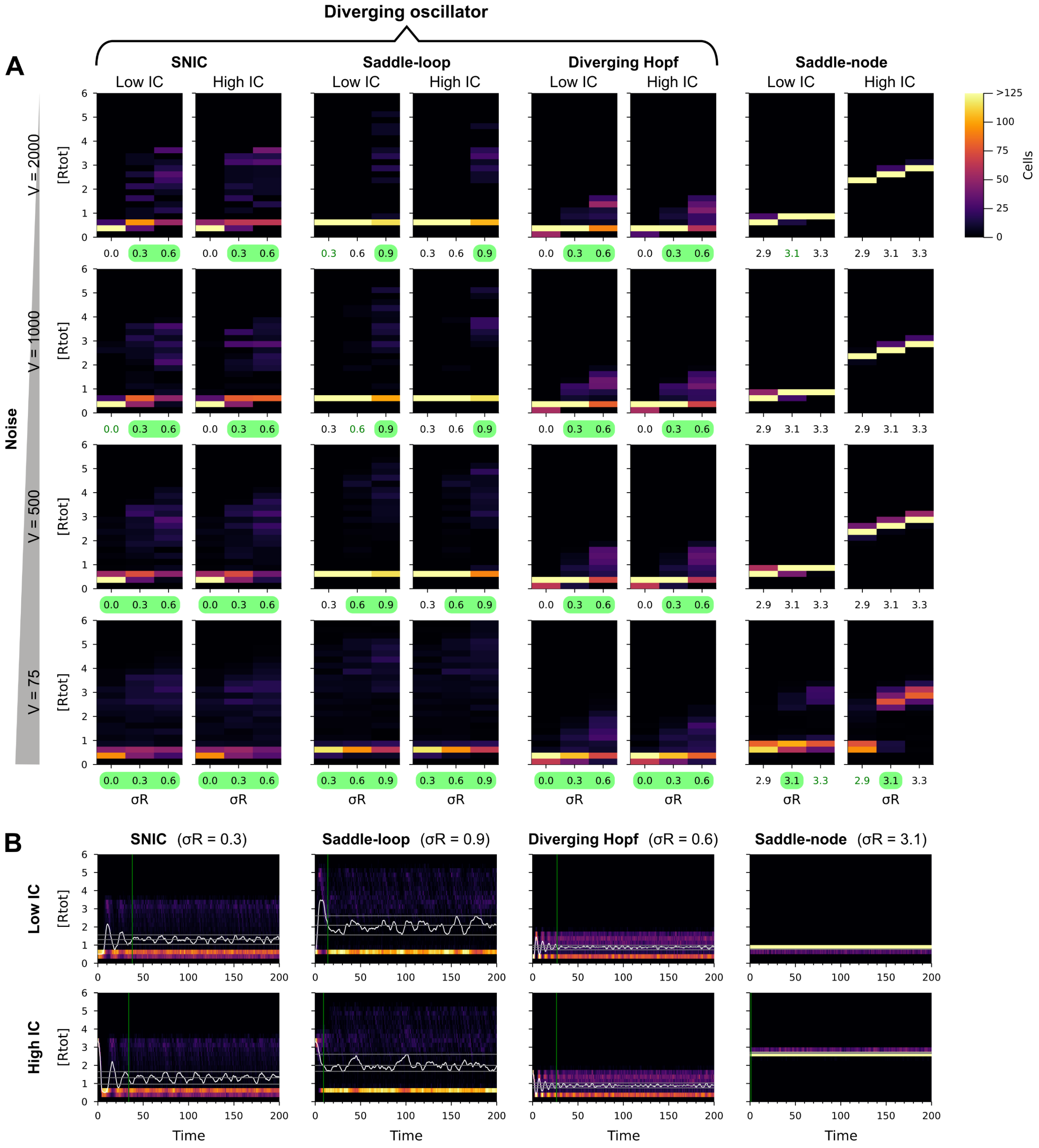

**Fig. S12. Behavior of the MMI2-SSB Model under exact Gillespie stochasticity.** (**A**) Final distributions of total mRNA concentration starting from two initial conditions under several noise levels. Noise—the impact of stochasticity—increases as the simulated cell volume $V$ decreases. IC, initial condition; light green highlight, consistently detectable bimodality starting from either initial condition; dark green text, inconsistently detectable bimodality. (**B**) Timecourses at $V = 1000$ of representative parameter sets with deterministically distinguishable low and high states. White line, population mean; gray lines, "late" maximum, mean, and minimum mean after time 100; green line, equilibrium time. Color scale as in A.

*2.1.8 Stochastic simulation with cell division*

To further test that our results are relevant to a growing cell population, we considered an extended version of the model in Eq 31. In addition to a stochastic differential equation for each species, each cell's system contained a stochastic differential equation for a variable representing its progress through the cell cycle:

$$dG = \frac{G_{\max}}{t_{\text{div}}}dt + \omega_G d\xi_G \tag{37}$$

where $G_{\max}$ is the value of $G$ at which the cell divides, $t_{\text{div}}$ is the average number of time units per division, and $\omega_G$ is the amplitude of the additive noise.

The simulation is initialized by finding an extreme initial condition as in Section 2.1.5. For each initial cell, one instance of the system is created with the initial condition's concentrations and a $G$ value uniformly randomly selected from $[0, G_{\max}]$. When a cell's $G$ value reaches $G_{\max}$, its $G$ is reset to zero and a new daughter cell is created. For each nonzero species concentration $x$, the proportion $p$ of the species' molecules inherited by the new daughter cell is selected from a continuous approximated binomial distribution: a truncated normal distribution on $[0, 1]$ with mean $1/2$ and standard deviation $1/(2\sqrt{2xV})$ where $V$ represents the number of molecules per unit concentration. The new daughter cell's concentration value is then $2px$ and the existing cell's is $2(1-p)x$.

The results of a representative simulation are shown in Fig. S13. As with the fixed-size population simulations, the diverging oscillator models can produce an $R_T$-high subpopulation following population-level oscillation even at low noise.

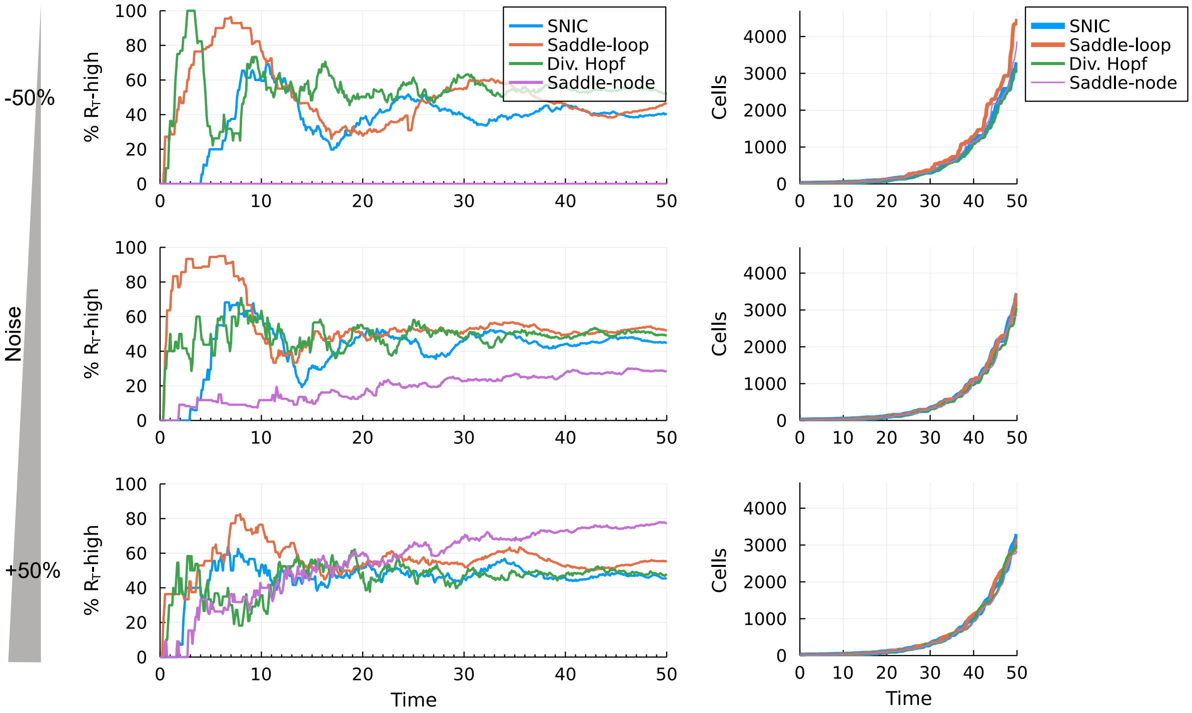

**Fig. S13. Behavior of a growing population under additive noise.** Left: timecourses of the percent of cells in the $R_T$-high state. Right: timecourses of the number of cells in the population. $G_{\max} = 75$, $t_{\text{div}} = 6$, $V = 2000$, and $\omega_G = \omega_R$. The population initially consisted of 10 cells in the low-expression state. Noise as in Section 2.1.5, other parameters as in Table S1, and all cutoffs as in Fig. 6E.

#### *2.2 The MMI2 Model with sequentially asymmetrical binding (MMI2-ASB)*

The chemical reactions of the MMI2-ASB Model are identical to those for the MMI2-SSB Model. There are two differences between the two models. First, $C_1$ (equivalently $\hat{C}_1$) in the MMI2-ASB Model represents a unique form of 1:1 complex instead of one of two identical species. Secondly, the association rate constants of the two sequential binding events are assumed to be independent, represented by $k_1^{\text{on}}$ and $k_2^{\text{on}}$, respectively. The same asymmetrical assumption is made for the dissociation constants $k_1^{\text{off}}$ and $k_2^{\text{off}}$. The nondimensionalized ODEs for this model are

$$
\begin{aligned}
dR/dt &= \sigma_R - \kappa_1^{\text{on}} Rr + \kappa_1^{\text{off}} C_1 - R + \beta_1 \gamma C_1 \\
dr/dt &= 1 - \kappa_1^{\text{on}} Rr + \kappa_1^{\text{off}} C_1 - \kappa_2^{\text{on}} C_1 r + \kappa_2^{\text{off}} C_2 - \gamma r + \alpha_1 C_1 + 2\alpha_2 C_2 \\
dC_1/dt &= \kappa_1^{\text{on}} Rr - \kappa_1^{\text{off}} C_1 - \kappa_2^{\text{on}} C_1 r + \kappa_2^{\text{off}} C_2 - \alpha_1 C_1 - \beta_1 \gamma C_1 + \beta_2 \gamma C_2 \\
dC_2/dt &= \kappa_2^{\text{on}} C_1 r - \kappa_2^{\text{off}} C_2 - \alpha_2 C_2 - \beta_2 \gamma C_2.
\end{aligned}
\tag{38}
$$

These equations are used to perform numerical simulations and bifurcation analysis for the MMI2-ASB Model. We found that it is not feasible to use the tQSSA to reduce this model to a 2-ODE system.

Applying the CRNT to the network showed that the MMI2-ASB Model can have two stable steady states. Consistent with this result, numerical bifurcation analysis with random parameter values using the same scheme described earlier produced parameter sets that allow the MMI2-ASB to be bistable. Representative bifurcation diagrams for Hopf and saddle-node bifurcation points are shown in Fig. S14.

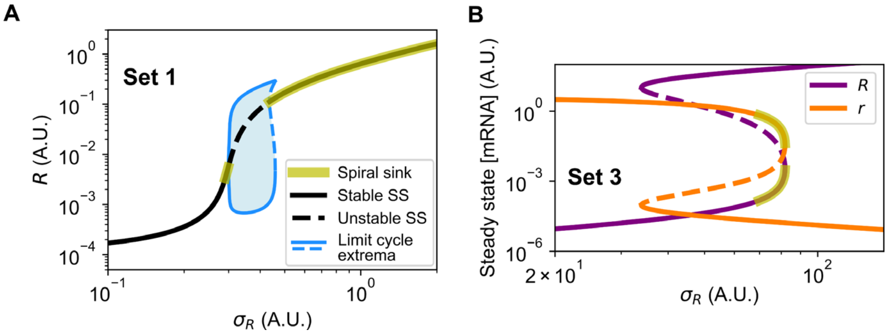

**Fig. S14. Hopf and saddle-node bifurcations with the MMI2-ASB Model**. Parameter values are identical to those used to generate Fig. 3B and F. (**A**) Bifurcation diagram showing levels of $R$ in response to transcription rate constant $\sigma_R$. Blue shade: limit cycles' inner basins of attraction. Solid curves: stable steady states. Dashed curves: unstable steady states. Parameter values: $K = 0.001$, $\gamma = 0.25$, $\alpha_1 = \beta_1 = 1$, $\alpha_2 = 12$, $\beta_2 = 7$. (**B**) Bifurcation diagram showing the steady states of unbound mRNA and unbound microRNA with respect to $\sigma_R$. Solid curves: stable steady states. Dashed curves: unstable steady states. The stable steady state contains a segment of spiral sink (yellow) but returns to a node before undergoing a saddle-node bifurcation. Parameter values: $K = 0.001$, $\gamma = 2$, $\alpha_1 = 1$, $\beta_1 = 0.5$, $\alpha_2 = 4$, $\beta_2 = 0.1$.

#### *2.3 The MMI2 Model with dual microRNAs (MMI2-DMI)*

The chemical reactions of the MMI2-DMI Model are

$$
\begin{aligned}
&\xrightarrow{s_R}\hat{R} \qquad \hat{R}\xrightarrow{k_R^0}\emptyset \qquad \xrightarrow{s_r}\hat{r} \qquad \hat{r}\xrightarrow{k_r^0}\emptyset \qquad \xrightarrow{s_r'}\hat{r}' \qquad \hat{r}'\xrightarrow{k_r^{0\prime}}\emptyset \\
&\hat{R}+\hat{r}\xrightarrow{\kappa^{\mathrm{on}}}\hat{C}_1 \qquad \hat{C}_1\xrightarrow{\kappa^{\mathrm{off}}}\hat{R}+\hat{r} \qquad \hat{C}_1\xrightarrow{k_r^1}\hat{R} \qquad \hat{C}_1\xrightarrow{k_R^1}\hat{r} \\
&\hat{R}+\hat{r}'\xrightarrow{\kappa^{\mathrm{on}\prime}}\hat{C}_1' \qquad \hat{C}_1'\xrightarrow{\kappa^{\mathrm{off}\prime}}\hat{R}+\hat{r}' \qquad \hat{C}_1'\xrightarrow{k_r^{1\prime}}\hat{R} \qquad \hat{C}_1'\xrightarrow{k_R^{1\prime}}\hat{r}' \qquad (39) \\
&\hat{C}_1+\hat{r}'\xrightarrow{\kappa^{\mathrm{on}}}\hat{C}_2 \qquad \hat{C}_2\xrightarrow{\kappa^{\mathrm{off}}}\hat{C}_1+\hat{r}' \qquad \hat{C}_2\xrightarrow{k_r^1}\hat{C}_1' \\
&\hat{C}_1'+\hat{r}\xrightarrow{\kappa^{\mathrm{on}\prime}}\hat{C}_2 \qquad \hat{C}_2\xrightarrow{\kappa^{\mathrm{off}\prime}}\hat{C}_1'+\hat{r} \qquad \hat{C}_2\xrightarrow{k_r^{1\prime}}\hat{C}_1 \qquad C_2\xrightarrow{k_R^2}2r.
\end{aligned}
$$

The nondimensionalized ODEs of this model are

$$
\begin{aligned}
dR/dt &= \sigma_R - \kappa^{\mathrm{on}}Rr - \kappa^{\mathrm{on}\prime}Rr' + \kappa^{\mathrm{off}}C_1 + \kappa^{\mathrm{off}\prime}C_1' - R + \beta_1\gamma C_1 + \beta_1'\gamma' C_1' \\
dr/dt &= 1 - \kappa^{\mathrm{on}}Rr + \kappa^{\mathrm{off}}C_1 - \kappa^{\mathrm{on}}C_1'r + \kappa^{\mathrm{off}}C_2 - \gamma r + \alpha_1 C_1 + \alpha_2 C_2 \\
dr'/dt &= \sigma_r' - \kappa^{\mathrm{on}\prime}Rr' + \kappa^{\mathrm{off}\prime}C_1' - \kappa^{\mathrm{on}\prime}C_1 r' + \kappa^{\mathrm{off}\prime}C_2 - \gamma' r' + \alpha_1' C_1' + \alpha_2 C_2 \\
dC_1/dt &= \kappa^{\mathrm{on}}Rr - \kappa^{\mathrm{off}}C_1 - \kappa^{\mathrm{on}\prime}C_1 r' + \kappa^{\mathrm{off}\prime}C_2 - \alpha_1 C_1 - \beta_1\gamma C_1 + \beta_2'\gamma' C_2 \qquad (40) \\
dC_1'/dt &= \kappa^{\mathrm{on}\prime}Rr' - \kappa^{\mathrm{off}\prime}C_1' - \kappa^{\mathrm{on}}C_1'r + \kappa^{\mathrm{off}}C_2 - \alpha_1' C_1' - \beta_1'\gamma' C_1' + \beta_2\gamma C_2 \\
dC_2/dt &= \kappa^{\mathrm{on}}C_1'r + \kappa^{\mathrm{on}\prime}C_1 r' - \kappa^{\mathrm{off}}C_2 - \kappa^{\mathrm{off}\prime}C_2 - 2\alpha_2 C_2 - \beta_2\gamma C_2 - \beta_2'\gamma' C_2.
\end{aligned}
$$

In addition to the variables and parameters defined for the MMI2-ASB Model, $r'$ represents (the concentration of) the second microRNA that binds to the second side on the mRNA ($R$), and $C_1'$ represents (the concentration of) the 1:1 complex formed by the second microRNA and the mRNA. $\kappa^{\mathrm{on}\prime}$ and $\kappa^{\mathrm{off}\prime}$ represent the association and dissociation rate constants of the second binding site respectively. $\alpha_1'$, $\beta_1'$ and $\beta_2'$ are RDFs. $\alpha_1'$ represents the degradation rate constant of the mRNA in the 1:1 complex $C_1'$ relative to that of the unbound mRNA. $\beta_1'$ and $\beta_2'$ represent the degradation rate constants of the second microRNA in the 1:1 and 2:1 complexes, respectively, relative to its degradation rate constant in the unbound form. $\gamma'$ represents the degradation rate constant of the unbound form of the second microRNA relative to that of the unbound mRNA. $\sigma_r'$ represents the synthesis rate constant of the second microRNA relative to that of the first microRNA.

These equations are used to perform numerical simulations and bifurcation analysis for the MMI2-DMI Model. We found that it is not feasible to use the tQSSA to reduce this model to a 2-ODE system.

Applying the CRNT to the network showed that the MMI2-ASB Model can have two stable steady states. Consistent with this result, numerical bifurcation analysis with random parameter values using the same scheme described earlier produced parameter sets that allow the MMI2-ASB to be bistable.

### 3. The structurally perturbed MMI2 Models

#### *3.1 The C2-Knockout (C2KO) Model*

To investigate whether the 2:1 complex is required to achieve oscillation and bistability, we removed the complex variable $C_2$ from the MMI2-ASB and the MMI2-SSB Models, both of which can be considered minimum MMI2 Models. Removing $C_2$ (or $\hat{C}_2$) from the MMI2-ASB Model gives rise to a reaction network identical to the MMI1 Model, which is analyzed in Section 1. We next considered a perturbed version of the MMI2-SSB Model

$$\xrightarrow{s_R} \hat{R} \qquad \hat{R} \xrightarrow{k_R^0} \emptyset \qquad \xrightarrow{s_r} \hat{r} \qquad \hat{r} \xrightarrow{k_r^0} \emptyset \qquad \hat{R} + \hat{r} \xrightarrow{k^{\text{on}}} \hat{C}_1 \qquad \hat{C}_1 \xrightarrow{k^{\text{off}}} \hat{R} + \hat{r} \qquad \hat{C}_1 \xrightarrow{k_r^1} \hat{R} \qquad \hat{C}_1 \xrightarrow{k_R^1} \hat{r}$$

$$\hat{R} + \hat{r} \xrightarrow{k^{\text{on}\prime}} \hat{C}_1' \qquad \hat{C}_1' \xrightarrow{k^{\text{off}\prime}} \hat{R} + \hat{r} \qquad \hat{C}_1' \xrightarrow{k_r^{1\prime}} \hat{R} \qquad \hat{C}_1' \xrightarrow{k_R^{1\prime}} \hat{r}. \tag{41}$$

In addition to the structural perturbation of removing $C_2$ (or $\hat{C}_2$), we also relaxed the assumption that the two binding sites are identical. This consideration of a more general case allows us to exclude the possibility that the two distinct binding sites may generate bistability or oscillation. Here, $\hat{C}_1'$ represents the 1:1 complex with the second site bound to microRNA and the concentration of the complex. $k^{\text{on}\prime}$ and $k^{\text{off}\prime}$ are the association rate constant and the dissociation rate constant for the second binding site. $k_R^{1\prime}$ and $k_r^{1\prime}$ are the degradation rate constants of the mRNA and microRNA, respectively, in the complex $\hat{C}_1'$. The definitions of other variables and rate constants are identical to those in the MMI2-SSB Model.

The nondimensionalized ODEs of the C2KO Model are

$$\begin{aligned}
dR/dt &= \sigma_R - \kappa^{\text{on}} Rr - \kappa^{\text{on}\prime} Rr + \kappa^{\text{off}} C_1 + \kappa^{\text{off}\prime} C_1' - R + \beta_1 \gamma C_1 + \beta_1' \gamma C_1' \\
dr/dt &= 1 - \kappa^{\text{on}} Rr - \kappa^{\text{on}\prime} Rr + \kappa^{\text{off}} C_1 + \kappa^{\text{off}\prime} C_1' - \gamma r + \alpha_1 C_1 + \alpha_1' C_1' \\
dC_1/dt &= \kappa^{\text{on}} Rr - \kappa^{\text{off}} C_1 - \alpha_1 C_1 - \beta_1 \gamma C_1 \\
dC_1'/dt &= \kappa^{\text{on}\prime} Rr - \kappa^{\text{off}\prime} C_1' - \alpha_1' C_1' - \beta_1' \gamma C_1'.
\end{aligned} \tag{42}$$

Application of the CRNT to this network gave rise to the conclusion that the C2KO Model cannot admit two stable positive steady states, and we obtained a consistent result with numerical bifurcation analysis with random parameter sets.

We apply the Routh-Hurwitz criterion to the ODE system in Eq 42 to prove that any positive steady state (if one exists) is locally stable for any choice of positive rate constants. For a brief discussion of the Routh-Hurwitz criterion, see Section 1.4 or (Bodson, 2020; Meinsma, 1995).

**Theorem (Routh-Hurwitz stability criterion for degree 4 polynomials):** Consider the polynomial $p(\lambda) = a_4\lambda^4 + a_3\lambda^3 + \cdots + a_0$ with $a_0 \neq 0$ and $a_4 > 0$. Build the table with five rows:

$$\begin{array}{ccc}
a_4 & a_2 & a_0 \\
a_3 & a_1 & 0 \\
\tilde{u} & a_0 & 0 \\
\tilde{v} & 0 & 0 \\
a_0 & 0 & 0
\end{array}$$

where $\tilde{u} = \frac{a_2 a_3 - a_4 a_1}{a_3}$ and $\tilde{v} = \frac{u a_1 - a_3 a_0}{u}$. All the roots of $p(\lambda)$ have negative real parts if and only if $a_3, a_0 > 0$, $\tilde{u} > 0$, and $\tilde{v} > 0$.

We may replace the last two inequalities with the following: $u := a_2 a_3 - a_4 a_1 > 0$ and $v := \tilde{u} a_1 - a_3^2 a_0 > 0$. Note that $a_3, a_0 > 0$, $\tilde{u} > 0$ and $\tilde{v} > 0$ if and only if $a_3, a_0 > 0$, $u > 0$ and $v > 0$.

The following expressions are obtained using Mathematica (C2KO.nb). The ODE system in Eq 42 has the Jacobian matrix

$$J = \begin{bmatrix} -1 - \kappa^{\text{on}} r - \kappa^{\text{on}'} r & -\kappa^{\text{on}} R - \kappa^{\text{on}'} R & \kappa^{\text{off}} + \beta_1 \gamma & \kappa^{\text{off}'} + \beta_1' \gamma \\ -\kappa^{\text{on}} r - \kappa^{\text{on}'} r & -\gamma - \kappa^{\text{on}} R - \kappa^{\text{on}'} R & \kappa^{\text{off}} + \alpha_1 & \kappa^{\text{off}'} + \alpha_1' \\ \kappa^{\text{on}} r & \kappa^{\text{on}} R & -\kappa^{\text{off}} - \alpha_1 - \beta_1 \gamma & 0 \\ \kappa^{\text{on}'} r & \kappa^{\text{on}'} R & 0 & -\kappa^{\text{off}'} - \alpha_1' - \beta_1' \gamma \end{bmatrix}. \quad (43)$$

The characteristic polynomial $p(\lambda) = \det(J - \lambda)$ is a degree 4 polynomial, with leading coefficient $a_4 = 1$. Symbolic computation using Mathematica reveals that

$$a_3 = 1 + \kappa^{\text{on}}(R + r) + \kappa^{\text{on}'}(R + r) + \kappa^{\text{off}} + \kappa^{\text{off}'} + \alpha_1 + \alpha_1' + \beta_1 \gamma + \beta_1' \gamma + \gamma, \quad (44)$$

which is clearly positive for any positive parameters and positive steady state values.

The term

$$\begin{aligned} a_0 = {} & \kappa^{\text{on}}\left(\alpha_1 \gamma \kappa^{\text{off}\prime} r + \beta_1 \gamma \kappa^{\text{off}\prime} R + \alpha_1 \alpha_1' \gamma r + \alpha_1' \beta_1 \gamma R + \alpha_1 \beta_1' \gamma^2 r + \beta_1 \beta_1' \gamma^2 R\right) \\ & + \kappa^{\text{on}\prime}\left(\alpha_1' \gamma \kappa^{\text{off}} r + \beta_1' \gamma \kappa^{\text{off}} R + \alpha_1 \alpha_1' \gamma r + \alpha_1 \beta_1' \gamma R + \alpha_1' \beta_1 \gamma^2 r + \beta_1 \beta_1' \gamma^2 R\right) + \gamma \kappa^{\text{off}} \kappa^{\text{off}\prime} \\ & + \kappa^{\text{off}\prime}(\beta_1 \gamma^2 + \alpha_1 \gamma) + \kappa^{\text{off}}(\beta_1' \gamma^2 + \alpha_1' \gamma) + \alpha_1 \alpha_1' \gamma + \alpha_1' \beta_1 \gamma^2 + \alpha_1 \beta_1' \gamma^2 \\ & + \beta_1 \beta_1' \gamma^3 \end{aligned} \quad (45)$$

is also always positive. Note that the expression for $a_0$ is starting to be unwieldy; the last two expressions that we must evaluate are even more so. The coefficients in $a_0$ are extracted and presented in the bar chart in Fig. S15A. The only coefficient value is 1, as we observe in the expression for $a_0$.

Next, we computed $u$, with 282 terms and whose coefficients are presented in the bar chart in Fig. S15B. The coefficients range from 1 to 3, hence $u > 0$.

Finally, $v$, with 5091 terms, has coefficients ranging from 1 to 20; thus $v > 0$. The distribution of coefficients is presented in Fig. S15C. By the Routh-Hurwitz criterion, we conclude that if a positive steady state exists, in which case it is the unique positive steady state, it is asymptotically stable.

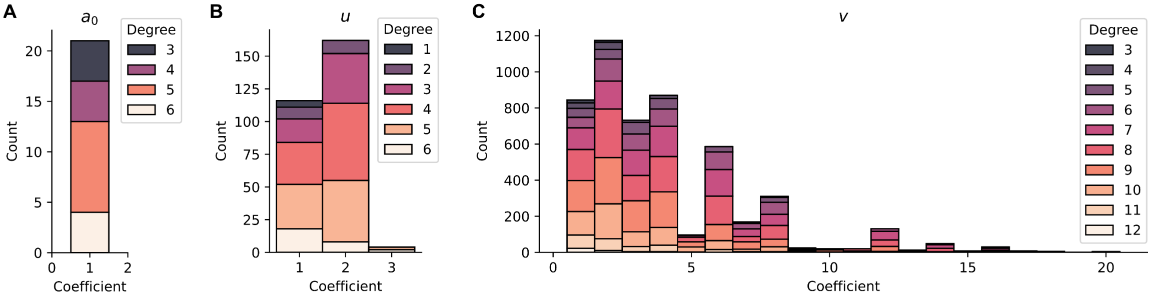

**Fig. S15. Coefficient distributions of $\boldsymbol{a_0}$, $\boldsymbol{u}$ and $\boldsymbol{v}$.** Stacked bar charts show coefficient distributions of the indicated elements. The variables of these polynomials include both state variables and parameters in the C2KO Model.

#### *3.2 The C1-Knockout (C1KO) Model*

We considered a structurally perturbed MMI2-SSB/ASB Model by removing the 1:1 complex(es). The reactions of the C1KO Model are

$$\xrightarrow{s_R} \hat{R} \qquad \hat{R} \xrightarrow{k_R^0} \emptyset \qquad \xrightarrow{s_r} \hat{r} \qquad \hat{r} \xrightarrow{k_r^0} \emptyset \qquad \hat{R} + 2\hat{r} \xrightarrow{k^{\text{on}}} \hat{C}_2 \qquad \hat{C}_2 \xrightarrow{k^{\text{off}}} \hat{R} + 2\hat{r}$$

$$\hat{C}_2 \xrightarrow{2k_r^2} \hat{r} + \hat{R} \qquad \hat{C}_2 \xrightarrow{k_R^2} 2\hat{r}. \tag{46}$$

In this model, two microRNA molecules bind to the two binding sites on the mRNA simultaneously in one reaction to form a 2:1 complex. This can be viewed as an extreme form of positive binding cooperativity. The degradation of any microRNA molecule in the complex leads to the dissociation of the mRNA and the remaining microRNA. The nondimensionalized ODEs of the model are

$$\begin{aligned} dR/dt &= \sigma_R - \kappa^{\text{on}} R r^2 + \kappa^{\text{off}} C_2 - R + 2\beta_2 \gamma C_2 \\ dr/dt &= 1 - 2\kappa^{\text{on}} R r^2 + 2\kappa^{\text{off}} C_2 - \gamma r + 2\beta_2 \gamma C_2 + 2\alpha_2 C_2 \\ dC_2/dt &= \kappa^{\text{on}} R r^2 - \kappa^{\text{off}} C_2 - \alpha_2 C_2 - 2\beta_2 \gamma C_2. \end{aligned} \tag{47}$$

Application of the CRNT to this network gave rise to the conclusion that the C1KO Model cannot admit two stable positive steady states, and we obtained a consistent result with numerical bifurcation analysis with random parameter sets.

Similar to the analytical analysis in the C2KO Model, we prove that any positive steady state of the ODE system in Eq 47, if it exists, is locally stable for any choice of positive rate constants. In particular, we apply the Routh-Hurwitz criterion for degree 3 polynomials (see Section 1.4).

The following expressions are obtained using Mathematica (C1KO.nb). The system has the Jacobian matrix

$$J = \begin{bmatrix} -1 - \kappa^{\text{on}} r^2 & -2\kappa^{\text{on}} R r & \kappa^{\text{off}} + 2\beta_2 \gamma \\ -2\kappa^{\text{on}} r^2 & -\gamma - 4\kappa^{\text{on}} R r & 2\kappa^{\text{off}} + 2\alpha_2 + 2\beta_2 \gamma \\ \kappa^{\text{on}} r^2 & 2\kappa^{\text{on}} R r & -\kappa^{\text{off}} - \alpha_2 - 2\beta_2 \gamma \end{bmatrix}. \tag{48}$$

The characteristic polynomial $p(\lambda) = \det(\lambda - J)$ is a degree-3 polynomial. We found that $p(\lambda) = \lambda^3 + a_2\lambda^2 + a_1\lambda + a_0$, where

$$a_2 = 1 + \alpha_2 + \gamma + 2\beta_2\gamma + \kappa^{\text{off}} + \kappa^{\text{on}}r^2 + 4\kappa^{\text{on}}Rr,$$

$$a_1 = \alpha_2 + \gamma + \alpha_2\gamma + 2\beta_2\gamma + 2\beta_2\gamma^2 + \kappa^{\text{off}} + \gamma\kappa^{\text{off}} + 4\kappa^{\text{on}}Rr + \alpha_2\kappa^{\text{on}}r^2 + \gamma\kappa^{\text{on}}r^2 + 4\beta_2\gamma\kappa^{\text{on}}Rr,$$

$$a_0 = \alpha_2\gamma + 2\beta_2\gamma^2 + \gamma\kappa^{\text{off}} + \alpha_2\gamma\kappa^{\text{on}}r^2 + 4\beta_2\gamma\kappa^{\text{on}}Rr. \quad (49)$$

It is clear from the expression that for any positive rate constants, with steady state values for $R$ and $r$, we always have $a_0, a_2 > 0$. Lastly, the minimum non-zero coefficient of $u$ is 1, so $u > 0$.

By the Routh-Hurwitz stability criterion, we conclude that for any positive rate constants, any eigenvalue of the system has negative real part, and the unique positive steady state (if one exists) is asymptotically stable.

#### *3.3 Other structurally perturbed models*

Removing the synthesis of the mRNA or microRNA from the system yields trivial cases. The long-term behavior of the system depends on either

$$dR/dt = \sigma_R - R \quad (50)$$

when the microRNA synthesis is removed, or

$$dr/dt = 1 - \gamma r \quad (51)$$

when the mRNA synthesis is removed. The vector fields of these one-dimensional autonomous systems are flows in a line and there is only one steady state for each system. Therefore, neither of these ODEs can produce spiral sinks, oscillation, or bistability.

#### *3.4 Summary of dynamical features of all MMI models*

| Model | Spiral sink under biologically relevant condition? | Sustained oscillation under biologically relevant condition? | Bistability under biologically relevant condition? |
|---|---|---|---|
| MMI1 | Yes | No | No |
| MMI2-SSB | Yes | Yes | Yes |
| MMI2-ASB | Yes | Yes | Yes |
| MMI2-DMI | Yes | Yes | Yes |
| C2KO | Yes | No | No |
| C1KO | Yes | No | No |

**Table S3.** Summary of dynamical features of all MMI models.

### 4. Comparison to repressilator

#### *4.1 Repressilator model*

To determine whether the rapidly diverging periods of the MMI2-SSB-produced limit cycles may contribute to the restoration of heterogeneity, we sought to compare the MMI2-SSB Models to an oscillator with a consistent, nondiverging period. We selected the repressilator (Elowitz and Leibler, 2000) as a well-characterized nondiverging oscillator (Fig. S16A). Its transcriptional and translational dynamics can be modeled with the dimensionless six-ODE system

$$\begin{aligned} dR_1/dt &= \alpha_0 + \frac{\alpha_1}{1+P_3^n} - k_R R_1 \\ dR_2/dt &= \alpha_0 + \frac{\alpha_2}{1+P_1^n} - k_R R_2 \\ dR_3/dt &= \alpha_0 + \frac{\alpha_3}{1+P_2^n} - k_R R_3 \\ dP_1/dt &= \beta R_1 - k_P P_1 \\ dP_2/dt &= \beta R_2 - k_P P_2 \\ dP_3/dt &= \beta R_3 - k_P P_3 \end{aligned} \tag{52}$$

where gene $i$'s mRNA concentration is $R_i$ and its protein concentration is $P_i$. The parameters, selected to produce oscillations of comparable period and amplitude to MMI2-SSB oscillations, are given in Table S4. The tested range of signal $\alpha_1$ was selected to cover the range in which the system exhibits limit cycle oscillations.

| Parameter | Description | Value |
|---|---|---|
| $\alpha_0$ | Basal transcription rate for all genes | 0.01 |
| $\alpha_1$ | Regulated transcription rate for gene 1 (signal) | 3 to 23 (steps of 1) |
| $\alpha_2$ | Regulated transcription rate for gene 2 | 5 |
| $\alpha_3$ | Regulated transcription rate for gene 3 | 5 |
| $n$ | Cooperativity of transcriptional repression | 2 |
| $\beta$ | Translation rate for all mRNAs | 1 |
| $k_R$ | Degradation rate for all mRNAs | 1 |
| $k_P$ | Degradation rate for all proteins | 1 |

**Table S4.** Parameters in the repressilator model.

#### *4.2 Heterogeneity restoration performance of the repressilator*

The repressilator model was simulated under both additive and multiplicative noise as in Section 2.1.5 with the following exceptions. 100 initial conditions were selected from a 3-dimensional Sobol sequence (Bratley and Fox, 1988) using Sobol.jl, scaled to $[0, 5]$, and perturbed by $+0.01$ in the first component to avoid unstable states. Because mRNA and protein species reach similar magnitudes in the repressilator model, each 3-long vector was used as the initial conditions for both the three mRNA concentrations and the three protein concentrations. Because the repressilator can exhibit very slowly damping oscillations, each deterministic simulation proceeded for 5000 time units, extreme initial conditions were selected after 4000 time units based on the value of $R_1$, and stochastic simulations proceeded for 1000 time units. Both mRNA and protein species were subject to the same noise level as the mRNA in the MMI2-SSB tests.

The simulation endpoints were analyzed for bimodality as in Section 2.1.7. While the oscillations caused clear variability in gene expression, the bimodality expected of distinguishable subpopulations was inconsistent or subtle at best (Fig. S16B-C). Even when the simulation endpoint met the mode separation cutoff of $D > 2$, the population exhibited extreme, long-lasting oscillations in gene expression (Fig. S16D and G) that are not consistent with observed restoration of heterogeneity in hours to a few days following subtle oscillation (Chang et al., 2008). We therefore propose that the "diverging" nature of oscillators produced by the MMI2-SSB system may contribute to observed restoration of heterogeneity restoration.

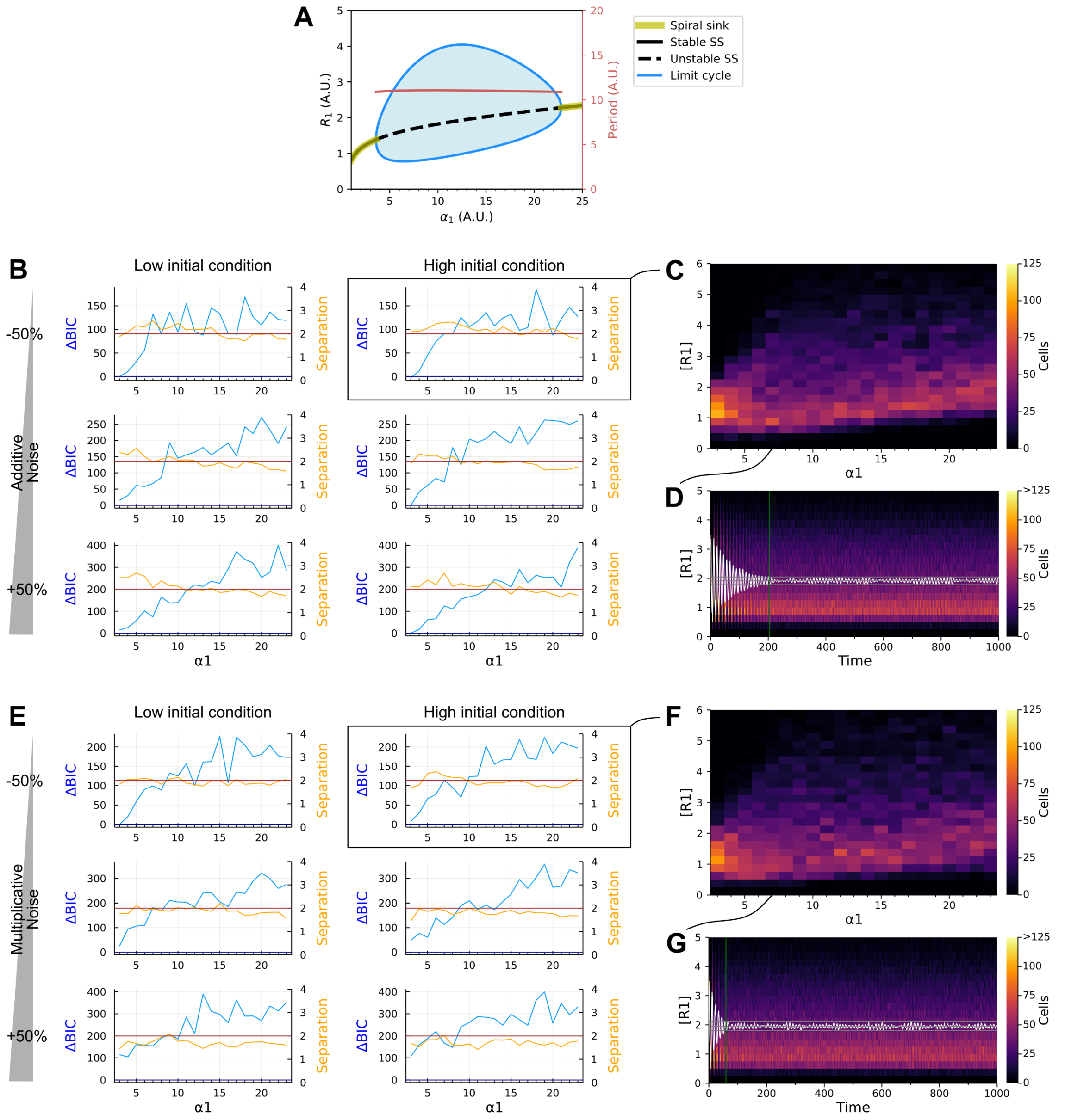

**Fig. S16. Heterogeneity restoration performance of the repressilator, a nondiverging oscillator.** Parameters are given in Table S5. (**A**) Bifurcation diagram with respect to regulated transcription rate $\alpha_1$, showing the stable limit cycle. Its period does not change significantly with respect to the signal. SS, steady state. (**B**) Heterogeneity restoration performance under additive noise. The populations' distributions of RNA 1 are nonnormal (high ΔBIC), but inconsistently and marginally bimodal (low separation, $D$). (**C**) Final distributions of RNA 1 concentration starting from the high initial condition at low noise. (**D**) RNA 1 concentration distribution over the timecourse starting from the high initial condition at low noise with $\alpha_1 = 7$. The population's bimodality appears poor. White line, population mean; gray lines, "late" maximum, mean, and minimum mean after time 400; green line, equilibrium time. (**E-G**) As B-D for multiplicative noise.

### References

Bodson, M. (2020). Explaining the Routh–Hurwitz Criterion: A Tutorial Presentation [Focus on Education]. IEEE Control Systems Magazine *40*, 45-51.

Bratley, P., and Fox, B.L. (1988). Algorithm 659: Implementing Sobol's quasirandom sequence generator. ACM Trans. Math. Softw. *14*, 88–100. 10.1145/42288.214372.

Chan, L.Y., Mugler, C.F., Heinrich, S., Vallotton, P., and Weis, K. (2018). Non-invasive measurement of mRNA decay reveals translation initiation as the major determinant of mRNA stability. Elife *7*, e32536.

Chang, H.H., Hemberg, M., Barahona, M., Ingber, D.E., and Huang, S. (2008). Transcriptome-wide noise controls lineage choice in mammalian progenitor cells. Nature *453*, 544-547. 10.1038/nature06965.

Choi, K., Medley, J.K., König, M., Stocking, K., Smith, L., Gu, S., and Sauro, H.M. (2018). Tellurium: An extensible python-based modeling environment for systems and synthetic biology. Biosystems *171*, 74-79.

Chubb, J.R., Trcek, T., Shenoy, S.M., and Singer, R.H. (2006). Transcriptional pulsing of a developmental gene. Curr. Biol. *16*, 1018-1025.

Coomer, M.A., Ham, L., and Stumpf, M.P.H. (2021). Noise distorts the epigenetic landscape and shapes cell-fate decisions. Cell Systems.

de la Mata, M., Gaidatzis, D., Vitanescu, M., Stadler, M.B., Wentzel, C., Scheiffele, P., Filipowicz, W., and Großhans, H. (2015). Potent degradation of neuronal miRNAs induced by highly complementary targets. EMBO Rep. *16*, 500-511.

Eichhorn, S.W., Guo, H., McGeary, S.E., Rodriguez-Mias, R.A., Shin, C., Baek, D., Hsu, S.-h., Ghoshal, K., Villén, J., and Bartel, D.P. (2014). mRNA destabilization is the dominant effect of mammalian microRNAs by the time substantial repression ensues. Mol. Cell *56*, 104-115.

Elowitz, M.B., and Leibler, S. (2000). A synthetic oscillatory network of transcriptional regulators. Nature *403*, 335-338. 10.1038/35002125.

Feinberg, M. (1988). Chemical reaction network structure and the stability of complex isothermal reactors—II. Multiple steady states for networks of deficiency one. Chem. Eng. Sci. *43*, 1-25.

Gantmacher, F.R., and Brenner, J.L. (2005). Applications of the Theory of Matrices (Courier Corporation).

Gillespie, D.T. (1977). Exact stochastic simulation of coupled chemical reactions. The Journal of Physical Chemistry *81*, 2340-2361. 10.1021/j100540a008.

Gingras, J., Rioux, R.M., Cuvelier, D., Geisse, N.A., Lichtman, J.W., Whitesides, G.M., Mahadevan, L., and Sanes, J.R. (2009). Controlling the orientation and synaptic differentiation of myotubes with micropatterned substrates. Biophys. J. *97*, 2771-2779.

Jarmoskaite, I., AlSadhan, I., Vaidyanathan, P.P., and Herschlag, D. (2020). How to measure and evaluate binding affinities. Elife *9*, e57264.

Lan, Y., Elston, T.C., and Papoian, G.A. (2008). Elimination of fast variables in chemical Langevin equations. The Journal of chemical physics *129*, 12B607.

Lecca, P. (2013). Stochastic chemical kinetics : A review of the modelling and simulation approaches. Biophys Rev *5*, 323-345. 10.1007/s12551-013-0122-2.

Marzi, M.J., Ghini, F., Cerruti, B., De Pretis, S., Bonetti, P., Giacomelli, C., Gorski, M.M., Kress, T., Pelizzola, M., and Muller, H. (2016). Degradation dynamics of microRNAs revealed by a novel pulse-chase approach. Genome Res. *26*, 554-565.

Meinsma, G. (1995). Elementary proof of the Routh-Hurwitz test. Systems & Control Letters *25*, 237-242.

Moore, M.J., Sebastian, J.A., and Kolios, M.C. (2019). Determination of cell nucleus-to-cytoplasmic ratio using imaging flow cytometry and a combined ultrasound and photoacoustic technique: a comparison study. J. Biomed. Opt. *24*, 106502.

Muratov, A.L., and Gnedin, O.Y. (2010). Modeling the Metallicity Distribution of Globular Clusters. The Astrophysical Journal *718*, 1266-1288. 10.1088/0004-637x/718/2/1266.

Pedregosa, F., Varoquaux, G., Gramfort, A., Michel, V., Thirion, B., Grisel, O., Blondel, M., Prettenhofer, P., Weiss, R., Dubourg, V., et al. (2011). Scikit-learn: Machine Learning in Python. Journal of Machine Learning Research *12*, 2825-2830.

Pinzón, N., Li, B., Martinez, L., Sergeeva, A., Presumey, J., Apparailly, F., and Seitz, H. (2017). microRNA target prediction programs predict many false positives. Genome Res. *27*, 234-245.

Rackauckas, C., and Nie, Q. (2017). Differentialequations. jl–a performant and feature-rich ecosystem for solving differential equations in julia. Journal of Open Research Software *5*.

Reichholf, B., Herzog, V.A., Fasching, N., Manzenreither, R.A., Sowemimo, I., and Ameres, S.L. (2019). Time-resolved small RNA sequencing unravels the molecular principles of microRNA homeostasis. Mol. Cell *75*, 756-768.

Sharova, L.V., Sharov, A.A., Nedorezov, T., Piao, Y., Shaik, N., and Ko, M.S.H. (2009). Database for mRNA half-life of 19 977 genes obtained by DNA microarray analysis of pluripotent and differentiating mouse embryonic stem cells. DNA Res. *16*, 45-58.

Vasudevan, S., Tong, Y., and Steitz, J.A. (2007). Switching from repression to activation: microRNAs can up-regulate translation. Science *318*, 1931-1934.

Wee, L.M., Flores-Jasso, C.F., Salomon, W.E., and Zamore, P.D. (2012). Argonaute divides its RNA guide into domains with distinct functions and RNA-binding properties. Cell *151*, 1055-1067.

Yang, E., van Nimwegen, E., Zavolan, M., Rajewsky, N., Schroeder, M., Magnasco, M., and Darnell, J.E. (2003). Decay rates of human mRNAs: correlation with functional characteristics and sequence attributes. Genome Res. *13*, 1863-1872.

Zlotorynski, E. (2019). Insights into the kinetics of microRNA biogenesis and turnover. Nature Reviews Molecular Cell Biology *20*, 511-511.
